# The proteasome regulator PSME4 drives immune evasion and abrogates anti-tumor immunity in NSCLC

**DOI:** 10.1101/2021.10.24.464690

**Authors:** Aaron Javitt, Merav D. Shmueli, Matthias P Kramer, Aleksandra A. Kolodziejczyk, Ivan J. Cohen, Iris Kamer, Kevin Litchfield, Elizabeta Bab-Dinitz, Oranit Zadok, Vanessa Neiens, Adi Ulman, Lihi Radomir, Hila Wolf-Levy, Avital Eisenberg-Lerner, Assaf Kacen, Michal Alon, Ana Toste Rêgo, Elvira Stacher-Priehse, Michael Lindner, Ina Koch, Jair Bar, Charles Swanton, Yardena Samuels, Yishai Levin, Paula C. A. da Fonseca, Eran Elinav, Nir Friedman, Silke Meiners, Yifat Merbl

## Abstract

Protein degradation by proteasomes is important for the immune response against tumors. Antigens generated by the proteasome promote immune cell infiltration into tumors and improve tumors’ responses to immunotherapy. For example, immunoproteasomes – a subset of proteasomes induced by inflammatory signals – may improve the response of melanomas to immune checkpoint inhibitors (ICI) by eliciting tumor inflammation. Yet, it is unclear whether and how protein degradation by proteasomes impacts cancer progression and contributes to immune evasion and resistance. Here, we profile the proteasome-cleaved peptides in lung cancers and find that PSME4 serves as a novel inhibitory regulator of the immunoproteasome, playing an anti-inflammatory role in cancer. Biochemical assays combined with scRNA-seq, immunopeptidomics and in vivo analyses demonstrate that PSME4 promotes an immunosuppressive environment around the tumor and abrogates anti-tumor immunity by inhibiting antigen presentation and attenuating tumor inflammation. Furthermore, we find that PSME4 expression is correlated with responsiveness to ICI across several cancer types. Our findings suggest that PSME4-mediated regulation of proteasome activity is a novel mechanism of immune evasion in non-small-cell lung carcinoma and may be targeted therapeutically for restoring anti-tumor immunity.

**Graphical Abstract:** 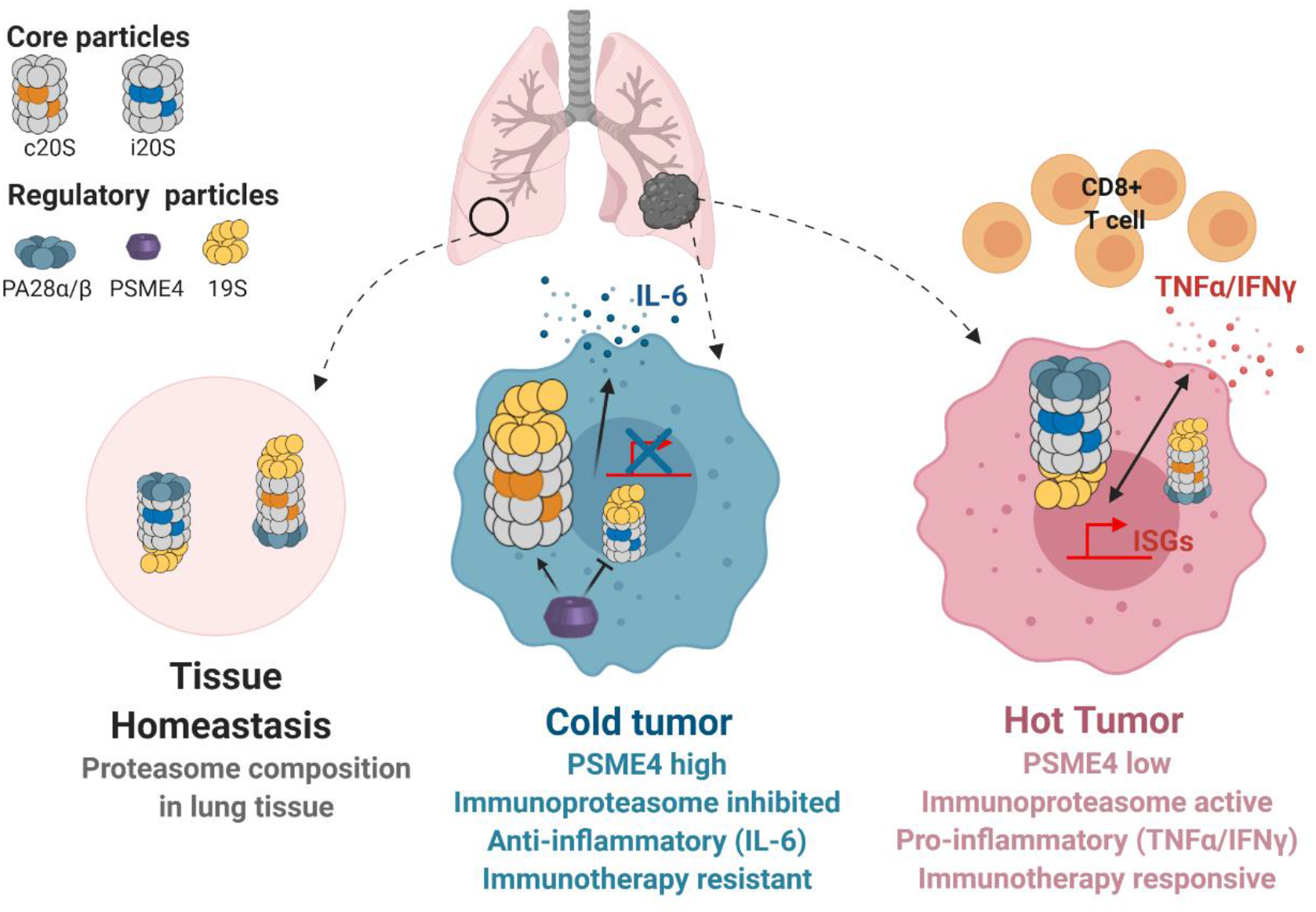

**Highlights:**

- Mapping the degradation landscape in Non-Small Cell Lung Cancer (NSCLC) uncovers altered proteasome activity and composition
- Proteasome regulator PSME4 plays an anti-inflammatory role in NSCLC by attenuating immunoproteasome activity
- PSME4 restricts tumor antigen presentation and cytokine secretion, defining a ‘cold’ tumor environment
- PSME4 drives tumor immune evasion and is associated with resistance to immunotherapy

## Introduction

Proteasomal degradation of proteins plays a key role in inflammation and antigen presentation. Further, immunoproteasome upregulation has been associated with response to immune checkpoint inhibitors (ICI; (Ayers et al., 2017; Harel et al., 2019; Kalaora et al., 2020; Riaz et al., 2017a; Rousseau and Bertolotti, 2018; Spits and Neefjes, 2016; Tripathi et al., 2016)), yet whether and how proteasomes may play a role in resistance to therapy remains poorly understood. Tumor immunogenicity is promoted by the switch from constitutive proteasomes to immunoproteasomes that are induced by inflammatory signals (Driscoll et al., 1993; Gaczynska et al., 1993; Salzmann et al., 1999; Winter et al., 2017). The canonical constitutive- and immunoproteasomes differ by three catalytic subunits Psmb8, Psmb9 and Psmb10 that are encoding the proteins LMP7, LMP2 and MECL1, respectively. The immunoproteasome is induced by inflammatory cytokines (e.g. IFN) and is basally expressed mainly in immune cells (Driscoll et al., 1993; Gaczynska et al., 1993; Salzmann et al., 1999). Previous studies uncovered an altered cleavage pattern by the immunoproteasome, which is responsible for generating more hydrophobic peptides, that are thought to be preferential for binding the Transporter associated with antigen processing (TAP) and MHC class I presentation (Chong et al., 2018; Javitt et al., 2019a; Rock et al., 2016). Beyond antigenicity, immunoproteasomes have also been suggested to increase cellular inflammation by attenuating IL-6 secretion and promoting inflammatory cytokine secretion (Arima et al., 2011; Ferrington and Gregerson, 2012; Kammerl and Meiners, 2016; Kitamura et al., 2011; Liu et al., 2012; Muchamuel et al., 2009; Schmidt et al., 2018). However, whether and how tumorigenic mechanisms modulate the immunoproteasome and constitutive proteasome functions to shape their microenvironments remains poorly understood.

In addition to the core particle of the proteasome, proteasome composition and the degradation landscape may be affected by specific regulatory particles such as the 19S regulatory complex (PSMC1-6 and PSMD1-14), or the regulatory caps PA28αβ (PSME1-2), PA28γ (PSME3) or PA200 (PSME4) (Collins and Goldberg, 2017; Coux et al., 2020; Rousseau and Bertolotti, 2018) that bind the catalytic core and affect gate opening and substrate selection. The 19S regulatory complex been shown to be pivotal for binding of ubiquitinated species and substrate unfolding, while the other regulatory subunits are suggested to have more specialized roles, including mediating inflammatory responses, histone degradation, and the response to DNA damage (Antoniou et al., 2012; Blickwedehl et al., 2012; Collins and Goldberg, 2017; Khor et al., 2006; Mandemaker et al., 2018a, 2018b; Qian et al., 2013; Rousseau and Bertolotti, 2018; Ustrell et al., 2002; Welk et al., 2016). The different catalytic cores, together with the diverse range of regulatory subunits, introduce great potential diversity for generating hybrid proteasomes of the 20S core particle with different combinations of regulatory caps, thereby altering protein cleavage (Fabre et al., 2014, 2015; Morozov and Karpov, 2019; Raule et al., 2014; Toste Rêgo and da Fonseca, 2019). However, it remains underexplored how the regulatory and catalytic subunits shape tumor-host interactions.

Here, we analyzed, for the first time the degradation landscape in clinical samples of resected tumors and found a distinct signature that was associated with NSCLC. Altered proteasome activity in the cancerous tissue revealed an altered composition of proteasomes, with significant upregulation of the regulatory cap, PSME4, which was associated with poor survival. Exacting functional and biochemical assays together with single cell RNA analysis and in vivo work surprisingly revealed that, PSME4-capped immunoproteasomes, which were not described before, directly attenuate the catalytic activity of the immunoproteasome. We could further show that PSME4-capped constitutive proteasomes exhibited an increase in caspase-like activity in NSCLC. These marked opposing effects upon binding of PSME4 to the constitutive or immuno-proteasomes, in turn, lead to decreased tumor immunogenicity and contribute to tumor growth *in vivo*. Further, we show that PSME4-associated proteasomes play an anti-inflammatory role in NSCLC, and that PSME4 promotes a ‘cold’ tumor signature associated with resistance to ICI across cancer types. Collectively, our findings uncover new insight into the degradation landscape of NSCLC and elucidate a causal role of PSME4 in shaping the tumor microenvironment to suppress anti-tumor immunity. Together, our work introduces a novel paradigm by which proteasome composition and heterogeneity should be examined in the context of cancer and response to immunotherapy in NSCLC and beyond.

## Results

### Proteasome profiling reveals an altered proteasome activity and distinct cleavage patterns in lung adenocarcinoma

To address these questions, we utilized our recently established proteasome profiling approach for Mass spectrometry analysis of proteolytic peptides(Javitt and Merbl, 2019; Wolf-Levy et al., 2018) (MAPP) to analyze tissue samples of non-small cell lung cancer (NSCLC) and adjacent tissue controls from the same lungs, processed in parallel. We then isolated proteasomes from tumor samples and analyzed the nascent peptides that were either ‘trapped’ inside or closely associated with the proteasomes to detect patterns of degradation in single peptide resolution (Figure 1A). Proteomic analysis of both protein abundance and degradation revealed that different tumors had shared patterns of degradation that were distinguished from those of the adjacent control tissues (Figures S1A-D). Detailed examination of the substrates highlighted cellular targets that were readily detected in tumor tissues, some of which (e.g. CASC5; Figures 1B) are associated with cancer and are not expected to be expressed in the healthy lung. Notably, some of the degradation targets were not detected at all by the proteomic analysis, suggesting that they might be rapidly degraded and therefore not accumulating in cells to reach a level that is detectable by mass spectrometry analysis (Figures 1C, S2A-B).

**Fig 1.**
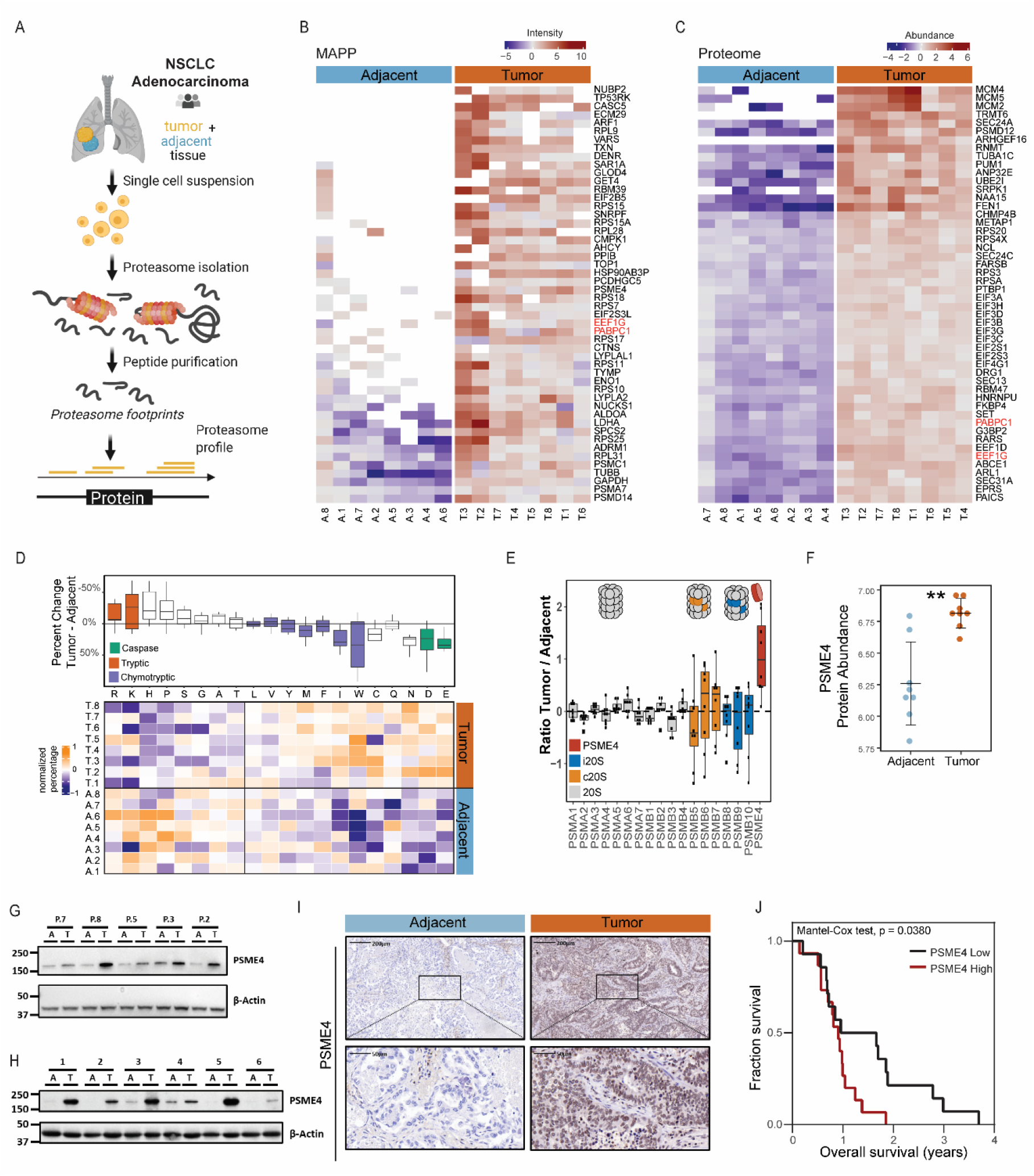
Proteasome profiling reveals an altered proteasome activity and distinct cleavage patterns in lung adenocarcinoma. **(A)** Schematic representation of the workflow. Analysis of lung adenocarcinoma and adjacent lung tissue was performed using standard whole cell extract proteomics to assess protein abundance and MAPP to study the proteasome composition and degradome of the sample (N=8 tumor and 8 adjacent sections). **(B-C)** The top 50 most tumor-associated proteins as identified by MAPP (B) or whole cell Proteomics (C) are shown for each of the tumor and adjacent samples. The proteins in both lists are marked in red. **(D)** MAPP identified peptides were divided based on their carboxy terminal residue. The percentage of peptides with each terminus was calculated for each sample and normalized across patient samples. Amino acids are annotated with their matching proteasome cleavage activity; rows are clustered using Pearson distances. The bars represent the percent change for each amino acid at the carboxy terminus comparing tumors and adjacent tissues. **(E)** The ratio between subunit abundance in the proteasome immunoprecipitated from tumor and adjacent tissue per patient. Colors and diagrams indicate the identity of the proteasome subunits. (**F)** PSME4 abundant in the pulldown from tumor samples and from adjacent tissues (**Wilcoxon P = 0.0078). **(G-H)** Tumor (T) and adjacent (A) tissues from an Israeli (G) or German (H) cohort was immunoblotted for PSME4. β-Actin was used as a loading control. **(I)** Immunohistochemistry of PSME4 in the tumor and adjacent tissues shows increased abundance in the tumor tissue with nuclear localization in both. (**J**) Surviving analysis shows that patients having lung tumors with increased staining of PSME4 (immunohistochemistry) had significantly reduced survival compared to patients with lung tumors expressing low levels of PSME4 (Mantel-Cox test p = 0.0380).

Beyond specific degradation substrates, we found an increase in nuclear proteins that were degraded in the cancerous tissues (Figures S3A-C). Further, we found significant alterations in the cleavage pattern of substrates in the cancerous tissue (Figure 1D). Specifically, the carboxy terminal residue of the degraded peptides, which is the site of proteasome cleavage, differed between the tumor and adjacent samples (Figures. 1D). These observations suggested that proteasome composition and function may be altered in NSCLC. We therefore examined the protein abundance of different proteasome subunits. Proteasome complexes are comprised of the core particle, which process the substrates, and regulatory caps, which control substrate entry(Collins and Goldberg, 2017; Coux et al., 2020; Livneh et al., 2016; Motosugi and Murata, 2019; Rodriguez et al., 2007; Rousseau and Bertolotti, 2018). Notably, while subunits of the proteasome core particle (e.g. 20S and i20S) were similar between the tumor and adjacent tissues, the regulatory cap PSME4 (also known as PA200) was significantly upregulated in the cancerous tissue (Figures 1E-F). We confirmed that PSME4 is upregulated in the cancerous tissue using CPTAC proteomics (Gillette et al., 2020) and 2 additional patient cohorts (Figures 1G-H, S4A-C). As previously reported (Khor et al., 2006) we found PSME4 was localized mainly to the nucleus (Figure 1I, S4D-E). Notably, we showed that high PSME4 levels in lung cancer were associated with poor prognosis of NSCLC patients (Mantel-Cox p value = 0.038, Figures 1J and S5A). However, we did not observe any association in this cohort between PSME4 abundance and NSCLC subtype, smoking or tumor stage (Figure S5A-E). Together, these results show that the proteasome composition and proteasome-associated cleavages are changed in NSCLC and suggested that PSME4 may impact tumor growth through modulating proteasome activity.

### PSME4 restricts immunoproteasome activity and reduces immunopeptidome diversity and cellular inflammation

Proteolytic activities of the constitutive proteasome and immunoproteasome are different(Driscoll et al., 1993; Gaczynska et al., 1993; Salzmann et al., 1999; Winter et al., 2017). We therefore assessed whether the PSME4 level was associated with the function of both proteasome subtypes in patient-derived tumor samples. Notably, the caspase- and chymotryptic-like cleavage activities of the proteasome positively correlated, whereas tryptic-like cleavage negatively correlated with the abundance of PSME4 (Figures 2A-B, S6A). This could not be explained by a general shift in the amino acid composition across the degraded substrates (Figure 2C). PSME4 has been previously shown to alter cleavage specificity of the constitutive proteasome(Blickwedehl et al., 2012; Toste Rêgo and da Fonseca, 2019; Ustrell et al., 2002). However, it was not known if and how PSME4 affected the activity of the immunoproteasome (Figure 2D). We found by biochemical examination that PSME4 bound both constitutive proteasome and immunoproteasome in a human lung cancer cell line (Figure 2E, S6B-D), suggesting that PSME4 can affect the activity of the two proteasome subpopulations independently. We then assessed the effect of supplementing recombinant PSME4 on the proteasome activity in vitro using model substrates for defined constitutive cleavage sites. This system allowed us to decouple the effect of PSME4 on the proteasome from potential transcriptional effects that have been reported in other settings(Blickwedehl et al., 2012; Qian et al., 2013). We found that the caspase-like (β1) activity increased (Figure 2F, S7A-D) whereas the tryptic-like (β2) activity decreased upon PSME4 supplementation (Figure 2G and S7E), corroborating our above results from clinical samples (Figures 2A-B). Surprisingly, PSME4 had opposite effects on the constitutive proteasome β1 and its immunoproteasome counterpart β1i subunit, increasing the former and inhibiting the latter (Figures 2H, S7F). In fact, PSME4 inhibited all immunoproteasome-associated activities (β1i, β2i and β5i) under inflammatory stimulation (Figures S7F-H), in which the immunoproteasome is predominant. Thus, PSME4 is the first proteasomal subunit identified that inhibits the immunoproteasome activity.

**Fig 2.**
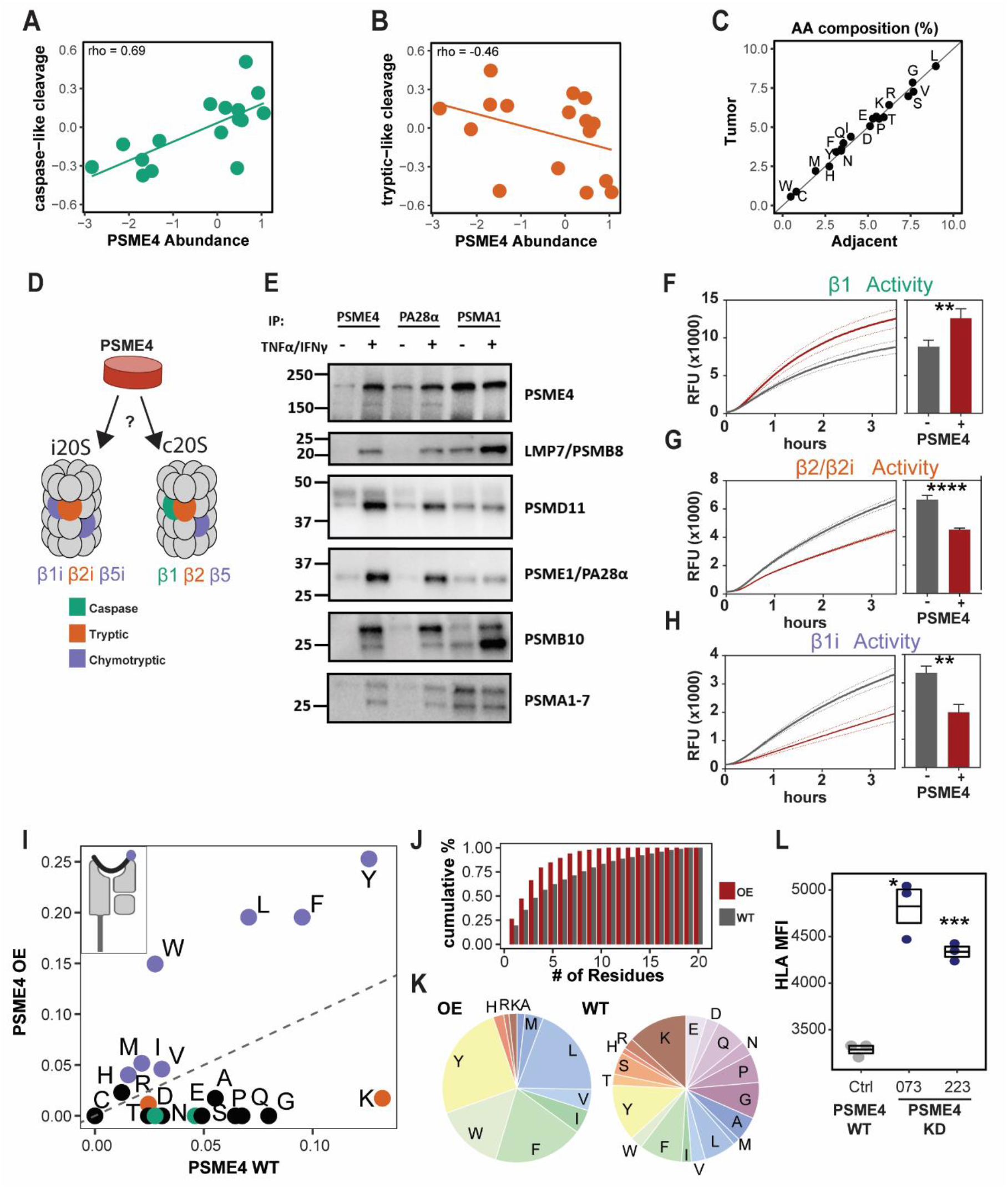
PSME4 restricts proteasome activities and reduces immunopeptidome diversity. **(A – B)** The carboxy terminal residue of peptides was used to classify peptides based on the proteasome activity attributed to their cleavage. The abundance of PSME4 in the samples based on WC proteomics correlated with the caspase-like signature (A; spearman rho = 0.69) and was anti-correlated with the tryptic-like signature (B; spearman rho = -0.46). **(C)** The average amino acid percent composition for all the peptides identified by MAPP in the tumor samples (y-axis) is plotted against the corresponding adjacent samples. (x-axis). **(D)** Schematic representation of the constitutive proteasome (c20S) and immunoproteasome (i20S) with associated catalytic subunits. **(E)** Proteasome complexes were immunoprecipitated (IP) with the indicated antibody from A549 cells that were either treated with TNFα and IFNγ or left untreated and blotted for the indicated proteasome subunits. **(F-H)** Proteasome activity assays using the nLPnLD-AMC (F), RLR-AMC (G) or PAL-AMC (H) fluorogenic substrate to measure the activity of indicated subunits. Relative fluorescence (RFU) of the substrate is shown across the 3.5 hours of the experiment (left) or at the endpoint (right). A549 lysates were untreated (F), to measure the constitutive proteasome activity, or treated with TNFα and IFNγ (G and H), to measure the immunoproteasome activity. Recombinant PSME4 was added to the lysate where indicated (red, paired T-test; **P <0.01, **** P < 0.0001). (**I**) The percentage of each amino acid in the carboxy terminus of the peptides differentially presented on MHC molecules in the A549 cells overexpressing PSME4 (PSME4 OE) vs. empty vector control (PSME4 WT) are shown in a scatter plot. Residues are colored based on the associated cleavage activity: green - caspase, purple – chymotryptic and orange – tryptic; and a dotted line is shown for X=Y. **(J)** The cumulative percentage of peptides explained by the number of different amino acids (from one to all twenty) at the carboxy terminus (end position) in PSME4 WT (WT, grey) and PSME4 OE (OE, red) immunopeptidome. **(K)** The percentage of each amino acid in the carboxy terminus of the peptides differentially presented on MHC molecules in the A549 cells overexpressing PSME4 (PSME4 OE) vs. empty vector control (PSME4 WT). **(L)** Median fluorescence intensity from flow cytometry analysis of MHC-I expression using a pan HLA-A/B/C antibody (W6/32) for A549 cells with PSME4 knockdown (shA223 and shA073) or control (shCtrl). The HLA mean fluorescent intensity (MFI) is significantly higher in the PSME4 knockdown compared to the control (* P = 0.01, *** P = 0.0001).

Proteasome-cleaved peptides serve as the major source of antigens presented by class I MHC. Both the observed PSME4-mediated increase in caspase-like activity and decrease in tryptic-like activity suggested generating peptides that are less favorable for MHC-I presentation. We therefore hypothesized that PSME4 might counteract the generation of antigenic epitopes by the immunoproteasome and decrease tumor antigenicity. To address this question, we performed HLA immunopeptidomics of lung cancer cells with and without overexpressing PSME4 (Figure S8A). We identified 463 peptides (10% of the identified peptides) that were significantly changed upon PSME4 overexpression compared to the control (Figure S8B). As caspase-cleaved peptides do not bind well to the MHC (Javitt et al., 2019a; O’Donnell et al., 2020), they are not highly presented in either condition. Yet, the percentage of peptides with tryptic termini were decreased following PSME4 overexpression (Figure 2I), corresponding to the PSME4-driven change in proteasome activity (Figure 2B). Further, examination of the protein source of the presented peptides, we found an enrichment of nuclear proteins in the immunopeptidome following PSME4 overexpression (Figure S8C). Finally, when examining the distribution of amino acids across the differentially presented peptides, we found that the peptides presented in the PSME4-overexpresing cells were less diverse than those in the control (Figures 2J-K). Notably, only 4 residues in the overexpressing cells, compared to 9 residues in the control, make up 80% of the carboxy-termini of the peptides associated with MHC binding, indicating that PSME4 overexpression significantly restricts the diversity of presented antigens (Figures 2J-K). As PSME4 was previously reported to be incorporated in only a small percentage of proteasome complexes (Fabre et al., 2014; Morozov and Karpov, 2019) we sought to examine whether depleting it would impact significantly the immunopeptidome. In accordance, we could show that depleting PSME4 (Figures S9A-C) significantly increased antigen presentation, as exhibited by 30% increase in surface HLA molecules (Figures 2L and S10). Together these results suggest that PSME4 decreases the diversity of the cellular immunopeptidome, attenuates antigen presentation and immunoproteasome activity.

### PSME4 reduces cellular inflammation and cytokine secretion

Beyond antigenicity, the immunoproteasome has been previously implicated in modulating inflammation by modulating cytokine secretion(Arima et al., 2011; Ferrington and Gregerson, 2012; Kammerl and Meiners, 2016; Kitamura et al., 2011; Liu et al., 2012; Muchamuel et al., 2009; Schmidt et al., 2018). We therefore hypothesized that PSME4 might promote an immunosuppressive environment by direct prohibiting tumor inflammation, in addition to modulating antigenicity. To gain insight into the functional consequences of upregulation of PSME4 in NSCLC, we stratified patient tumors from The Cancer Genome Atlas (TCGA) based on their PSME4 expression levels and associated them with cellular functions (Figure S11A). We found that PSME4-high tumors were enriched with pathways related to cell cycle and DNA replication (Figure S11B), whereas PSME4-low tumors were associated with a number of immune and inflammation-related pathways (Figure 3A). To examine whether there might be a causal relationship between PSME4 and cancer inflammation, Proteomic analysis revealed that IFN signaling pathways were enriched in the PSME4-depleted cells following inflammatory stimulation (Figure 3B, S11C-D).

**Fig. 3.**
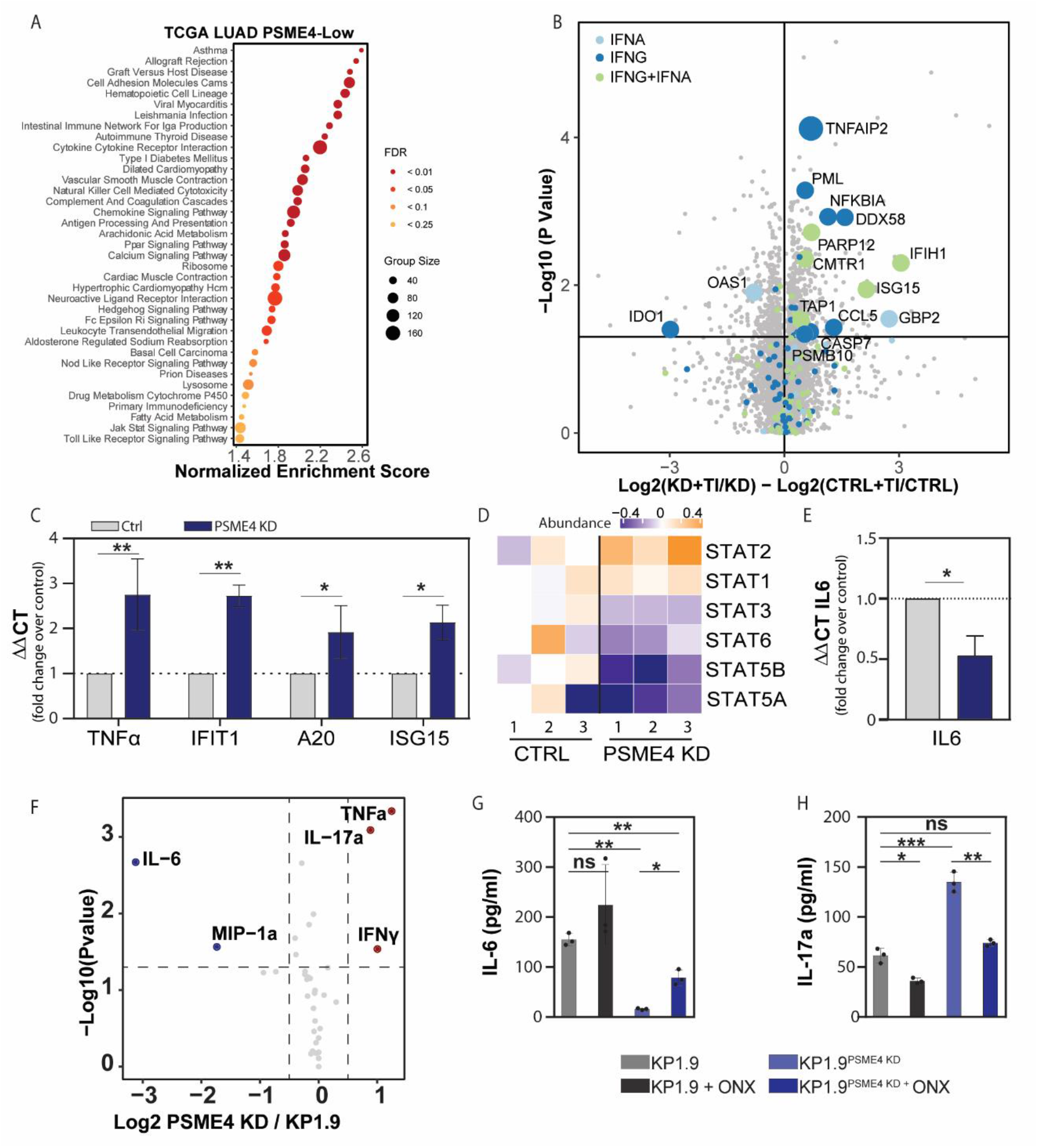
Depleting PSME4 increases cellular inflammation and cytokine secretion. **(A)** TCGA LUAD samples were stratified by PSME4 expression and PSME4-high samples were compared to PSME4-low (n = 230 samples). The normalized enrichment score for KEGG pathways significantly overrepresented in PSME4-low samples are shown. Dot color indicates FDR threshold and size indicates number of genes in the pathway (Group Size). **(B)** The difference of the log2 transformed ratio of protein abundances following stimulation with TNFα and IFNγ (TI) compared to unstimulated (UT) between PSME4 depleted A549 cells (shPSME4) and control (shCtrl) is plotted against the negative log10 transformed p value for a student’s t test comparing the two ratios. Proteins involved in IFN signaling are marked (light blue: response to IFN-alpha (GO: 0035455) dark blue: response to IFN-gamma (GO:0034341), or green: proteins in both lists). Points that significantly differed between the two conditions (student’s t test P ≤ 0.05) and are annotated as IFN signaling pathway components are labeled with the gene name. (**C**) qPCR of genes downstream to IFNγ activation (TNFα, IFIT1, A20 and ISG15) following stimulation with TNFα and IFNγ. A549 cells expressing shPSME4 are compared to those expressing shCtrl (paired T test **P < 0.01 and *P ≤ 0.05). **(D)** Abundance of STAT proteins from shPSME4 -or shCtrl-expressing cells stimulated with TNFα and IFNγ. **(E)** qPCR of IL6 in A549 cells expressing shPSME4 (blue) or shCtrl (grey) following stimulation with TNFα and IFNγ (students’ T test *P ≤ 0.05). **(F)** The log2 transformed ratio of the level of secreted indicated cytokines between PSME4-deficient (KD) and control (KP1.9) KP1.9 cells following stimulation with TNFα and IFNγ is plotted against the negative log10 transformed p value. **(G-H)** The amount of IL-6 (G) or IL-17a (H) secreted from KP1.9 or PSME4-deficient KP1.9 cells following stimulation with TNFα and IFNγ, with or without the immunoproteasome inhibitor ONX-0914 (ONX). Cytokines are measures in pg per ml (student’s T test (ns P>0.05, *P<0.05,**P < 0.01, ***P < 0.001).

Furthermore, we found increased abundance of multiple components of the antigen presentation machinery (e.g. TAP1; Figure 3B) in the PSME4 depleted cells in accordance with the observed increase in surface HLA (Figure 2L). By contrast, KRAS signaling and epithelial to mesenchymal transition (EMT) pathways, which have been previously associated with immunoproteasome-deficient lung cancer (Tripathi et al., 2016) were also enriched here in control conditions with high PSME4 (Figure S11D). Likewise, we found that depleting PSME4 significantly increased the expression of IFN-inducible genes, such as IFIT1 and ISG15 (Figure 3C, S11E-F). Interestingly, STAT1 and STAT2 were more abundant whereas STAT3 and STAT5 were reduced in PSME4-depleted cells than in control cells (Figures 3D). STAT1 and STAT3 have previously been shown to play opposing roles in cancer-related inflammation with STAT3 serving a pro-oncogenic role(Avalle et al., 2012). Consistent with this, we also observed decreased IL6 expression upon PSME4 depletion in two lung cancer cell lines (Figures. 3E, S11D-E). IL-6 is a hallmark of STAT3 signaling and serves an anti-inflammatory and pro-oncogenic role in tumor cells(Avalle et al., 2012; Johnson et al., 2018). Thus, we hypothesized that the upregulation of PSME4 in lung cancer may directly attenuate tumor inflammation and drive immune suppression in the microenvironment by modulating IL-6 expression.

To determine the effect of PSME4 on cytokine secretion in the tumor microenvironment, we utilized the KP1.9 cells derived from a *Kras*^G12D^;*Trp53*^-/-^ murine lung adenocarcinoma in an orthotopic murine model. We established KP1.9 isogenic cell lines with either ectopic expression (KP1.9^PSME4_OE^) or knockdown (KP1.9^PSME4_KD^) of PSME4 (Figures S12A-F). Depletion of PSME4 reduced its levels to be comparable with those of normal lung epithelial cells (Figure S12C). We first examined the cytokine profile of these cell lines and found that TNFα, IFNγ and IL-17a were Increased while IL-6 and MIP-1α were decreased by the depletion of PSME4 (Figure 3F), consistent with the observed changes in the human lung cancer cell line. Importantly, the immunoproteasome has been previously implicated in modulating cytokine secretion, primarily in the context of lymphocytes(Arima et al., 2011; Kitamura et al., 2011; Liu et al., 2012; Muchamuel et al., 2009; Schmidt et al., 2018). For example, mutations in immunoproteasome subunits that limit the proteasomal activity increase IL-6 secretion(Arima et al., 2011). We therefore hypothesized that PSME4-mediated restriction of immunoproteasome activity in cancer cells might directly increase tumor IL-6 secretion. Indeed, blocking immunoproteasome activity with ONX0914 partially rescued the loss of IL6 secretion from PSME4-depleted KP1.9 cells, suggesting that modulation of immunoproteasome activity contributes to the effect of PSME4 on IL-6 expression (Figures 3G, S12G-I). Furthermore, inhibiting the immunoproteasome almost completely abrogated the increase in IL-17 secretion in PSME4-depleted cells (Figures 3H).

### PSME4 promotes an immunosuppressive tumor microenvironment

As tumor inflammation and antigenicity are key to anti-tumor immunity, we examined the effect of PSME4 *in vivo* by injecting mice with KP1.9, KP1.9^PSME4 OE^ or KP1.9^PSME4_KD^ (Figures 4A). Notably, mice bearing KP1.9^PSME4 OE^ showed marked reduction in their body weight starting a month from tumor cell injection (33 days), as compared to mice bearing KP1.9 tumors (Figure S13A). Lungs of mice bearing KP1.9^PSME4 OE^ tumor were significantly larger, contained numerous lesions whereas lungs of mice bearing KP1.9^PSME4 KD^ did not show any detectable lesions (Figures 4B-C, S13B-D). These effects were not driven by a change in the proliferation rate of tumor cells (Figures S13E-F). Next, we analyzed the changes in the immune milieu by performing single cell RNA sequencing (sc-RNA seq) of CD45+ cells 3 weeks after the injection of parental KP1.9 or KP1.9^PSME4 KD^ tumor cells. This allowed us to examine the effect of PSME4 on the early immune response to the tumor, without bias due to dramatically different levels of tumor burden (Figures 4D-E S14A-B). The overall distribution of cell types was similar among groups (Figure S14A). However, the relative sizes of the alveolar and cycling macrophage clusters as well as *Ccr7*+ regulatory DCs, which have been previously shown to play an immunosuppressive role in NSCLC (Lavin et al., 2017; Maier et al., 2020), in the lung of mice bearing KP1.9^PSME4 KD^ tumors were smaller than in the lung of mice bearing KP1.9 tumors (Figures 4F-H). Interestingly, we concurrently observed a decrease in the secretion of the macrophage recruiting chemokine MIP-1α/CCL3 (Baba and Mukaida, 2014) from the KP1.9 tumor cells upon PSME4 depletion (Figure 3F). We also observed an increase in plasmacytoid DCs (pDCs) in the mice bearing KP1.9^PSME4 KD^ tumors (Figure 4I). While pDCs have been proposed to have both pro- and anti-inflammatory roles in cancers(Li et al., 2017), here we see the expression of *Il12a* and an enrichment for pathways related to antigen presentation and processing (Figure 4J, S15A-B), suggesting that pDCs serve a pro-inflammatory role in this context. This is in contrast to the mature DCs enriched in immunoregulatory molecules (mregDCs), which are decreased in KP1.9^PSME4_KD^ and express higher levels of *Il6* (Figure 4K, S15C-D). Likewise, the increased secretion of IFNγ upon PSME4 depletion was accompanied by an increase in the expression of IFN response gene *Ifi272a* in several cell types, including interstitial macrophages, cycling dendritic cells and Tregs (Figure 4L). Finally, to examine if there is a skew towards particular cell fates, we characterized the T cell populations and T cell receptors (TCR). We reasoned that TCR clones that were not unique were result of expansion (Figures. 4M, S16A-B). We found that cytotoxic T cell and regulatory T cell populations had considerably expanded (Figures S17A-B, S18A-D). There were more cytotoxic CD8 T cells in the KP1.9^PSME4 KD^ milieu whereas Tregs expanded in the KP1.9 bearing mice, with a significant difference in the Treg/CD8 T cell ratio between the two groups (Figure 4N). Together, our results suggest that upregulation of PSME4 may both drive immune evasion by reducing cytotoxic T cell activity and play an active role in tumorigenesis by promoting an immune-suppressed microenvironment *in vivo*.

**Fig. 4.**
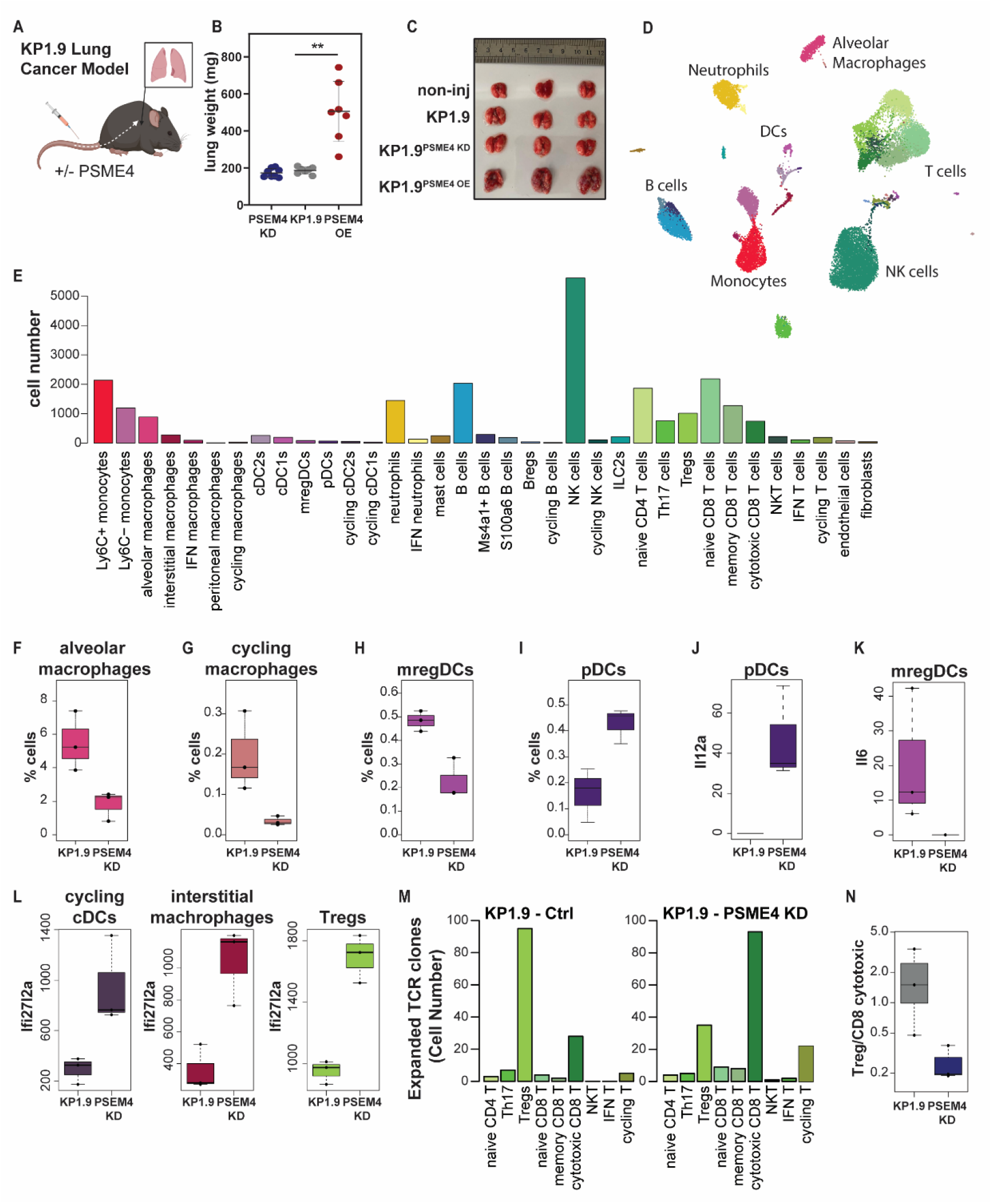
PSME4 promotes an immunosuppressive tumor environment. **(A)** Schematic of orthotopic lung tumor models by tail vein injection of KP1.9 cells overexpressing (KP1.9^PSME4 OE^) or deficient of (KP1.9^PSME4 KD^) PSME4 to C57/B6 mice (**B**) The lung weight of mice bearing parental, PSME4-overexpressing (OE), or PSME4-deficint (KD) KP1.9 tumors (**P = 0.0020). **(C)** Photograph of representative lungs from mice not injected with tumor cells (non-inj) 41 days after the injection of parental, PSME4 overexpressing (OE), or PSME4 deficient (KD) KP1.9 cells. **(D)** Uniform manifold approximation and projection visualization (UMAP) showing 35 cell populations across the three replicates from mice bearing KP1.9 (Ctrl) or KP1.9^PSME4 KD^ tumors. **(E)** Bar graph showing total number of cells per cell type identified across the three replicates from mice bearing KP1.9 (Ctrl) or KP1.9^PSME4 KD^ tumors (KD). **(F-I)** Boxplot showing relative abundance of alveolar macrophages (F), cycling macrophages (G), mature DCs enriched in immunoregulatory molecules (mregDCs) (H), and plasmacytoid DCs (pDCs) (I) in the lung of mice injected with KP1.9 (n = 3) or KP1.9^PSME4 KD^ (PSEM4KD) (n = 3) cells. Boxplot shows 25th to 75th percentiles with the 50th denoted by a line; whiskers show 1.5× interquartile range, or maximum or minimum if smaller. **(J, K)** Boxplot showing pseudobulk tpm expression of *Il6* in mregDCs (J) or expression of *Il12a* in pDCs (K) in the lung of mice injected with KP1.9 (n = 3) or KP1.9^PSME4 KD^ (PSEM4KD) (n = 3) cells. **(L)** Boxplots showing pseudobulk tpm expression of *Ifi27l2a* in cycling cDCs, interstitial macrophages and regulatory T cells in the lung of mice injected with KP1.9 (n = 3) or KP1.9 PSEM4KD) (n = 3) cells. **(M)** Barplot showing number of TCR clones which were expanded (at least 2 cells sharing the same TCR clone) in the different T cell populations in a representative samples from mice injected with KP1.9 or KP1.9 PSEM4 depleted (PSEM4KD) cells. **(N)** Ratio of number of expanded (i.e. having non-unique TCR) Tregs to cytotoxic CD8 ratio in each sample.

To further validate the changes observed by scRNA-seq, we analyzed the lung of tumor-bearing mice and stained for tumor infiltrating lymphocytes (TILs, Figure 5A, S19A-D). Here too, we found that KP1.9^PSME4 OE^ tumors exhibited a decrease in the CD8^+^ T cell/Treg ratio (Figures 5A-B). Moreover, our analysis revealed a significant increase in exhausted PD1^+^ CD8^+^ T cells (Figure 5C and S19A) and a decrease in IFNγ in tumor infiltrating lymphocytes (TILs) in KP1.9^PSME4 OE^ tumor-bearing mice (Figure 5D-E), suggesting decreased tumor inflammation. Furthermore, we observed a decrease in naïve CD8^+^ T cells (CD62L+, CD44-; Figures 5F-G, S20B-C) suggesting that KP1.9^PSME4 OE^ exhibit reduced inflammation and T cell activation. To determine whether the PSME4-induced effect on tumor growth was indeed immune-mediated, we repeated the experiment using RAG1-deficient mice, which lack lymphocytes. In contrast to WT mice where the KP1.9-bearing mice lost weight more quickly than the KP1.9^PSME4 KD^, no differences in weight loss or survival between mice bearing PSME4-deficient and those bearing KP1.9 tumors (Figure S21) were observed, further corroborating that the PSME4-driven changes in tumor progression is mediated by lymphocytes *in vivo*. To further consolidate this point we tested the ability of splenocytes isolated from tumor-bearing mice to kill the tumor cells. We found that splenic lymphocytes from mice bearing KP1.9^PSME4 OE^ tumors exhibited reduced propensity to kill KP1.9 cells (Figures 5H S22A). By contrast, splenic lymphocytes from mice bearing KP1.9^PSME4 KD^ tumor demonstrated killing even better than those from mice bearing KP1.9 tumors (Figure 5H). These results indicated that the anti-tumor activity we observed indeed depended on loss of PSME4. Accordingly, we found a significant increase in a Th17-associated cytokine signature (IL-22, IL-17a and IFNγ), which is associated with promoting CD8 cell anti-tumor responses(Knochelmann et al., 2018), in the secretome of splenic lymphocytes from mice bearing KP1.9^PSME4 KD^ tumors (Figures 5I-K, S22B-D). These findings were in agreement with our observation that depletion of PSME4 induces the secretion of IL-6, a key cytokine regulating the Treg / Th17 balance by inhibiting Treg and promoting Th17 differentiation (Kimura and Kishimoto, 2010).

**Fig 5.**
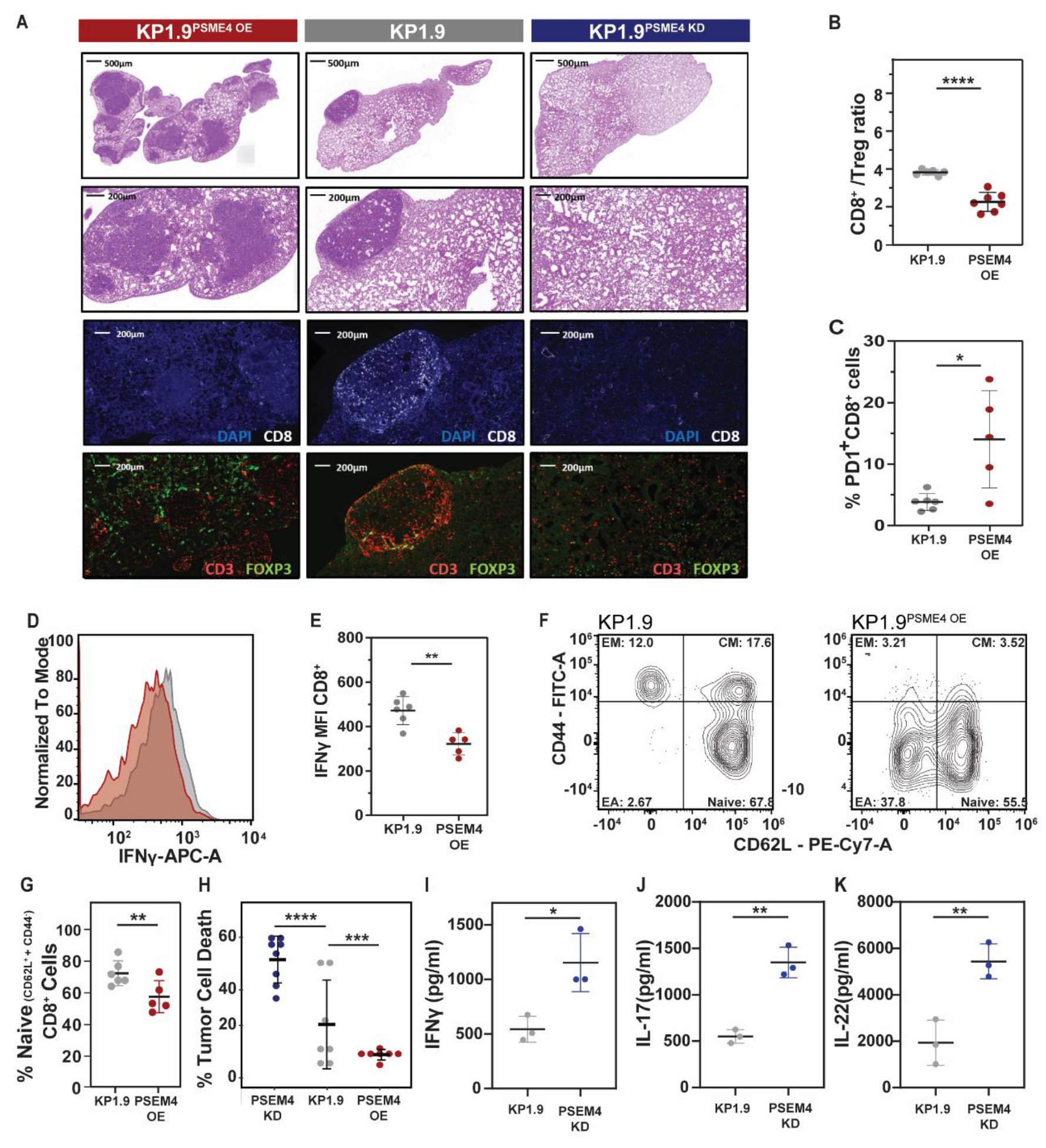
PSME4 abrogates anti-tumor immunity. **(A)** H&E staining of lungs from mice bearing KP1.9, KP1.9^PSME4 OE^, or KP1.9^PSME4 KD^ tumors (top) and immunofluorescence staining of the tumor sections with indicated antibodies (bottom). **(B)** The ratio of CD8 T cells to Tregs in the spleens of mice bearing KP1.9^PSME4 OE^ or KP1.9 tumors (****P <0.0001). **(C)** The percent of PD1 positive CD8 T cells in the lung of the mice bearing KP1.9^PSME4 OE^ or KP1.9 tumors (*P = 0.0440). **(D)** A histogram of IFNγ stained CD8 T cells of mice bearing KP1.9 (grey) or KP1.9^PSME4 OE^ (pink) tumors. **(E)** The median fluorescent intensity (MFI) of IFNγ stained CD8 T cells in the lung of mice bearing KP1.9^PSME4 OE^ or KP1.9 tumors (E; *P = 0.0017). **(F-G)** The percent of different subsets of CD8-positive lymphocytes (naïve, early activated, effector memory [EM] or central memory [CM], which are CD62L and/or CD44 positive, in the lung of mice bearing KP1.9^PSME4 OE^ or KP1.9 tumors. **(H)** The percent of KP1.9 cells co-cultured with splenic lymphocytes from mice bearing KP1.9, KP1.9^PSME4 OE^ or KP1.9^PSME4 KD^ tumors that is dead (****P <0.0001, ***P = 0.0021). (**I-K)** The amount (pg/ml) of IFNγ (I), IL-17 (J) or IL-22 (K) secreted from splenic lymphocytes from mice bearing KP1.9 or KP1.9^PSME4 OE^ tumors (*P<0.05, **P<0.01).

### PSME4 drives a ‘cold’ tumor signature in NSCLC and is associated with reduced response to ICI across multiple cancer types

Our findings that PSME4 plays an anti-inflammatory role in cancer raise the intriguing possibility that it promotes immune evasion in cancer and may be associated with resistance to immunotherapy. Indeed, NSCLC with high levels of PMSE4 had reduced signature of T cell infiltration and inflammation (Figures 6A, S23). PSME4 and immunoproteasome levels vary greatly among different types of cancer (Figures. 6B and S24A). Therefore, we analyzed RNA-seq data from the CPI1000+ cohort (Litchfield et al., 2021), comprising 6 different patient cohorts of three different cancer types treated with ICI (Van Allen et al., 2015; Hugo et al., 2016; Mariathasan et al., 2018; McDermott et al., 2018; Riaz et al., 2017b; Snyder et al., 2014) and found that PSME4 varies greatly among individual tumors and cancer types and that tumors with high expression of PSME4 were less likely to respond to immunotherapy (Figures. 6C-D). This was true within each cancer type examined separately, and when response rate was measured across the CPI1000+ cohort (ref) (Figure S24B). Specifically, melanoma, in which the immunoproteasome has been reported to be upregulated and associated with response to therapy(Harel et al., 2019; Riaz et al., 2017a), expressed PSME4 lowly on average whereas renal and bladder cancers exhibited a wide range of expression for both the immunoproteasome and PSME4 (Figure 6C). Since we showed that PSME4 attenuates immunoproteasome activity, we hypothesized that the ratio between PSME4 and the immunoproteasome levels, and not their absolute level, may be a better measure to examine response to ICI. Furthermore, we found that in the PSME4-low cellular state, PSMB10 is the immunoproteasome subunit that is most increased (Figure 3B), so we chose it to categorize immunoproteasome levels in the cohorts. Notably, the ratio of PSME4 to PSMB10 yielded a more significant association to the response to ICI than either subunit alone (Figure 6E, S24B-C). To control for the potential bias of data pooling, we also calculated the effect size across the cohorts by combining the individual effect sizes of each biomarker in each cohort (Figure 6F and S25A-B). PSME4, PSMB10 and the ratio between them all had strong effect sizes on responsiveness, even in comparison with tumor mutational burden, previously shown to be one of the strongest classifiers of responsiveness to ICI (Litchfield et al., 2021). Furthermore, we showed that PSME4 did not strongly correlate with other biomarkers from the cohort and that is still had a significant contribution to a general linear regression model of responsiveness even when the other significant biomarkers, such as TMB were included (Figure S25A; PSME4 - P = 0.0194).

**Fig 6.**
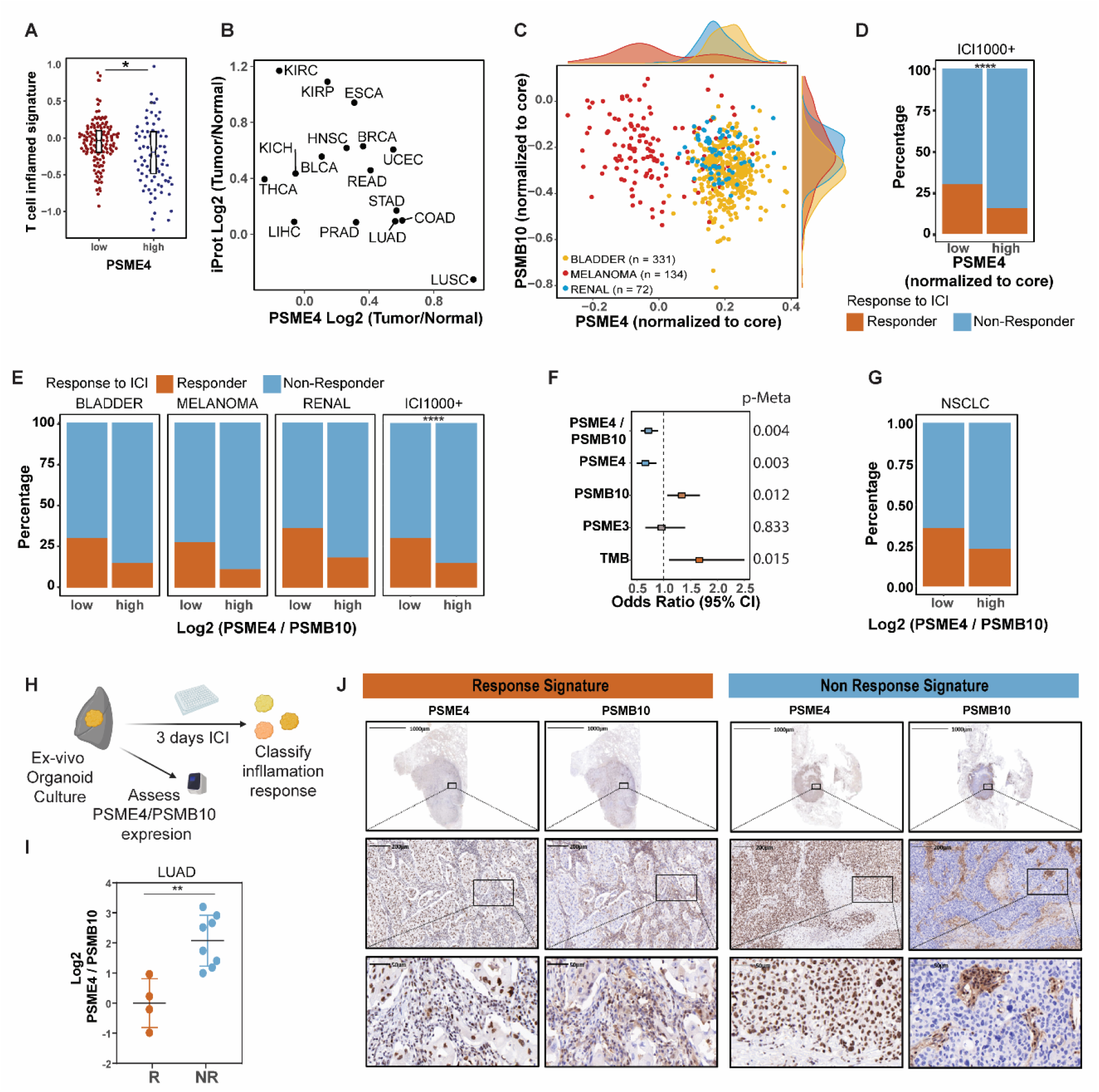
Responsiveness to ICI is reduced in PSME4-high tumors across multiple cancer types. **(A)** T-cell inflammation signature was calculated for the LUAD samples in CPTAC stratified according to the PSME4 expression level. **(B)** The ratio of PSME4 expression in the tumor versus normal tissue from the indicated TCGA cohort is plotted against the ratio of the mean of the immunoproteasome (iProt) subunits PSMB8-10 and PSME1-2 in the tumor versus normal tissue. **(C-E)** The expression of PSME4 normalized to the core proteasome subunits is calculated for bladder cancer, melanoma and renal tumors across the Hugo, Mariathasan, McDermot, Riaz, Snyder, and Van Allen cohorts. PSME4 is plotted against PSMB10 (C). PSME4 normalized to the core proteasome subunits (D) or the ratio of PSME4 and PSMB10 expression (E) was used to stratified patients (upper and lower 50% of samples). The percentage of responders and non-responders to ICI was counted for each group per cancer type and across the CI1000+ cohort (χ2 test D****P= B**** P = 0.000096, E**** P = 0.000016). **(F)** The effect size (calculated as the log2 odds ratio for response versus no response) and significance of the proteasome subunit biomarkers and Tumor Mutational Burden (TMB) in meta-analysis across all cohorts incorporating the effect sizes and standard errors from each individual cohort. The meta p values (p-Meta) are shown on the right. (**G**) The ratio of PSME4 and PSMB10 expression was used to stratified patients who responded to ICI in the NSCLC Kim cohort. **(H)** Workflow of the Ex-Vivo Organoid Culture System (EVOC). A fraction of each resected tumor was assessed for PSME4 and PSMB10 expression using quantitative PCR and the rest was treated for 3 days with immune checkpoint inhibitors (ICI). Response-like signature to ICI was classified by IFNγ expression. **(I)** The ratio of PSME4 to PSMB10 expression determined by qPCR in patient tumors with EVOC ICI responder or non-responder signature (** P = 0.0056). **(J)** Immunohistochemistry for PSME4 and PSMB10 of sections with an ICI response-signature and an ICI non-response-signature EVOC samples.

To demonstrate that the stratification of patients by the ratio of PSME4 and PSMB10 expression translates to NSCLC we validated the signature we observed in the ICI1000+ cohort in a 27 patient NSCLC cohort (Kim et al., 2020). Indeed, we saw that NSCLC patients with low levels of PSME4 in the tumor had a higher ICI response rate, consistent with the findings in Melanoma, Bladder and Kidney cancer, albeit not significantly due to the small size. To strengthen this observation, we utilized an Ex-Vivo Organoid Culture model (EVOC(Kamer et al., 2021)), which contain tumor tissue, infiltrating lymphocytes and stroma cells (Figure 6G, S26A-B). We assessed the expression of PSME4 and PSMB10 as well as stained for CD8^+^ T cell infiltrates and PDL1 from a section of the resected tumor (Figure S26A-B). In parallel, we tested the IFNγ levels following treatment with either Durvalumab alone (αPD-L1) or together with Ipilimumab (αCTLA-4) in the EVOCs. IFNγ levels (Kamer et al., 2021) were then used to define a response-like signature in the EVOCs. Notably, we found that tumors with responder hallmarks had significantly lower PSME4/PSMB10 ratios compared to non-responders (Figures 6H-I, S26C-F) and that neither PSME4 nor PSMB10 individually associated with the response (Figures S26H-I). This result did not merely reflect immune cell infiltration as both PSME4 and PSMB10 were expressed in the epithelial tissue, and PSMB10 was more highly expressed in the lymphocyte infiltrates, consistent with the known expression of the immunoproteasome in immune cells (Figure 6I).

## Discussion

We show that altered proteasome composition and function shape tumor-host interactions in NSCLC and that PSME4-capped proteasome plays an anti-inflammatory role in cancer by attenuating immunoproteasome activity. The attenuated immunoproteasome activity in turn leads to reduced antigenicity by restricting immunopeptidome diversity and altering surface HLA presentation as well as reducing cellular inflammation in an IL-6 dependent manner. While label-free quantitative MS analysis of cell lines suggested that <5% of 20S proteasomes bear PSME4 regulators (Fabre et al., 2014; Morozov and Karpov, 2019), our results suggest that this frequency is increased in many cancer types. As PSME4 is associated with DNA damage and histone organization, we speculate that it may be upregulated to offer a protective mechanism to cope with the associated proteotoxic stress and DNA damage (Ahmed, 2019; Kammerl et al., 2019; M. Cantin and V. Richter, 2012; Srinivas et al., 2019). Further, as PSME4 is known to play a nuclear role and was suggested to modulate transcription (Jiang et al., 2021) it will be intriguing to comprehensively examine the effect its upregulation has on the transcriptional landscape, which is beyond the scope of our study. Nevertheless, the dominant effect PSME4 has on the degradation landscape, and the altered catalytic activity of the immunoproteasome even in cell-free systems, strongly suggest that PSME4 exerts its effect also by directly modulating proteasome activity, post-translationally.

Moreover, we found PSME4 to be the first known example of a regulator differentially modulating the activities of the constitutive proteasome and immunoproteasome. PSME4 is one of a small number of markers of resistance to ICI, as opposed to markers of responsiveness such as the immunoproteasome, TMB, infiltrate signatures and others. As such, it highlights the potential of targeting PSME4 expression or its binding to proteasomes as a novel therapeutic approach in treating NSCLC and in sensitizing response to ICI. Because PSME4 levels are tumor-type specific, such approaches are expected to differ between cancer types. Together, our results highlight the degradation landscape and the balance between different proteasome compositions as an understudied factor in pathogenesis of human diseases in general and cancer proteostasis in particular. This novel proteasome-dependent mechanism of immune evasion allows tumors to abrogate anti-tumor immunity is of particular interest as it provides another layer to the understanding of tumor proteostasis control.

## Materials and Methods

### Purification of proteasome complexes

Lung Adenocarcinoma tumors and adjacent tissues were mechanically disrupted and passed through a 70 µm cell strainer 93070 (SPL). Cells were lysed with 25 mM HEPES, pH 7.4, 10% glycerol, 5 mM MgCl2, 1 mM ATP, and 1:400 protease-inhibitor mixture (Calbiochem), then homogenized through freeze–thaw cycles and passed through a needle. The lysates were cleared by 30-min centrifugation at 21,130g at 4 °C. Lysates were treated with 2 mM 1,10-phenanthroline (Sigma), cross-linked with 0.5 mM DSP (Thermo Fisher Scientific) for 30 min at room temperature, and quenched in 100 mM Tris-HCl, pH 8, 5 mM L-cysteine for 10 min at room temperature. For immunoprecipitation, the lysates were then incubated with Protein G–Sepharose beads (Santa Cruz) with antibodies to PSMA1 and eluted with 100 mM Tris-HCl, pH 8, 8M urea and 50 mM DTT for 30 min at 37 °C. Subsequently, 1% trifluoroacetic acid (TFA) was added. Aliquots of each elution fraction were analyzed by SDS–PAGE to evaluate yield and purity.

### Purification and concentration of proteasome peptides

A critical step in our procedure is the separation of peptides from the proteins eluted in the proteasome pulldown. MAPP analyzes endogenously cleaved peptides, whereas the proteasome complex and associated proteins are physically excluded. Immunoprecipitated proteasomes and their encompassed peptides were loaded on C18 cartridges (Waters) that were prewashed with 80% acetonitrile (ACN) in 0.1% TFA, then washed with 0.1% TFA only. After loading, the cartridges were washed with 0.1% TFA. Peptides were eluted with 30% ACN in 0.1% TFA. Protein fractions were eluted with 80% ACN in 0.1% TFA.

### Mass spectrometry sample processing

Protein fraction after proteasome purification; Samples were loaded onto 3 kDa molecular weight cut-off spin columns. Volume was reduced to 25μL by centrifugation at 14,000g for 10min. 175μL 8M urea was added and centrifuged at 14,000g for 10min. Filters were reversed and centrifuged to extract the proteins. Proteins were reduced with 5 mM dithiothreitol (Sigma) for 1hr at room temperature and alkylated with 10 mM iodoacetamide (Sigma) in the dark for 45 min at room temperature. Samples were diluted to 2M urea with 50mM ammonium bicarbonate. Proteins were then subjected to digestion with trypsin (Promega; Madison, WI, USA) overnight at 37°C at 50:1 protein:trypsin ratio, followed by a second trypsin digestion for 4 hr. The digestions were stopped by addition of trifluroacetic acid (1% final concentration). Following digestion, peptides were desalted using Oasis HLB, μElution format (Waters, Milford, MA, USA). The samples were vacuum dried and stored in -80°C until further analysis.

Total proteomics; Lysates in 5% SDS in 50 mM Tris-HCl were incubated at 96 °C for 5 min, followed by six cycles of 30 s of sonication (Bioruptor Pico, Diagenode, USA). Proteins were reduced with 5 mM dithiothreitol and alkylated with 10 mM iodoacetamide in the dark. Each sample was loaded onto S-Trap microcolumns (Protifi, USA) according to the manufacturer’s instructions. In brief, after loading, samples were washed with 90:10% methanol/50 mM ammonium bicarbonate. Samples were then digested with trypsin for 1.5 h at 47 °C. The digested peptides were eluted using 50 mM ammonium bicarbonate; trypsin was added to this fraction and incubated overnight at 37 °C. Two more elutions were made using 0.2% formic acid and 0.2% formic acid in 50% acetonitrile. The three elutions were pooled together and vacuum-centrifuged to dry. Samples were kept at −80 °C until analysis.

HLA immunopeptidomics was performed as described previously (Kalaora et al., 2020). In brief, cell-line pellets were collected in triplicate from 2×10E8 cells. Cell pellets were lysed on ice with a lysis buffer containing 0.25% sodium deoxycholate, 0.2 mM iodoacetamide, 1 mM EDTA, 1:200 Protease Inhibitor Cocktail (Sigma-Aldrich), 1 mM PMSF and 1% octyl-b-D glucopyranoside in PBS. Samples were then incubated in rotation at 4°C for 1 hour. The lysates were cleared by centrifugation at 48,000 g for 60 minutes at 4°C and then passed through a pre-clearing column containing ProteinA Sepharose beads. HLA-I molecules were immunoaffinity purified from cleared lysate with the panHLA-I antibody (W6/32 antibody purified from HB95 hybridoma cells). Affinity columns were washed first with 400 mM NaCl, 20 mM Tris–HCl and then with 20 mM Tris–HCl pH 8.0. The HLA-peptide complexes were then eluted with 1% trifluoracetic acid followed by separation of the peptides from the proteins by binding the eluted fraction to Sep-Pak (Waters). Elution of the peptides was done with 28% acetonitrile in 0.1% trifluoracetic acid. The peptides were dried by vacuum centrifugation.

### Liquid chromatography mass spectrometry

Peptide fraction; ULC/MS grade solvents were used for all chromatographic steps. Each sample was loaded using split-less nano-Ultra Performance Liquid Chromatography (10 kpsi nanoAcquity; Waters, Milford, MA, USA). The mobile phase was: A) H2O + 0.1% formic acid and B) acetonitrile + 0.1% formic acid. Desalting of the samples was performed online using a reversed-phase Symmetry C18 trapping column (180 µm internal diameter, 20 mm length, 5 µm particle size; Waters). The peptides were then separated using a T3 HSS nano-column (75 µm internal diameter, 250 mm length, 1.8 µm particle size; Waters) at 0.35 µL/min. Peptides were eluted from the column into the mass spectrometer using the following gradient: 4% to 35%B in 120 min, 35% to 90%B in 5 min, maintained at 90% for 5 min and then back to initial conditions.

The nanoUPLC was coupled online through a nanoESI emitter (10 μm tip; New Objective; Woburn, MA, USA) to a quadrupole orbitrap mass spectrometer (Q Exactive Plus, Thermo Scientific) using a FlexIon nanospray apparatus (Proxeon).

Data was acquired in data dependent acquisition (DDA) mode, using a Top10 method. MS1 resolution was set to 70,000 (at 400m/z), mass range of 375-1650m/z, AGC of 3e6 and maximum injection time was set to 100msec. MS2 resolution was set to 17,500, quadrupole isolation 1.7m/z, AGC of 1e5, dynamic exclusion of 40sec and maximum injection time of 150msec.

### Mass spectrometry data analysis

Raw data were analyzed in MaxQuant software (version 1.6.0.16) with the default parameters for the analysis of the proteasomal peptides, except for the following: unspecific enzyme, LFQ minimum ratio count of 1, minimum peptide length for unspecific search of 6, maximum peptide length for unspecific search of 40, and match between runs enabled. A stringent false discovery rate (FDR) of 1% was applied for peptide identification. For the analysis of tryptic digests, the default parameters were set, apart from a minimum peptide length of 6. Masses were searched against the human proteome database from UniprotKB (last update September 2018).

### Proteomics processing and label-free quantification

Peptides resulting from MaxQuant were initially filtered to remove reverse sequences and known MS contaminants. For MAPP peptide fraction we removed antibody and proteasome peptides as contaminants. To decrease ambiguity, we allowed peptides that had at least two valid LFQ intensities out of sample replicates, and we included razor peptides, which belong to a unique MaxQuant ‘protein group’. MAPP protein intensities were inferred with MaxQuant. For graphical representation, intensities were log transformed, and in Python v3.6, zero intensity was imputed to a random value chosen from a normal distribution of 0.3 s.d. and downshifted 1.8 s.d. For clinical cohorts, zero intensity was imputed to half the minimum. The presence of missing values reflects both technical and biological variation in the samples. In cases of matched samples ratios were calculated per pair and paired t tests were used. Otherwise, ratios were calculated based on the median of each group and a non-paired student t test was used. The protein fraction of the MAPP analysis was normalized to the mean of the core proteasome subunits (PSMA1-7 and PSMB1-4) to control for efficiency. For comparison of WC proteomics and MAPP data, proteins were ranked by their signed pvlaue: the sign of the fold-change between conditions multiplied by the negative log 10 transformed p value. Proteins which had positive (tumor-increased) values were binned into 10 groups of equal size. Any proteins which were detected in MAPP but not detected or defined as adjacent -associated in WC proteomics were termed not-detected. The proteins in MAPP or WC proteomics with the top 50 signed p values were presented in heatmaps in Figure 1. For the immunopeptidomics, values from a peptide that was not detected in a treatment pair were not included in the analysis. The binding of peptides was predicted by netMHC based on A549 haplotypes as described previously (Javitt et al., 2019b). Any peptide with a binding rank greater than 5 was considered as a contaminant for analysis of the differentially presented peptides.

### TCGA and CPTAC analysis

The cancer genome atlas (TCGA) data was mined using the xenaPython package in Python 3.6. The results shown in this analysis are in whole or part based upon data generated by the TCGA Research Network: http://cancergenome.nih.gov/. The full lung carcinoma cohort (both lung adenocarcinoma [LUAD] and lung squamous cell carcinoma [LUSC] designations, n = 1128, 3A and S10) or the LUAD cohort alone (n = 576,S2) as indicated. For the cross-cancer proteasome ratio the GDC pan cancer dataset was used. PSME4 high tumors in Figure 6 and Figures. S24-25 were defined as those with above average expression of PSME4. Only cancer types with 10 or more normal controls were used. Ratios presented are the difference between the means of the normal and tumor groups. Data used in this publication were generated by the Clinical Proteomic Tumor Analysis Consortium (NCI/NIH). The CPTAC LUAD cohort (n = 213) was used. In cases where ratios between the adjacent and tumor samples are presented only those samples with matching controls are used. Otherwise, all the tumor and normal (adjacent control) samples were utilized. The T cell inflammation signature was based on Spranger et al. (Spranger et al., 2015; Trujillo et al., 2018).

### Immune checkpoint inhibitor response analysis

Data reprocessing of six cohorts of ICI treated patients across three cancer types (Van Allen et al., 2015; Hugo et al., 2016; Mariathasan et al., 2018; McDermott et al., 2018; Riaz et al., 2017b; Snyder et al., 2014) was performed as described previously (Litchfield et al., 2021). The expression of different proteasome subunits was examined across the samples. Where noted, proteasome subunit expression was normalized to the core proteasome subunits (PSMA1-7 and PSMB1-4) for each sample. Patients were then stratified by the expression of different subunits (50 percent highest and lowest expressers) and this was correlated with response to ICI which was defined as in (Litchfield et al., 2021). To avoid data pooling a meta-statistics approach was used as described in (Litchfield et al., 2021). Likewise, to confirm the independence of different biomarkers a general linear regression model was used. The model was first generated for 35 biomarkers and then iteratively reduced to only the markers which significantly contributed to the classification. The signatures were confirmed in an independent cohort of NSCLC (Kim et al., 2020).

### Statistical analysis and data visualization

Statistical analyses were performed in R v 3.6.2 and GraphPad Prism v 7.04. In R data was visualized using the complexheatmap (Zuguang Gu et al., 2016) and ggplot (Wickam, 2016) packages. Defaults were used unless otherwise noted. Protein annotation and gene ontology analysis were performed with the gene-set enrichment analysis and protein subcellular localization was extracted from the Human protein atlas (Thul et al., 2017; Uhlen et al., 2017; Uhlén et al., 2015) with “uncertain” or “approved” localization reliabilities excluded.

### Antibodies and plasmids

Rabbit anti PSME4 HPA060922 1:1000 WB, 1:300 IF (Sigma), Rabbit anti PSMD11 ab99414 1:1000 WB, Mouse anti proteasome 20s PSMA1-7 subunits ab22674 1:1000 WB, Rabbit anti PSMB10/MECL1 ab183506 1:1000 WB, Mouse anti β Actin ab170325 1:5000 WB (Abcam), Rabbit anti PA28alpha/PSME1 9643 1:1000 WB, Rabbit anti PSMB8/LMP7 13635 1:1000 WB and Rabbit anti PA28γ 2412 1:1000 WB (Cell Signaling).

human HLA-A,B,C-PE (W6/32, Biolegend 311406), FC, 1:20; mouse CD45.2-PerCP/Cy5.5 (104, eBioscience 45-0454-82), FC, 1:200; mouse CD3e-Super Bright 436 (145-2C11, eBioscience 62-0031-82), FC, 1:100; mouse CD4-APC/Cy7 (GK1.5, Biolegend 100414), FC, 1:200; mouse CD8a-Super Bright 702 (53-6.7, Biolegend 100748), FC, 1:200; mouse CD279 (PD-1)-PE (RPM1-30, eBioscience 12-9981-81), FC, 1:100; mouse IFNg-APC (XMG1.2, eBioscience 17-7311-82), FC, 1:160; mouse/human CD44-FITC (IM7, Biolegend 103005), FC, 1:200; mouse CD62L-PE/Cy7 (MEL-14, Biolegend 104418), FC, 1:6000; mouse FoxP3-Pe (FJK-16s, eBioscience 12-5773-82), FC, 1:200; mouse H-2Db-FITC (KH95, Biolegend 111505), FC, 1:200; mouse H-2kb-PE (AF6-88.5.5.3, eBioscience 12-5958-82), FC, 1:500; Zombie Aqua (Biolegend 423102), FC, 1:300; Propidium Iodide (Sigma Aldrich P4170), FC, 1 ug/mL; CFSE (Biolegend 423801), FC, 5 uM. Mouse anti alpha-6, produced from hybridoma was a kind gift from Keiji Tanaka. IgG1k isotype antibody (BioLegend, BLG-400139) was used as a control of MAPP method.

Goat anti Mouse 488 A11029 1:400, Goat anti mouse 647 A31571 1:400 (Invitrogen), Goat anti Rabbit 647 AB150075 1:400 (Abcam), Goat anti mouse HRP 115-035-205 1:5000, Goat anti Rabbit HRP 111-035-003 1:5000 (Jackson labs).

MISSION shRNA targeting mouse or human PSME4 or RFP were obtained from Sigma (TRCN0000176569, TRCN0000178428, TRCN0000158223, TRCN0000157073).

pcDNA3.1_PSME4 (Welk et al., 2019) was transfected into A549/KP1.9 cells with Lipofectamine 2000 (Thermo Fisher).

### Cell culture and drug treatments

A549 were originally obtained from ATCC grown in DMEM and used as model systems for cell biology studies including western blot, biochemistry, and imaging. KP1.9, kindly provided by Alfred Zippelius, were grown in Iscove’s MDM. Cells were routinely tested for mycolplasma contamination. Media were supplemented with 10% fetal bovine serum, 1% Penicillin/streptomycin and L-glutamine (2mmol/l) (Biological industries), unless otherwise indicated, at 37 °C with 5% CO2.

For TNF-α/IFN-γ activation cells were allowed to seed overnight, and subsequently treated with TNF-α (PeproTech, human; 300-01A or mouse; 315-01A) and IFN-γ (PeproTech, human; 300-02 or mouse; 315-05) at 20 ng/ml and 10 ng/ml, respectively, for 24h.

Cell death assessment was done by trypan blue staining and counted with Countess II™ automated cell counter (Thermo Fisher) and by Cell titerGlo assay (Promega). Proliferation of mammalian cells were measured by KIT8 assay (Sigma).

### Proteasome cleavage reporter assay

Cells were lysed with 25 mM HEPES, pH 7.4, 10% glycerol, 5 mM MgCl2, 1 mM ATP, and 1:400 protease-inhibitor mixture (Calbiochem), then homogenized through freeze–thaw cycles and passed through a needle. The lysates were cleared by 30-min centrifugation at 21,130g at 4 °C to remove cell debris. Protein concentration of the supernatant was determined by a NanoDrop spectrophotometer (Thermo Fisher Scientific) by measuring the absorbance at 280 nm. Proteasomal activity in cell fractions was determined by cleavage of the fluorogenic precursor substrate Suc-Leu-Leu-Val-Tyr-AMC (Suc-LLVY-AMC), Ac-Pro-Ala-Leu-AMC (Ac-PAL-AMC), Z-Leu-Leu-Glu-AMC (Z-LLE-AMC), Ac-Arg-Leu-Arg-AMC (Ac-RLR-AMC), Ac-Ala-Asn-Trp-AMC (Ac-ANW-AMC) (Boston Biochem), Z-Leu-Leu-Glu-βNA (Z-LLE-βNA), Ac-Nle-Pro-Nle-Asp-AMC (Ac-nLPnLD-AMC) (Bachem). 10 μM substrate was added to 10 µg total protein/well, and incubated in a reaction buffer (50 mM Hepes, pH 7.5, 1 mM DTT, 5 mM MgCl2 and 2 Mm ATP). When using recombinant proteins such as PSME4 [purified as described in(Toste Rêgo and da Fonseca, 2019)] or PSME1 (AB-206168, abcam) 1 nM of each protein was add to 100 ul final reaction. Fluorescent increase resulting from degradation of peptide-AMC at 37°C was monitored over time by means of a fluorometer (Synergy H1 Hybrid Multi-Mode Microplate Reader, BioTek) at 340 nm excitation and at 460 nm emission, using the proteasome inhibitor, MG132 (Calbiochem) as background. Resulting product curves were followed for up to 3.5 hours. Each value of fluorescence intensity represents a mean value obtained from three independent experiments.

### Immuno-blotting

Cells were lysed in STET buffer (50 mM Tris-HCl, pH 87.5, 150 mM NaCl, 2 Mm EDTA 1% Triton-X and1:400 protease-inhibitor mixture (Calbiochem)). Protein concentration was assessed using BC Assay Protein Quantitation Kit (Interchim). 20 μg of total protein was separated by SDS– PAGE on 4-20% gradient ExpressPlus PAGE M42015 (GenScript) and transferred onto PVDF membranes using iBlot™ 2 Gel Transfer Device (Thermo Fisher Scientific). The membranes were blocked in 5% milk prepared in TBS-0.1% Tween and incubated in primary antibodies overnight at 4°C followed by washing and incubation with secondary antibody. Blots were developed using the ChemiDoc XRS+ Imaging System (Bio-Rad) and band intensities were quantified with imageJ analyzer software.

### Immuno-precipitation

Cells lysed in 25 mM HEPES, pH 7.4, 10% glycerol, 5 mM MgCl2, 1 mM ATP, and 1:400 protease-inhibitor mixture (Calbiochem), then homogenized through freeze–thaw cycles and passed through a needle. The lysates were cleared by 30-min centrifugation at 21,130g at 4 °C to remove cell debris. Pre-cleared cell lysates were prepared by incubating with Protein A/G conjugated agarose beads for 30 min at 4°C with slow rotation. 1 mg Pre-cleared cell lysates were then added to 20 µL of fresh Protein A/G conjugated agarose beads along with 4 µg of antibodies (as described). The mixture was incubated over-night at 4°C with slow rotation followed by three 1 ml washes with chilled PBS buffer. The target conjugates were then eluted from beads by mixing protein loading dye (with reducing agents) and heating for 5 min at 95°C followed by denaturing Western blot.

### Gel Filtration

Gel filtration experiments were performed using Superose^®^ 6 Increase 10/300 GL size exclusion column (GE Healthcare) ÄKTA pure protein purification system (Cytiva). The running buffer used was 25 mM HEPES, pH 7.4, 10% glycerol, 5 mM MgCl2 and 1 mM ATP. The column was calibrated using gel filtration molecular weight standard (Bio-Rad). The following standards were used for calibration: Thyroglobulin (670 kDa, Rs=8.6 nm), γ-globulin (158 kDa, Rs=5.1 nm), Ovalbumin (44 kDa, Rs=2.8 nm), Myoglobin (17 kDa, Rs=1.9 nm), Vitamin B12 (1.35 kDa), and Dextran blue (2 MDa).

A 0.1 ml protein sample at a final concentration of 0.4 mg/ml was filtered and chromatographically analyzed using a flow rate of 0.5 ml/min. Absorbance was monitored at 280 nm, elution volumes were determined from UV chromatogram. The partition coefficient, Kav, was calculated from the elution volume of the sample, Ve, and total bed volume, Vt, using the expression: *Kav* = (*Ve* − *V*0)/(*Vt* − *V*0). Calibration curves and equations were established.

### Surface HLA staining in flow cytometry

A549 and KP1.9 cells were treated with hTNFα/hIFNγ (as described above) or mTNFα/mIFNγ (as described above), respectively, for 24 hr. Cells were harvested and stained with Live/Dead Aqua staining kit from BioLegend as per manufacturer’s protocol. Approximately 1×10^6^ cells were washed, blocked with 2% FCS for 5 min and stained with anti-HLA-A/B/C-PE (A549) or anti-MHCI-Kb-PE/MHCI-Db-FITC (KP1.9) in 100ul for 30 min. Cells were washed and acquired.

### Tissues fixation, processing and embedding

Tissues were fixed in 4% paraformaldehyde for one week. Dehydration of the tissue was done by serial immersion in increasing concentrations of alcohol (70 to 100%) and removal of the dehydrant with Xylene. The tissue was embedded in paraffin, sectioned at 4 µm and mounted on microscope slides. The slides were placed in an oven for one hour at 60°C or overnight at 37°C before immunohistochemically staining.

### Immunohistochemistry (IHC) and Immunofluorescence staining

H&E stains were performed on an automated device according to manufacturer’s instructions. IHC stains were performed on a Benchmark XT staining module (Ventana Medical Systems Inc.; USA) using iVIEW DAB Detection Kit (760-091, Ventana Medical Systems Inc.) or Ultra VIEW Universal DAB Detection Kit (760-500, Ventana Medical Systems Inc.; USA). Antibody details are described above. Following immunostaining, sections were counterstained with hematoxylin (Ventana Medical Systems Inc.), rinsed in distilled water and dehydrated manually in graded ethanols. Finally, the sections were cleared in xylene and mounted with Entellan (Surgipath Medical Industries Inc) on glass slides.

The immunofluorescent staining was performed at Molecular Cytology Core Facility of MSKCC using Discovery XT processor (Ventana Medical Systems). Sections were stained with antibodies for CD31 (endothelial cell marker), CD3 (T cells), CD8 (T cells) and FoxP3 (Tregs). Scanning of slides was done by PANNORAMIC SCAN II (3DHISTECH).

### IHC staining PSME4

Paraffin embedded mouse and human tissues were cut in 3 μm thick sections using the Hyrax M55 microtome (Zeiss). Tissue sections were incubated for one hour at 60 °C in order to melt paraffin, deparaffinized by incubating two times in xylene for 5 min and rehydrated in a descending alcohol series (100 %, 90 %, 80 % and 70 % (v/v)) for 1 min. To block endogenous protease activity and to permeabilize sections for nuclear staining they were incubated in a methanol/hydrogen peroxide (80 %/1.8 % (v/v)) solution for 20 min. Tissue sections were rinsed in Milli-Q^®^ water and heat-induced antigen retrieval was performed in citrate buffer pH 6 using a decloaking chamber (Biocare Medical). After washing with TBST unspecific binding sites were blocked for 30 min with Rodent Block M (Biocare Medical). The slides were washed again in TBST and incubated with anti-PSME4 antibody (sc-135512, Santa Cruz) diluted in Antibody Diluent (DAKO) for 1 h at RT. After extensive washing in TBST sections were incubated with MACH 2 Rabbit AP-Polymer (Biocare Medical) for 30 min at RT. Sections were rinsed again in TBST and incubated in Vulcan Fast Red AP substrate solution (Biocare Medical) for 10 min. Tissue sections were washed in TBST and MilliQ^®^ water and hematoxylin counterstaining (Carl Roth) was performed to visualize nuclei. After repeated washing in TBST, sections were dehydrated in ethanol and xylene and mounted using Entellan mounting medium (Merck Millipore). Slides were imaged using the MIRAX scanning system (Zeiss).

Stainings were analyzed by an expert clinical pathologist blinded to the sample identity. Tumor staging was done as described previously (Klotz et al., 2019). Semiquantitative scores for PSME4 expression were obtained by defining the percentage of PSME4 positively stained tumor areas multiplied by the intensity of staining as graded between 1 (weak) to 3 (strong). Scores were dichotomized into high (>80) and low (<80) expressing tumors with the score 80 representing the median of all samples and PSME4 expression scores were correlated with survival of the patients for this cohort.

### q-PCR analysis

RNA was extracted using Direct-zol™ RNA MiniPrep R2051 (ZYMO research) mRNA levels were ascertained by RT using High-Capacity cDNA Reverse Transcription Kit (ThermoFisher) real time quantitative PCR using sybr-green (Kapa Biosystems) and the StepOnePlus™ Real-Time PCR Systems (Life Technologies) using the following primers:

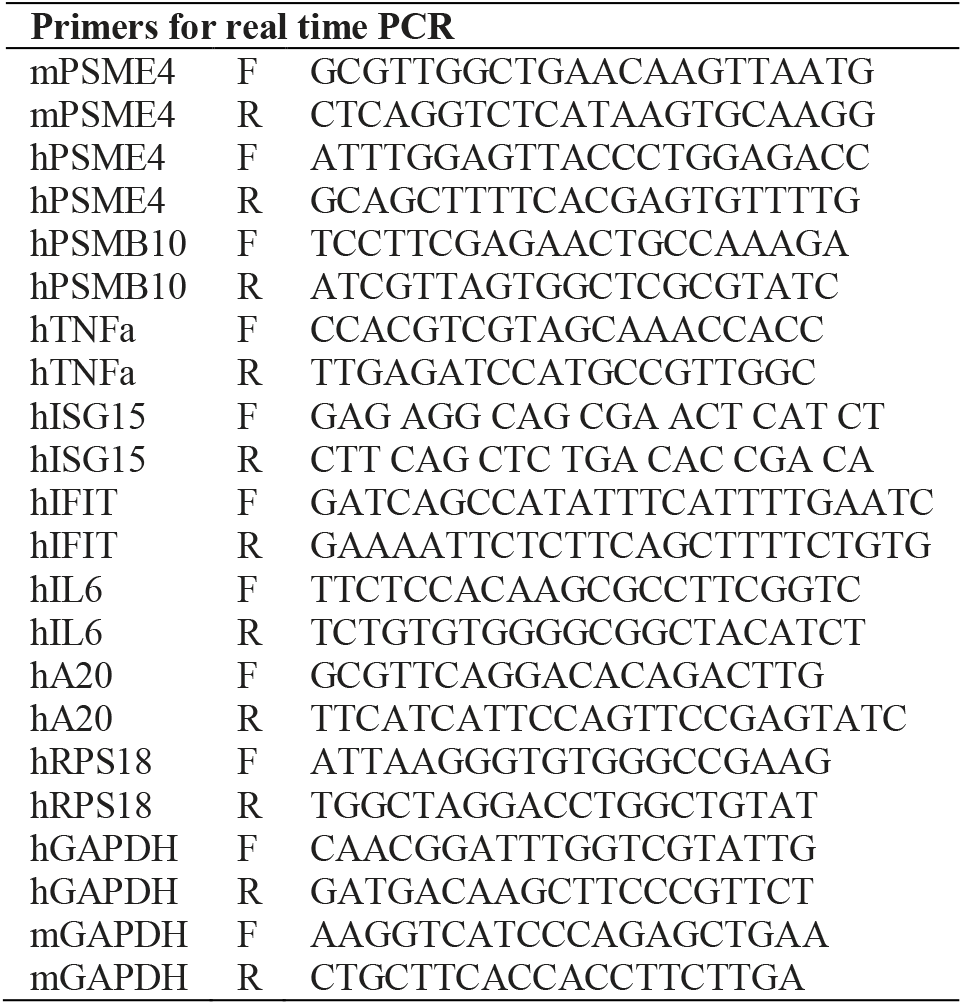

All values were normalized to the mRNA abundance of housekeeping genes (RPS18 or GAPDH in Human; GAPDH in Mouse). Each primers pair was calibrated using the Absolute Quantification program with increasing concentrations of cDNA.

### EVOC preparation

Resected tumors were treated for 3 days, in an Ex Vivo Organ Culture (EVOC) with ICI and responders were classified based on their IFNγ expression as described (Kamer et al.). In short, IFNγ levels were tested following treatment with Durvalumab alone (αPD-L1), or combined with Ipilimumab (αCTLA-4) in the EVOCs. In cases where IFNγ was significantly enriched by at least 1.9-fold induction the EVOCs were designated as ‘responder’, while the rest were termed ‘non-responder’. This was recently shown to be a correlate for responsiveness to ICI in patients (Kamer et al.).

### Orthotopic lung cancer model

Mouse experiments were conducted according to approved experimental procedures (Approval numbers 04400520-2 and 04990620-1). Male C57Bl/6J mice (Envigo, Israel) or RAG1 -/- (Jackson Laboratory) at the age of 8-10 weeks were injected i.v. with 2×10^5 cells in 200 μL PBS (Biological Industries, Israel). Mice were observed for adverse effects and weights noted twice a week. Peripheral blood was collected from the tail vein. At the end of experiment, mice were sacrificed by CO2 asphyxiation and tissues collected to cold PBS.

### Tissue processing and flow cytometry staining

Peripheral blood was washed and red blood cells removed by ACK lysis buffer (150 mM NH4Cl, 10 mM KHCO3, 0.1 mM EDTA in 0.1x PBS-/-) and washed in flow cytometry buffer (PBS-/-, 0.5% BSA, 2mM EDTA).

Spleens and lungs were weighted before further processing. The left lung lobe was used for histological analysis of tumor development (hematoxylin and eosin) after fixation in 4% formaldehyde (Biolabs, Israel). Right lung lobes were minced and digested with 5 mL digestion buffer: Collagenase 4 (200U/mL, Worthington Biochemicals), DNase I (100 ug/mL, Sigma Aldrich) in PBS++ supplemented with 2 mM CaCl2 for 20 minutes at 37°C under shaking. Single cell suspensions of spleen and lungs were obtained by straining cells through 100 um strainers and washed in cold flow cytometry buffer. Red blood cells were removed by ACK lysis buffer, washed with flow cytometry buffer and strained again.

For intracellular cytokine and transcription factor stainings, cells were incubated for 4 hours with Brefeldin A (Biolegend) and Monensin (Sigma Aldrich). First, viability staining was done with Zombie Aqua (Biolegend) according to the protocols. Cell surface staining was performed in flow cytometry buffer in 100 μL at appropriate dilutions. When needed, cells were fixed and permeabilized with the FoxP3 transcription factor kit from eBioscience (Thermo Fisher Scientific) and stained for FoxP3 and IFNg according to the manufacturers protocol. Samples were acquired on an Attune Nxt with Autosampler and analyzed in FlowJo (V10.7.1, Becton, Dickinson and Company). Exemplary gating strategies are provided Figures S10, S20 and S22)

### Cytotoxicity assay

10,000 KP1.9 WT cells were stained with CFSE (5uM; BioLegend), at a cell concentration of 1×106/ml), plated onto a 96-well plate and allowed to attach overnight. The following day, splenocytes were isolated from naïve mice, or from mice bearing KP1.9-WT, -PSME4 KD, or - PSME4 OE tumors. Red blood cells were lysed using ACK buffer, cells were washed in PBS, and 100,000 splenocytes were plated on top of the tumor cells in RPMI medium + beta-mercaptoethanol (50uM). Co-culture was maintained for 7 days. On day 7, cell supernatants were collected onto a new 96-well plate, and the attached tumor cells were detached with Trypsin containing Propidium Iodide (PI). After cell detachment, the supernatant was used to resuspend the detached cells, and the entire sample was acquired.

### Cytokine and chemokine secretion

Tumor microenvironment; 10,000 KP1.9 WT cells were plated onto a 96-well plate and allowed to attach overnight. The following day, splenocytes were isolated from naïve mice, or from mice bearing KP1.9-WT, -PSME4 KD, or -PSME4 OE tumors. Red blood cells were lysed using ACK buffer, cells were washed in PBS, and 100,000 splenocytes were plated on top of the tumor cells in RPMI medium + beta-mercaptoethanol (50uM). Co-culture was maintained for 48h or 7 days. The medium was collected at 48h and 7days by centrifugation. Cytokines and chemokines concentration in the tumor microenvironment were determined by 36-Plex Mouse ProcartaPlex (ThermoFisher, EPX360–26092–901) as per manufacturer’s instructions. Samples were acquired on MAGPIX System (Luminex) and analysed by MILLIPLEX Analyst V5.1 Flex.

Tumor cells secretion; 10,000 KP1.9 shCtrl or KP1.9 PSME4 KD cells were plated onto a 96-well plate and allowed to attach overnight. The following day cells were treated with mTNFα and mIFNγ mix for 24h in the last 6h ONX-0914 (1µM) (APExBIO, A4011) was added to desired wells. Cells were washed twice and incubated for 24h in fresh medium. The medium was collected flowed be centrifugation. Cytokines and chemokines secretion were determined by 36-Plex Mouse ProcartaPlex (ThermoFisher, EPX360–26092–901) as per manufacturer’s instructions. Samples were acquired on MAGPIX System (Luminex) and analysed by MILLIPLEX Analyst V5.1 Flex.

### Single cells RNA-Seq of CD45+ cells from lung tissues

#### CD45+ cell isolation from lung tissues

C57Bl/6J mice were injected with KP1.9 shCtrl or shPSME4 cells (0.25×10^6 cells, i.v.). After 21 days, lungs were perfused manually with 3 mL cold PBS w/o Mg/Ca through the right ventricle of the heart and harvested (4x shCtrl, 4x shPSME4) into cold PBS. The lung tissue was cut into pieces and processed using a GentleMACS Octo Dissociator (program 37C_m_LDK_1, Miltenyi Biotec) with the mouse lung dissociation kit (130-095-927, Miltenyi Biotec) according to the manufacturer’s protocol. After the run, cells were strained through a 70 um mesh and washed with FACS buffer (0.5% BSA, 2 mM EDTA in PBS without Mg/Ca). Red blood cells were lysed with ACK buffer for 4 minutes, the reaction stopped by addition of cold FACS buffer. Finally, cells were resuspended in FACS buffer with FC block (anti-CD16/CD32, 1:500, Biolegend 101330) and incubated for 5 minutes. Lung cells from shCtrl or shPSME4 were stained with four differently labeled CD45 antibodies (APC, PE/Cy7, FITC, violetFluor 450) for 30 minutes at 4C. Cells were washed with FACS buffer twice, and the shCtrl or shPSME4 samples unified for parallel sorting of the four lung populations. Propidium iodide was added fresh (1 ug/mL) to exclude dead cells, and cells were sorted on BD FACSAria III and BD SORP FACSAria II sorters running on BD FACSDiva software (V8.0.1) using a 85 um nozzle. A gating strategy is provided in the supplementary single cell documentation. Finally, cells were washed twice (PBS w/o Mg/Ca, 0.04% BSA) and counted on a Neubauer chamber with trypan blue, and three samples from each condition were chosen (high cell count and viability >90%) for further processing.

| Target | clone | color | Company | lot | Cat# | dilution |
| --- | --- | --- | --- | --- | --- | --- |
| CD45.2 | 104 | APC | eBioscience | 2265360 | 17-0454-82 | 300 |
| CD45.2 | 104 | Pe/Cy7 | eBioscience | 4330432 | 25-0454-82 | 300 |
| CD45.2 | 104 | FITC | Biolegend | B250186 | 109806 | 300 |
| CD45 | 30-F11 | violetFluor 450 | Tonbo | C0451092017753 | 75-0451 | 300 |

#### Single cell RNA-seq using Chromium 10x genomics platform

Single cell RNA-seq libraries were prepared at the Crown Genomics institute of the Nancy and Stephen Grand Israel National Center for Personalized Medicine, Weizmann Institute of Science. Cells were counted and diluted to a final concentration of approximately 1000 cells/ L in PBS supplemented with 0.04% BSA. Cellular suspension was loaded onto Next GEM Chip K targeting 6000 cells and then ran on a Chromium Controller instrument to generate GEM emulsion (10x Genomics). Single-cell gene expression libraries as well as single cell V(D)J libraries were generated according to the manufacturer’s protocol using the Chromium Next GEM Single Cell 5’ Reagent Kits v2 (Dual Index) workflow. Final libraries were quantified using NEBNext Library Quant Kit for Illumina (NEB) and high sensitivity D1000 TapeStation (Agilent). Libraries were pooled according to targeted cell number, aiming for at least 5,000 reads per cell for V(D)J libraries and 20,000 reads per cell for gene expression libraries. Pooled libraries were sequenced on a NovaSeq 6000 instrument using an SP 200 cycles reagent kit (Illumina).

#### Raw reads processing

Raw sequencing bcl files were converted to fastq files, aligned to GRCm38 mouse genome, unique molecular identifiers (UMIs) were quantified using Cell Ranger suite (v 6.0.0). V(D)J libraries were mapped to GRCm38 VDJ reference provided by 10x Genomics (v 5.0.0), and clones were identified, counted and summarized with Cell Ranger suite (v 6.0.0). Filtered count matrices were imported into R for further processing.

#### Clustering and cell type identification

In gene expression libraries low quality cells with more than 10% reads mapping to mitochondrial genome were removed. The Seurat R package (version 4.0.3) was used to normalize and scale expression values for total UMI counts per cell (Stuart et al., 2019). Due to complexity and variability in the immune cell population we performed stepwise clustering (Grün et al., 2015). A detailed clustering strategy is shown in supplementary single cell documentation. First, 2000 highly variable genes were identified using vst method. Dimensionality reduction was done with principal component analysis, with the first 30 principal components used for nearest neighbor graph construction and clustering. For each cluster based on their marker expression we identified whether it belongs to one of the groups: non-immune, B cells, T cells, ILCs, macrophages and DCs, cycling cells and other. Subsequently we performed clustering within each group individually as described above, for each cluster we identified markers (Seurat FindMarkers function) and removed clusters containing doublets, i.e. clusters that did not have any uniquely expressed genes and contained markers specific for of at least 2 abundant clusters. For example, clusters with high expression of Ncr1 (NK cell marker) and Cd79b (B cell marker) was considered doublets.

Finally, for each cluster we identified markers (Seurat FindMarkers function) and annotated them through comparison with markers in literature and ImmGen database(Heng et al., 2008; Maier et al., 2020; Rosser and Mauri, 2015)

#### Functional analysis of cell populations

Gene Ontology analysis was performed using g:Profiler2(Raudvere et al., 2019) with default settings, and multiple hypothesis testing adjustment using all mouse genes as background control. The log10 FDR-adjusted P values were plotted as barplots.

#### Differential expression analysis

To compare gene expression of each cell type cluster between conditions, we calculated pseudobulk by adding reads from all cells within each cluster in a sample. We then used DESeq2 with default parameters to determine differential expression(Love et al., 2014).

#### Statistics and reproducibility

All experiments were performed in three individual replicates unless otherwise mentioned. For each experiment, all compared conditions were analyzed by MS at the same time. For the NSCLC study, nine samples from tumors and adjacent tissue were analyzed, one was excluded for poor technical quality. The samples were processed independently and analyzed by MS at the same time to maintain comparability across samples and decrease batch effects.

#### Ethics Statement

Human lung tissues obtained from patients surgically treated for lung cancer were provided by the Asklepios Biobank for Lung Disease, Gauting, Germany or obtained from the Israel National Biobank for Research. Samples were obtained under the approval of the Ministry of Health IRB approval for the Israel National Biobank for Research, protocol no. 118-2018, or the ethics committee of the Ludwig-Maximilians University Munich according to national and international guidelines (project number 333-10).

For the ex-vivo organoid cultures, all patients provided written informed consent for the use of blood samples and tumor specimens for research. This study was approved by the Sheba Medical Center ethics committee.

## Code availability

All the code used is available from the corresponding author upon reasonable request.

## Acknowledgments

We thank the members of the Merbl lab as well as Ayelet Erez and Bareket Dassa for discussion and critical reading of the manuscript. We would like to thank Alfred Zippelius for KP1.9 cells.

Y.M. is supported by the European Research Council (ERC) under the European Union’s Horizon 2020 research and innovation program (grant agreement No 677748); The I-CORE Program of the Planning and Budgeting Committee and The Israel Science Foundation (Grant No. 1775/12) and the Israeli Science Foundation (Grant No. 2109/18); The Gruber Peter & Patricia award. M.D.S. is supported by Marie Sklodowska-Curie Individual Fellowship (Horizon 2020 Grant No. GAP-845066). Y.M. the incumbent of the Leonard and Carol Berall Career Development Chair. This manuscript was edited at Life Science Editors.

## Author Contributions

AJ, MDS and YM conceived, designed and interpreted experiments and wrote the manuscript. AL-E and SM provided feedback on the manuscript. MDS, MR, AU, HW-L, AE-L proteomics preparation and SM for in vitro work. MPK and IC in vivo and immunofluorescence. AAK, MPK and AJ performed and analyzed the scRNAseq experiment. AJ and KL analyzed the ICI cohort. IK, EBD and OZ conducted the EVOC and immunohistochemistry. VW, ESP, ML, and IK analysis of German patient cohort. AK managed NSCLC Israeli cohort. MA and MDS immunopeptidomics. YL Mass spectrometry. ATR and PDF purified recombinant PSME4. MDS performed all other cellular and biochemical assays. AJ performed all other bioinformatics. JB, YS,EE, NF, CS SM, and YM funded and supervised the work of respective group members and YM supervised the study.

## References

Ahmed, A.M. (2019). The Dual Role of Oxidative Stress in Lung Cancer. In Oxidative Stress in Lung Diseases, (Springer Singapore), pp. 99–113.

Van Allen, E.M., Miao, D., Schilling, B., Shukla, S.A., Blank, C., Zimmer, L., Sucker, A., Hillen, U., Foppen, M.H.G., Goldinger, S.M., et al. (2015). Genomic correlates of response to CTLA-4 blockade in metastatic melanoma. Science (80-.). 350, 207–211.

Antoniou, A.N., Lenart, I., Guiliano, D.B., and Powis, S.J. (2012). Antigen Processing and Presentation by MHC Class I, II, and Nonclassical Molecules. In Vaccinology: Principles and Practice, (Department of Infection and Immunity/Centre for Rheumatology, University College London, London, United Kingdom: Wiley-Blackwell), pp. 29–46.

Arima, K., Kinoshita, A., Mishima, H., Kanazawa, N., Kaneko, T., Mizushima, T., Ichinose, K., Nakamura, H., Tsujino, A., Kawakami, A., et al. (2011). Proteasome assembly defect due to a proteasome subunit beta type 8 (PSMB8) mutation causes the autoinflammatory disorder, Nakajo-Nishimura syndrome. Proc. Natl. Acad. Sci. U. S. A. 108, 14914–14919.

Avalle, L., Pensa, S., Regis, G., Novelli, F., and Poli, V. (2012). STAT1 and STAT3 in tumorigenesis. JAK-STAT 1, 65–72.

Ayers, M., Lunceford, J., Nebozhyn, M., Murphy, E., Loboda, A., Kaufman, D.R., Albright, A., Cheng, J.D., Kang, S.P., Shankaran, V., et al. (2017). IFN-γ-related mRNA profile predicts clinical response to PD-1 blockade. J. Clin. Invest. 127, 2930–2940.

Baba, T., and Mukaida, N. (2014). Role of macrophage inflammatory protein (MIP)-1α/CCL3 in leukemogenesis. Mol. Cell. Oncol. 1.

Blickwedehl, J., Olejniczak, S., Cummings, R., Sarvaiya, N., Mantilla, A., Chanan-Khan, A., Pandita, T.K., Schmidt, M., Thompson, C.B., and Bangia, N. (2012). The proteasome activator PA200 regulates tumor cell responsiveness to glutamine and resistance to ionizing radiation. Mol. Cancer Res. 10, 937–944.

Chong, C., Marino, F., Pak, H., Racle, J., Daniel, R.T., Müller, M., Gfeller, D., Coukos, G., and Bassani-Sternberg, M. (2018). High-throughput and Sensitive Immunopeptidomics Platform Reveals Profound Interferonγ-Mediated Remodeling of the Human Leukocyte Antigen (HLA) Ligandome. Mol. Cell. Proteomics 17, 533–548.

Collins, G.A., and Goldberg, A.L. (2017). The Logic of the 26S Proteasome. Cell 169, 792–806.

Coux, O., Zieba, B.A., and Meiners, S. (2020). The proteasome system in health and disease. In Advances in Experimental Medicine and Biology, (Springer), pp. 55–100.

Driscoll, J., Brown, M.G., Finley, D., and Monaco, J.J. (1993). MHC-linked LMP gene products specifically alter peptidase activities of the proteasome. Nature 365, 262–264.

Fabre, B., Lambour, T., Garrigues, L., Ducoux-Petit, M., Amalric, F., Monsarrat, B., Burlet-Schiltz, O., and Bousquet-Dubouch, M.P. (2014). Label-free quantitative proteomics reveals the dynamics of proteasome complexes composition and stoichiometry in a wide range of human cell lines. J. Proteome Res. 13, 3027–3037.

Fabre, B., Lambour, T., Garrigues, L., Amalric, F., Vigneron, N., Menneteau, T., Stella, A., Monsarrat, B., Van den Eynde, B., Burlet-Schiltz, O., et al. (2015). Deciphering preferential interactions within supramolecular protein complexes: the proteasome case. Mol. Syst. Biol. 11, 771.

Ferrington, D.A., and Gregerson, D.S. (2012). Immunoproteasomes: structure, function, and antigen presentation. Prog. Mol. Biol. Transl. Sci. 109, 75–112.

Gaczynska, M., Rock, K.L., and Goldberg, A.L. (1993). |[gamma]|-Interferon and expression of MHC genes regulate peptide hydrolysis by proteasomes., Publ. Online 16 Sept. 1993; | Doi10.1038/365264a0 365, 264.

Gillette, M.A., Satpathy, S., Cao, S., Dhanasekaran, S.M., Vasaikar, S. V., Krug, K., Petralia, F., Li, Y., Liang, W.W., Reva, B., et al. (2020). Proteogenomic Characterization Reveals Therapeutic Vulnerabilities in Lung Adenocarcinoma. Cell 182, 200–225.e35.

Grün, D., Lyubimova, A., Kester, L., Wiebrands, K., Basak, O., Sasaki, N., Clevers, H., and Van Oudenaarden, A. (2015). Single-cell messenger RNA sequencing reveals rare intestinal cell types. Nature 525, 251–255.

Harel, M., Ortenberg, R., Varanasi, S.K., Mangalhara, K.C., Mardamshina, M., Markovits, E., Baruch, E.N., Tripple, V., Arama-Chayoth, M., Greenberg, E., et al. (2019). Proteomics of Melanoma Response to Immunotherapy Reveals Mitochondrial Dependence. Cell 179, 236–250.e18.

Heng, T.S.P., Painter, M.W., Elpek, K., Lukacs-Kornek, V., Mauermann, N., Turley, S.J., Koller, D., Kim, F.S., Wagers, A.J., Asinovski, N., et al. (2008). The immunological genome project: Networks of gene expression in immune cells. Nat. Immunol. 9, 1091–1094.

Hugo, W., Zaretsky, J.M., Sun, L., Song, C., Moreno, B.H., Hu-Lieskovan, S., Berent-Maoz, B., Pang, J., Chmielowski, B., Cherry, G., et al. (2016). Genomic and Transcriptomic Features of Response to Anti-PD-1 Therapy in Metastatic Melanoma. Cell 165, 35–44.

Javitt, A., and Merbl, Y. (2019). Global views of proteasome-mediated degradation by mass spectrometry. Expert Rev. Proteomics 16.

Javitt, A., Barnea, E., Kramer, M.P., Wolf-Levy, H., Levin, Y., Admon, A., and Merbl, Y. (2019a). Pro-inflammatory cytokines alter the immunopeptidome landscape by modulation of HLA-B expression. Front. Immunol. 10.

Javitt, A., Barnea, E., Kramer, M.P.M.P., Wolf-Levy, H., Levin, Y., Admon, A., and Merbl, Y. (2019b). Pro-inflammatory cytokines alter the immunopeptidome landscape by modulation of HLA-B expression. Front. Immunol. 10, 141.

Jiang, T.-X., Ma, S., Han, X., Luo, Z.-Y., Zhu, Q.-Q., Chiba, T., Xie, W., Lin, K., and Qiu, X.-B. (2021). Proteasome activator PA200 maintains stability of histone marks during transcription and aging. Theranostics 11, 1458.

Johnson, D.E., O’Keefe, R.A., and Grandis, J.R. (2018). Targeting the IL-6/JAK/STAT3 signalling axis in cancer. Nat. Rev. Clin. Oncol. 15, 234–248.

Kalaora, S., Lee, J.S., Barnea, E., Levy, R., Greenberg, P., Alon, M., Yagel, G., Bar Eli, G., Oren, R., Peri, A., et al. (2020). Immunoproteasome expression is associated with better prognosis and response to checkpoint therapies in melanoma. Nat. Commun. 11.

Kamer, I., Bab-Dinitz, E., Zadok, O., Ofek, E., Gottfried, T., Daniel-Meshulam, I., Hout-Siloni, G., Nun, A. Ben, Barshack, I., Onn, A., et al. Immunotherapy response modeling by ex-vivo organ culture for lung cancer. Rev.

Kamer, I., Bab-Dinitz, E., Zadok, O., Ofek, E., Gottfried, T., Daniel-Meshulam, I., Hout-Siloni, G., Ben Nun, A., Barshack, I., Onn, A., et al. (2021). Immunotherapy response modeling by ex-vivo organ culture for lung cancer. Cancer Immunol. Immunother.

Kammerl, I.E., and Meiners, S. (2016). Proteasome function shapes innate and adaptive immune responses. Am. J. Physiol. - Lung Cell. Mol. Physiol. 311.

Kammerl, I.E., Caniard, A., Merl-Pham, J., Ben-Nissan, G., Mayr, C.H., Mossina, A., Geerlof, A., Eickelberg, O., Hauck, S.M., Sharon, M., et al. (2019). Dissecting the molecular effects of cigarette smoke on proteasome function. J. Proteomics 193, 1–9.

Khor, B., Bredemeyer, A.L., Huang, C.-Y., Turnbull, I.R., Evans, R., Maggi, L.B., White, J.M., Walker, L.M., Carnes, K., Hess, R.A., et al. (2006). Proteasome activator PA200 is required for normal spermatogenesis. Mol. Cell. Biol. 26, 2999–3007.

Kim, J.Y., Choi, J.K., and Jung, H. (2020). Genome-wide methylation patterns predict clinical benefit of immunotherapy in lung cancer. Clin. Epigenetics 2020 121 12, 1–10.

Kimura, A., and Kishimoto, T. (2010). IL-6: Regulator of Treg/Th17 balance. Eur. J. Immunol. 40, 1830–1835.

Kitamura, A., Maekawa, Y., Uehara, H., Izumi, K., Kawachi, I., Nishizawa, M., Toyoshima, Y., Takahashi, H., Standley, D.M., Tanaka, K., et al. (2011). A mutation in the immunoproteasome subunit PSMB8 causes autoinflammation and lipodystrophy in humans. J. Clin. Invest. 121, 4150–4160.

Klotz, L., Y, C., M, L., A, P.-C., A, S., I, K., ME, E., I, L., A, M.-H., KAM, A., et al. (2019). Comprehensive clinical profiling of the Gauting locoregional lung adenocarcinoma donors. Cancer Med. 8, 1486–1499.

Knochelmann, H.M., Dwyer, C.J., Bailey, S.R., Amaya, S.M., Elston, D.M., Mazza-McCrann, J.M., and Paulos, C.M. (2018). When worlds collide: Th17 and Treg cells in cancer and autoimmunity. Cell. Mol. Immunol. 15, 458–469.

Lavin, Y., Kobayashi, S., Leader, A., and Rahman, A. (2017). Innate Immune Landscape in Early Lung Adenocarcinoma by Paired Single-Cell Analyses In Brief Comparing single tumor cells with adjacent normal tissue and blood from patients with lung adenocarcinoma charts early changes in tumor immunity and provides insi. Cell 169, 750–757.e15.

Li, S., Wu, J., Zhu, S., Liu, Y.J., and Chen, J. (2017). Disease-associated plasmacytoid dendritic cells. Front. Immunol. 8, 1268.

Litchfield, K., Reading, J.L., Puttick, C., Thakkar, K., Abbosh, C., Bentham, R., Watkins, T.B.K., Rosenthal, R., Biswas, D., Rowan, A., et al. (2021). Meta-analysis of tumor-and T cell-intrinsic mechanisms of sensitization to checkpoint inhibition. Cell 184, 596–614.e14.

Liu, Y., Ramot, Y., Torrelo, A., Paller, A.S., Si, N., Babay, S., Kim, P.W., Sheikh, A., Lee, C.C.R., Chen, Y., et al. (2012). Mutations in proteasome subunit β type 8 cause chronic atypical neutrophilic dermatosis with lipodystrophy and elevated temperature with evidence of genetic and phenotypic heterogeneity. Arthritis Rheum. 64, 895–907.

Livneh, I., Cohen-Kaplan, V., Cohen-Rosenzweig, C., Avni, N., and Ciechanover, A. (2016). The life cycle of the 26S proteasome: from birth, through regulation and function, and onto its death. Cell Res. 26, 869–885.

Love, M.I., Huber, W., and Anders, S. (2014). Moderated estimation of fold change and dispersion for RNA-seq data with DESeq2. Genome Biol. 15, 550.

M. Cantin A., and V. Richter M. (2012). Cigarette Smoke-Induced Proteostasis Imbalance in Obstructive Lung Diseases. Curr. Mol. Med. 12, 836–849.

Maier, B., Leader, A.M., Chen, S.T., Tung, N., Chang, C., LeBerichel, J., Chudnovskiy, A., Maskey, S., Walker, L., Finnigan, J.P., et al. (2020). A conserved dendritic-cell regulatory program limits antitumour immunity. Nature 580, 257–262.

Mandemaker, I.K., Geijer, M.E., Kik, I., Bezstarosti, K., Rijkers, E., Raams, A., Janssens, R.C., Lans, H., Hoeijmakers, J.H., Demmers, J.A., et al. (2018a). DNA damage-induced replication stress results in PA200-proteasome-mediated degradation of acetylated histones. EMBO Rep. 19.

Mandemaker, I.K., Geijer, M.E., Kik, I., Bezstarosti, K., Rijkers, E., Raams, A., Janssens, R.C., Lans, H., Hoeijmakers, J.H., Demmers, J.A., et al. (2018b). DNA damage-induced replication stress results in PA200-proteasome-mediated degradation of acetylated histones. EMBO Rep. 19, e45566.

Mariathasan, S., Turley, S.J., Nickles, D., Castiglioni, A., Yuen, K., Wang, Y., Kadel, E.E., Koeppen, H., Astarita, J.L., Cubas, R., et al. (2018). TGFβ attenuates tumour response to PD-L1 blockade by contributing to exclusion of T cells. Nature 554, 544–548.

McDermott, D.F., Huseni, M.A., Atkins, M.B., Motzer, R.J., Rini, B.I., Escudier, B., Fong, L., Joseph, R.W., Pal, S.K., Reeves, J.A., et al. (2018). Clinical activity and molecular correlates of response to atezolizumab alone or in combination with bevacizumab versus sunitinib in renal cell carcinoma. Nat. Med. 24, 749–757.

Morozov, A. V., and Karpov, V.L. (2019). Proteasomes and several aspects of their heterogeneity relevant to cancer (Frontiers Media S.A.).

Motosugi, R., and Murata, S. (2019). Dynamic regulation of proteasome expression. Front. Mol. Biosci. 6.

Muchamuel, T., Basler, M., Aujay, M.A., Suzuki, E., Kalim, K.W., Lauer, C., Sylvain, C., Ring, E.R., Shields, J., Jiang, J., et al. (2009). A selective inhibitor of the immunoproteasome subunit LMP7 blocks cytokine production and attenuates progression of experimental arthritis. Nat. Med. 15, 781–787.

O’Donnell, T.J., Rubinsteyn, A., and Laserson, U. (2020). MHCflurry 2.0: Improved Pan-Allele Prediction of MHC Class I-Presented Peptides by Incorporating Antigen Processing. Cell Syst. 11, 42–48.e7.

Qian, M.-X., Pang, Y., Liu, C.H., Haratake, K., Du, B.-Y., Ji, D.-Y., Wang, G.-F., Zhu, Q.-Q., Song, W., Yu, Y., et al. (2013). Acetylation-Mediated Proteasomal Degradation of Core Histones during DNA Repair and Spermatogenesis. Cell 153, 1012–1024.

Raudvere, U., Kolberg, L., Kuzmin, I., Arak, T., Adler, P., Peterson, H., and Vilo, J. (2019). G:Profiler: A web server for functional enrichment analysis and conversions of gene lists (2019 update). Nucleic Acids Res. 47, W191–W198.

Raule, M., Cerruti, F., Benaroudj, N., Migotti, R., Kikuchi, J., Bachi, A., Navon, A., Dittmar, G., and Cascio, P. (2014). PA28αβ Reduces Size and Increases Hydrophilicity of 20S Immunoproteasome Peptide Products. Chem. Biol. 21, 470–480.

Riaz, N., Havel, J.J., Makarov, V., Desrichard, A., Urba, W.J., Sims, J.S., Hodi, F.S., Martín-Algarra, S., Mandal, R., Sharfman, W.H., et al. (2017a). Tumor and Microenvironment Evolution during Immunotherapy with Nivolumab. Cell 171, 934–949.e15.

Riaz, N., Havel, J.J., Makarov, V., Desrichard, A., Urba, W.J., Sims, J.S., Hodi, F.S., Martín-Algarra, S., Mandal, R., Sharfman, W.H., et al. (2017b). Tumor and Microenvironment Evolution during Immunotherapy with Nivolumab. Cell 171, 934–949.e16.

Rock, K.L., Reits, E., and Neefjes, J. (2016). Present Yourself! By MHC Class I and MHC Class II Molecules. Trends Immunol. 37, 724–737.

Rodriguez, A., Perez-Gonzalez, A., and Nieto, A. (2007). Influenza Virus Infection Causes Specific Degradation of the Largest Subunit of Cellular RNA Polymerase II. J. Virol. 81, 5315–5324.

Rosser, E.C., and Mauri, C. (2015). Regulatory B Cells: Origin, Phenotype, and Function. Immunity 42, 607–612.

Rousseau, A., and Bertolotti, A. (2018). Regulation of proteasome assembly and activity in health and disease. Nat. Rev. Mol. Cell Biol. 19, 697–712.

Salzmann, U., Kral, S., Braun, B., Standera, S., Schmidt, M., Kloetzel, P.-M., and Sijts, A. (1999). Mutational analysis of subunit iβ2 (MECL-1) demonstrates conservation of cleavage specificity between yeast and mammalian proteasomes. FEBS Lett. 454, 11–15.

Schmidt, C., Berger, T., Groettrup, M., and Basler, M. (2018). Immunoproteasome Inhibition Impairs T and B Cell Activation by Restraining ERK Signaling and Proteostasis. Front. Immunol. 9, 2386.

Snyder, A., Makarov, V., Merghoub, T., Yuan, J., Zaretsky, J.M., Desrichard, A., Walsh, L.A., Postow, M.A., Wong, P., Ho, T.S., et al. (2014). Genetic Basis for Clinical Response to CTLA-4 Blockade in Melanoma. N. Engl. J. Med. 371, 2189–2199.

Spits, M., and Neefjes, J. (2016). Immunoproteasomes and immunotherapy—a smoking gun for lung cancer? J. Thorac. Dis. 8, E558–E563.

Spranger, S., Bao, R., and Gajewski, T.F. (2015). Melanoma-intrinsic β-catenin signalling prevents anti-tumour immunity. Nature 523, 231–235.

Srinivas, U.S., Tan, B.W.Q., Vellayappan, B.A., and Jeyasekharan, A.D. (2019). ROS and the DNA damage response in cancer. Redox Biol. 25, 101084.

Stuart, T., Butler, A., Hoffman, P., Hafemeister, C., Papalexi, E., Mauck, W.M., Hao, Y., Stoeckius, M., Smibert, P., and Satija, R. (2019). Comprehensive Integration of Single-Cell Data. Cell 177, 1888–1902.e21.

Thul, P.J., Akesson, L., Wiking, M., Mahdessian, D., Geladaki, A., Ait Blal, H., Alm, T., Asplund, A., Björk, L., Breckels, L.M., et al. (2017). A subcellular map of the human proteome. Science (80-.). 356.

Toste Rêgo, A., and da Fonseca, P.C.A. (2019). Characterization of Fully Recombinant Human 20S and 20S-PA200 Proteasome Complexes. Mol. Cell 76, 138–147.e5.

Tripathi, S.C., Peters, H.L., Taguchi, A., Katayama, H., Wang, H., Momin, A., Jolly, M.K., Celiktas, M., Rodriguez-Canales, J., Liu, H., et al. (2016). Immunoproteasome deficiency is a feature of non-small cell lung cancer with a mesenchymal phenotype and is associated with a poor outcome. Proc. Natl. Acad. Sci. U. S. A. 113, E1555–64.

Trujillo, J.A., Sweis, R.F., Bao, R., and Luke, J.J. (2018). T cell–inflamed versus Non-T cell–inflamed tumors: a conceptual framework for cancer immunotherapy drug development and combination therapy selection. Cancer Immunol. Res. 6, 990–1000.

Uhlen, M., Zhang, C., Lee, S., Sjöstedt, E., Fagerberg, L., Bidkhori, G., Benfeitas, R., Arif, M., Liu, Z., Edfors, F., et al. (2017). A pathology atlas of the human cancer transcriptome. Science (80-.). 357.

Uhlén, M., Fagerberg, L., Hallström, B.M., Lindskog, C., Oksvold, P., Mardinoglu, A., Sivertsson, Å., Kampf, C., Sjöstedt, E., Asplund, A., et al. (2015). Tissue-based map of the human proteome. Science (80-.). 347.

Ustrell, V., Hoffman, L., Pratt, G., and Rechsteiner, M. (2002). Pa200, a nuclear proteasome activator involved in DNA repair. EMBO J. 21, 3516–3525.

Vizcaíno, J.A., Csordas, A., del-Toro, N., Dianes, J.A., Griss, J., Lavidas, I., Mayer, G., Perez-Riverol, Y., Reisinger, F., Ternent, T., et al. (2016). 2016 update of the PRIDE database and its related tools. Nucleic Acids Res. 44, D447–D456.

Welk, V., Coux, O., Kleene, V., Abeza, C., Trumbach, D., Eickelberg, O., and Meiners, S. (2016). Inhibition of proteasome activity induces formation of alternative proteasome complexes. J. Biol. Chem. 291, 13147–13159.

Welk, V., Meul, T., Lukas, C., Kammerl, I.E., Mulay, S.R., Schamberger, A.C., Semren, N., Fernandez, I.E., Anders, H.J., Günther, A., et al. (2019). Proteasome activator PA200 regulates myofibroblast differentiation. Sci. Rep. 9, 1–11.

Wickam, H. (2016). ggplot2: Elegant Graphics for Data Analysis (Springer-Verlag New York).

Winter, M.B., La Greca, F., Arastu-Kapur, S., Caiazza, F., Cimermancic, P., Buchholz, T.J., Anderl, J.L., Ravalin, M., Bohn, M.F., Sali, A., et al. (2017). Immunoproteasome functions explained by divergence in cleavage specificity and regulation. Elife 6, e27364.

Wolf-Levy, H., Javitt, A., Eisenberg-Lerner, A., Kacen, A., Ulman, A., Sheban, D., Dassa, B., Fishbain-Yoskovitz, V., Carmona-Rivera, C., Kramer, M.P., et al. (2018). Revealing the cellular degradome by mass spectrometry analysis of proteasome-cleaved peptides. Nat. Biotechnol. 36, 1110–1116.

Zuguang Gu, Roland Eils, and Matthias Schlesner (2016). Complex Heatmaps Reveal Patterns and Correlations in Multidimensional Genomic Data - PubMed. Bioinformatics.

