## SupplementaryFigures1-26 for "The proteasome regulator PSME4 drives immune evasion and abrogates anti-tumor immunity in NSCLC"

Figure S1

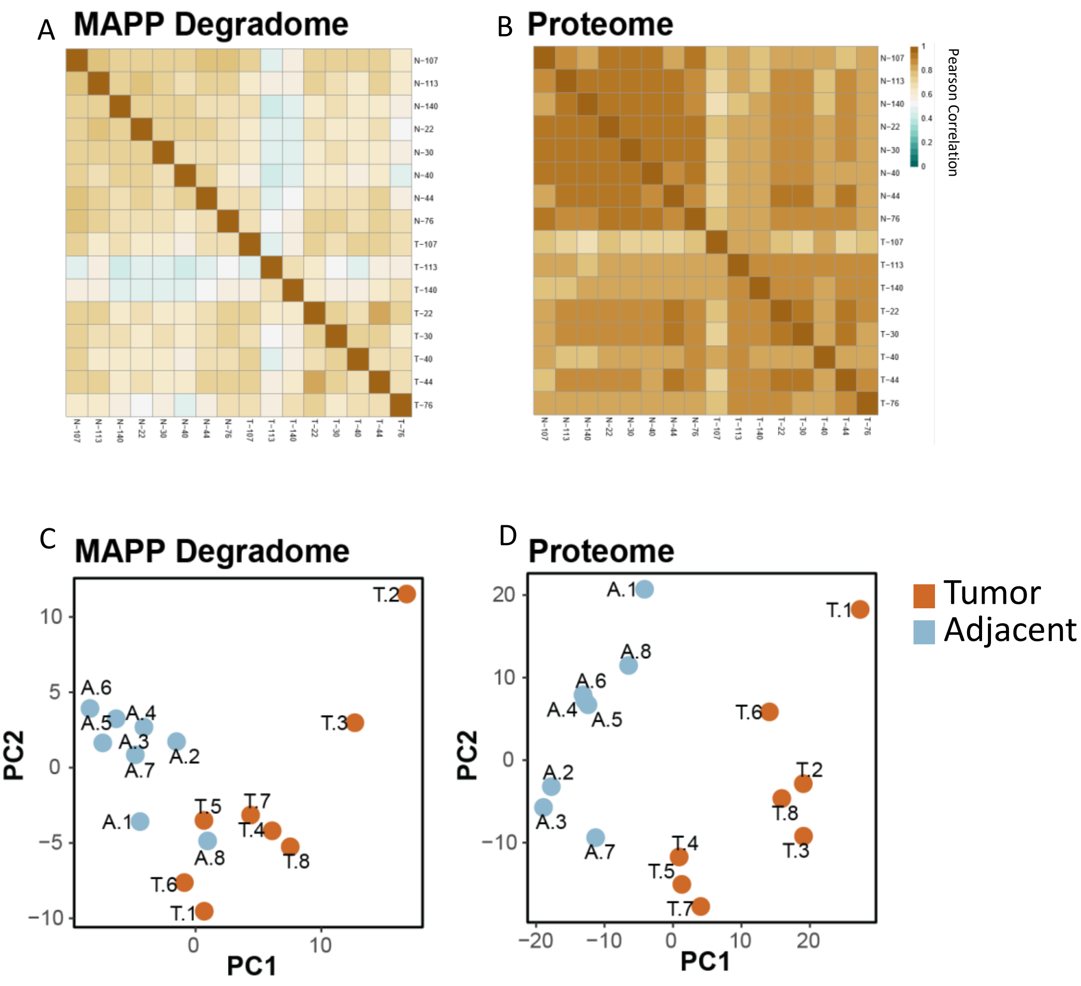
A
MAPP Degradome
B
Proteome
Pearson Correlation
C
MAPP Degradome
D
Proteome
PC1
PC2
Tumor
Adjacent

### Figure S2

A

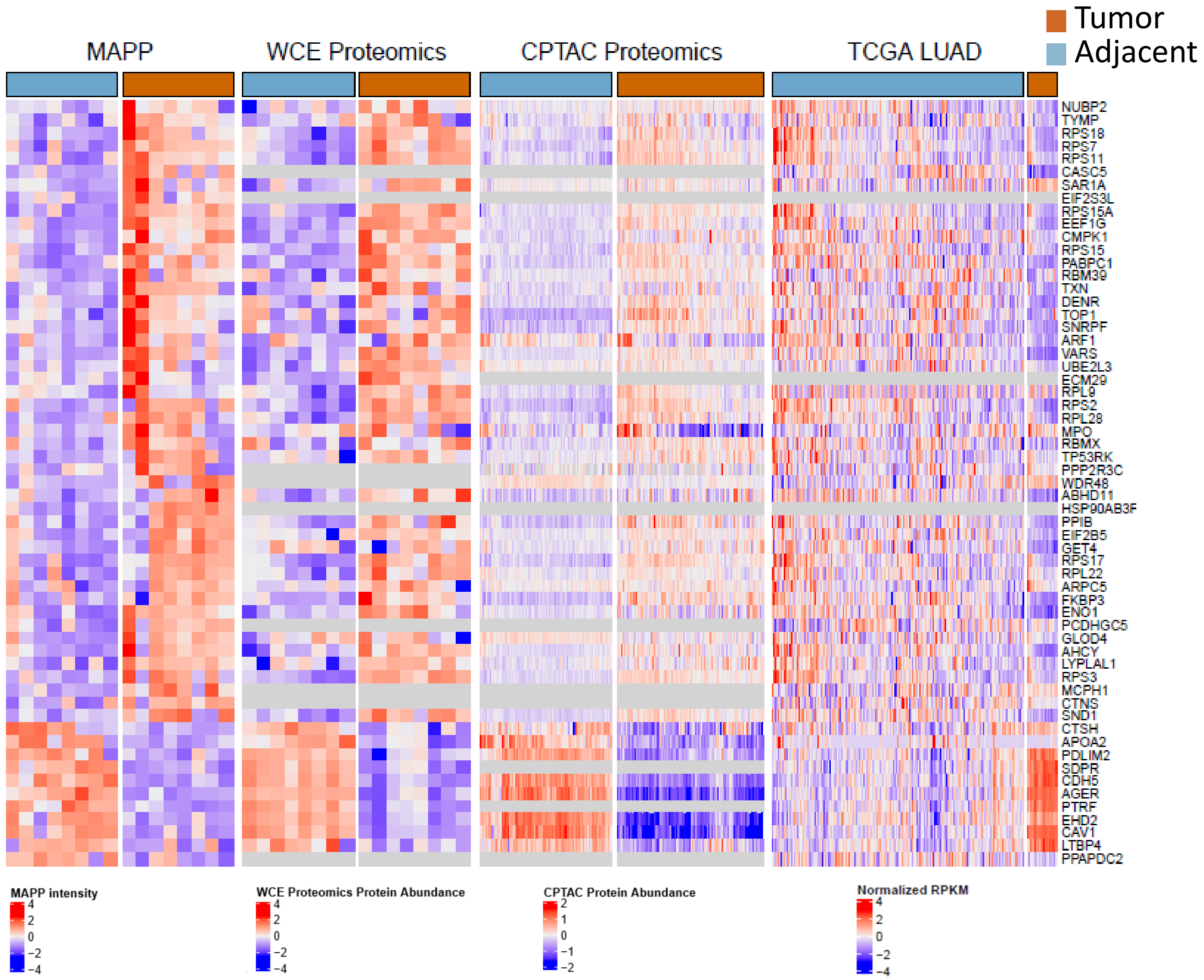
MAPP
WCE Proteomics
CPTAC Proteomics
TCGA LUAD
Tumor
Adjacent
NUBP2
TYMP
RPS18
RPS7
RPS11
CASC5
SAR1A
EIF2S3L
RPS15A
EEF1G
CMPK1
RPS15
PABPC1
RBM39
TXN
DENR
TOP1
SNRPF
ARF1
VARS
UBE2L3
ECM29
RPL9
RPS2
RPL28
MPO
RBMX
TP53RK
PPP2R3C
WDR48
ABHD11
HSP90AB3F
PPIB
EIF2B5
GET4
RPS17
RPL22
ARPC5
FKBP3
ENO1
PCDHGC5
GLOD4
AHCY
LYPLAL1
RPS3
MCPH1
CTNS
SND1
CTSH
APOA2
PDLIM2
SDPR
CDH5
AGER
PTRF
EHD2
CAV1
LTBP4
PPAPDC2
MAPP intensity
WCE Proteomics Protein Abundance
CPTAC Protein Abundance
Normalized RPKM

B

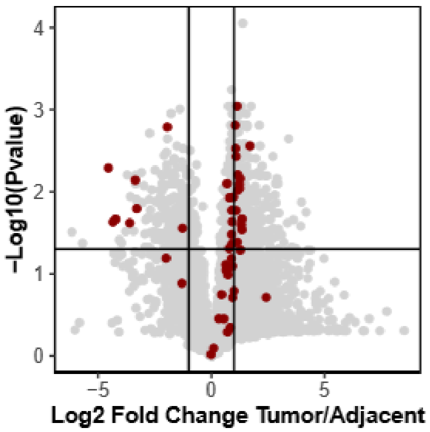
−Log10(Pvalue)
Log2 Fold Change Tumor/Adjacent

Figure S3

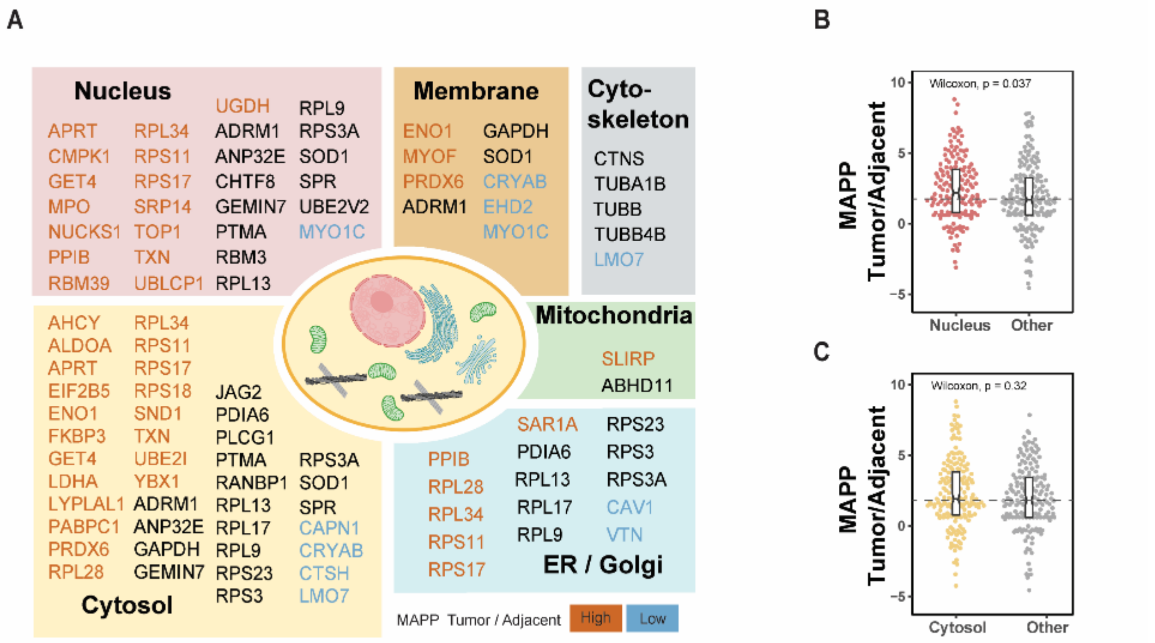
A
Nucleus
APRT RPL34 UGDH ADRM1 RPL9 RPS3A
CMPK1 RPS11 ANP32E SOD1
GET4 RPS17 CHTF8 SPR
MPO SRP14 GEMIN7 UBE2V2
NUCKS1 TOP1 PTMA MYO1C
PPIB TXN RBM3
RBM39 UBLCP1 RPL13
Membrane
ENO1 GAPDH
MYOF SOD1
PRDX6 CRYAB
ADRM1 EHD2
MYO1C
Cyto-skeleton
CTNS
TUBA1B
TUBB
TUBB4B
LMO7
Mitochondria
SLIRP
ABHD11
AHCY RPL34
ALDOA RPS11
APRT RPS17
EIF2B5 RPS18 JAG2
ENO1 SND1 PDIA6
FKBP3 TXN PLCG1
GET4 UBE2I PTMA RPS3A
LDHA YBX1 RANBP1 SOD1
LYPLAL1 ADRM1 RPL13 SPR
PABPC1 ANP32E RPL17 CAPN1
PRDX6 GAPDH RPL9 CRYAB
RPL28 GEMIN7 RPS23 CTSH
RPS3 LMO7
Cytosol
SAR1A RPS23
PPIB PDIA6 RPS3
RPL28 RPL13 RPS3A
RPL34 RPL17 CAV1
RPS11 RPL9 VTN
RPS17
ER / Golgi
MAPP Tumor / Adjacent High Low
B
Wilcoxon, p = 0.037
MAPP Tumor/Adjacent
10 5 0 −5
Nucleus Other
C
Wilcoxon, p = 0.32
MAPP Tumor/Adjacent
10 5 0 −5
Cytosol Other

### Figure S4

A

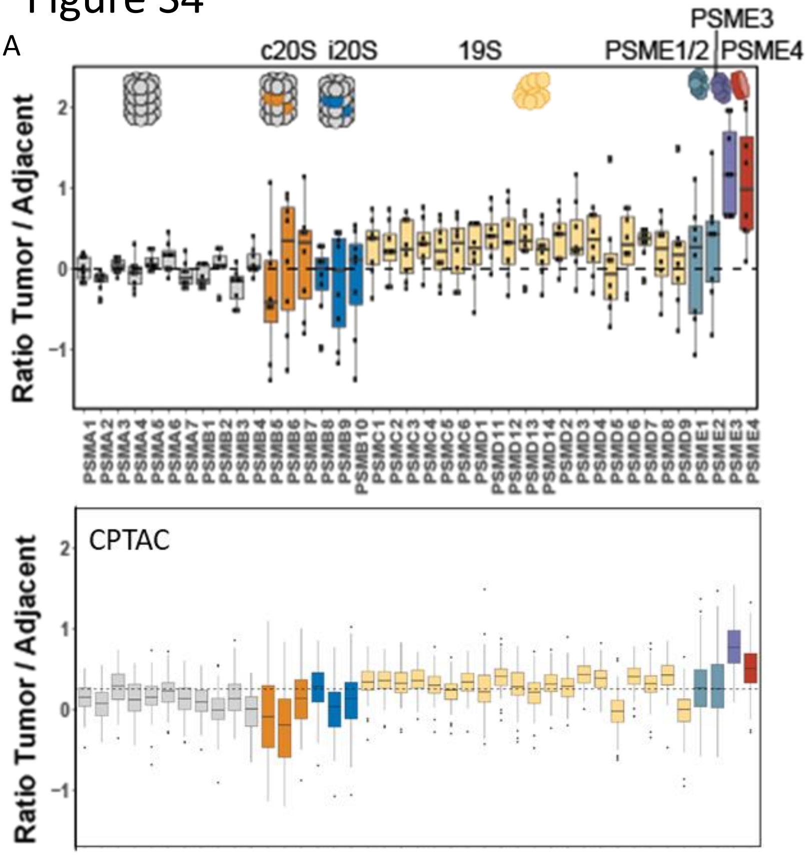


B

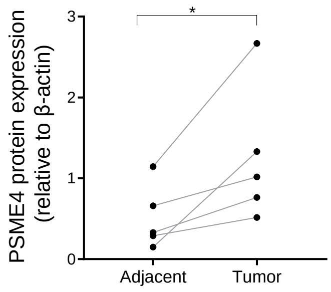


C

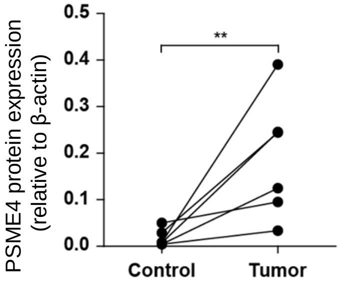


D

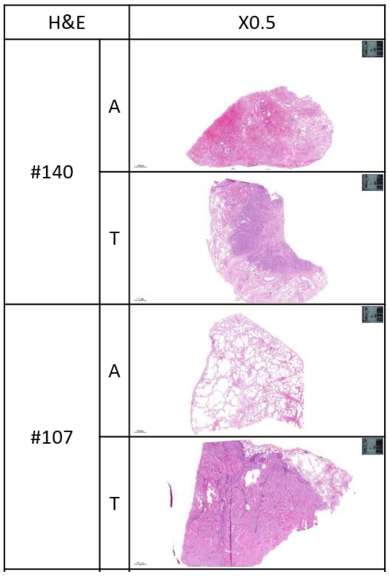


E

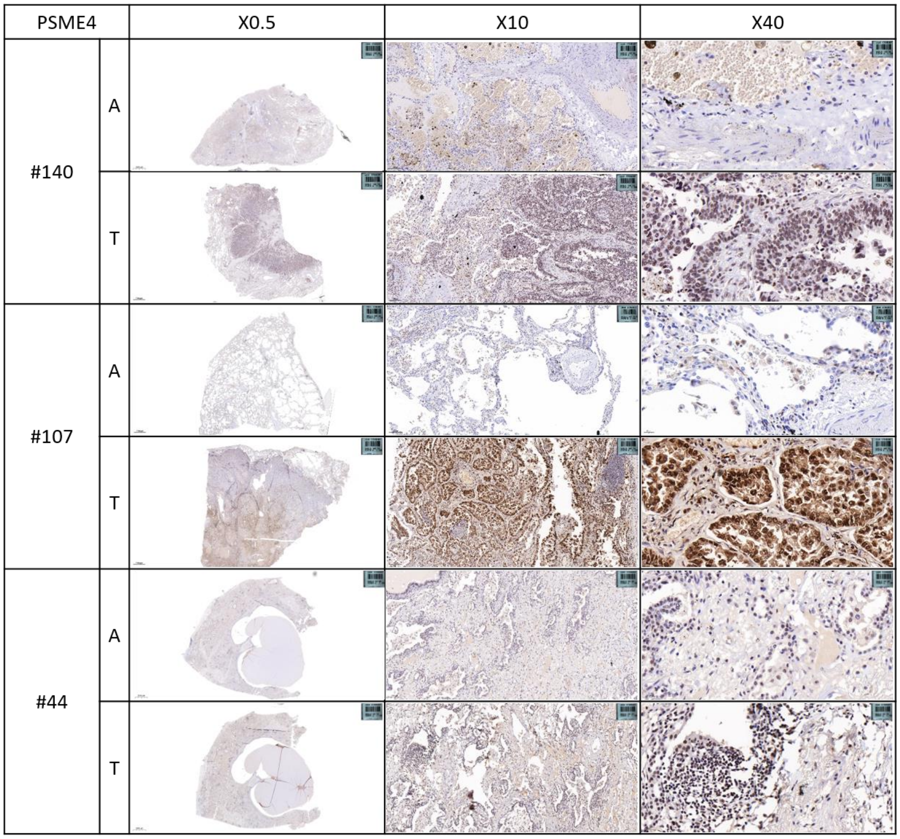

### Figure S5

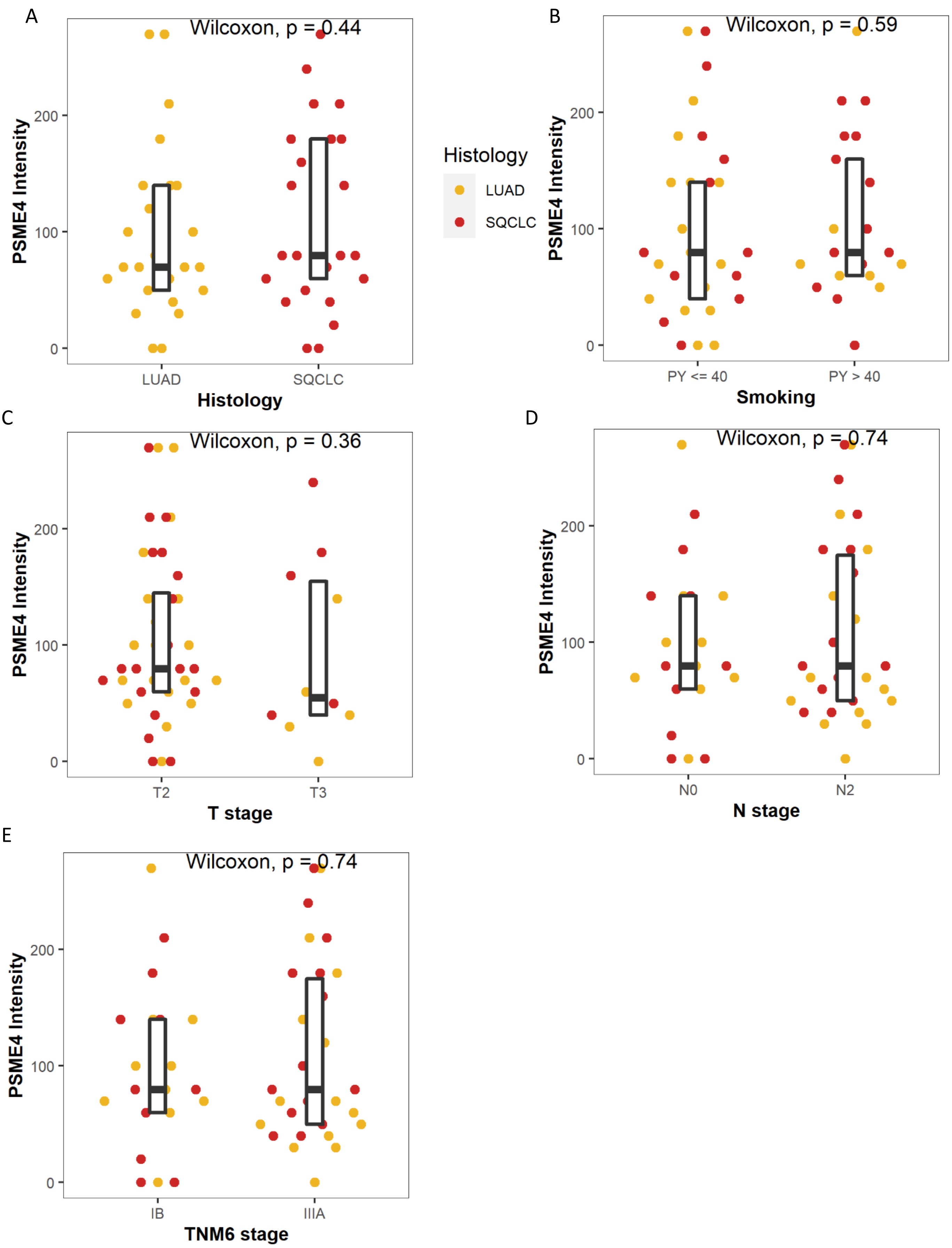

A
Wilcoxon, p = 0.44
PSME4 Intensity
0
100
200
LUAD
SQCLC
Histology
Histology
LUAD
SQCLC
B
Wilcoxon, p = 0.59
PSME4 Intensity
PY <= 40
PY > 40
Smoking
C
Wilcoxon, p = 0.36
PSME4 Intensity
T2
T3
T stage
D
Wilcoxon, p = 0.74
PSME4 Intensity
N0
N2
N stage
E
Wilcoxon, p = 0.74
PSME4 Intensity
IB
IIIA
TNM6 stage

### Figure S6

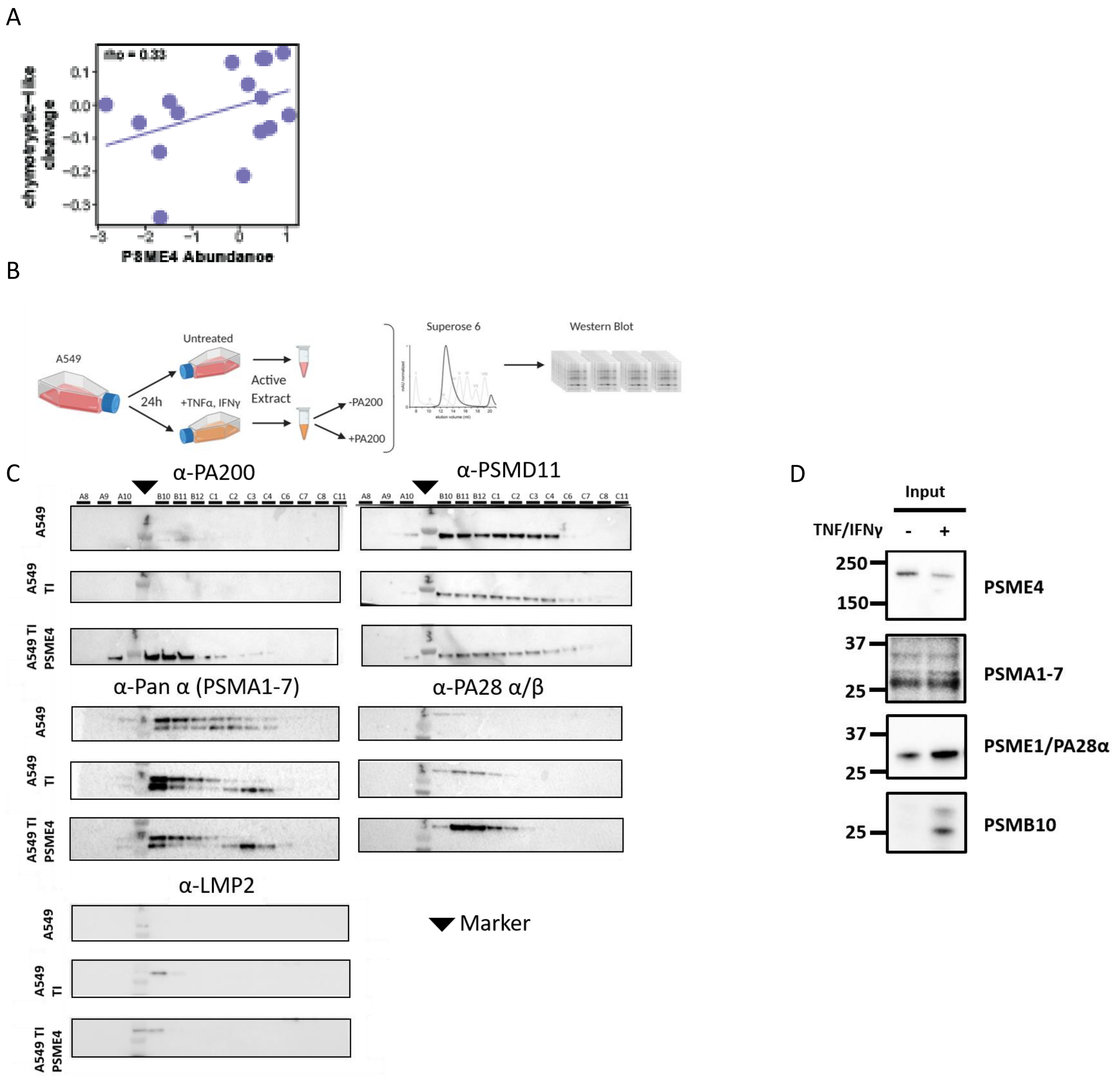
A
rho = 0.33
chymotrypsic-like cleavage
PSME4 Abundance
B
A549
Untreated
24h
+TNFα, IFNγ
Active Extract
-PA200
+PA200
Superose 6
Western Blot
C
α-PA200
α-PSMD11
A8 A9 A10 B10 B11 B12 C1 C2 C3 C4 C6 C7 C8 C11
A549
A549 TI
A549 TI PSME4
α-Pan α (PSMA1-7)
α-PA28 α/β
α-LMP2
Marker
D
Input
TNF/IFNγ - +
250
150
PSME4
37
25
PSMA1-7
PSME1/PA28α
PSMB10

### Figure S7

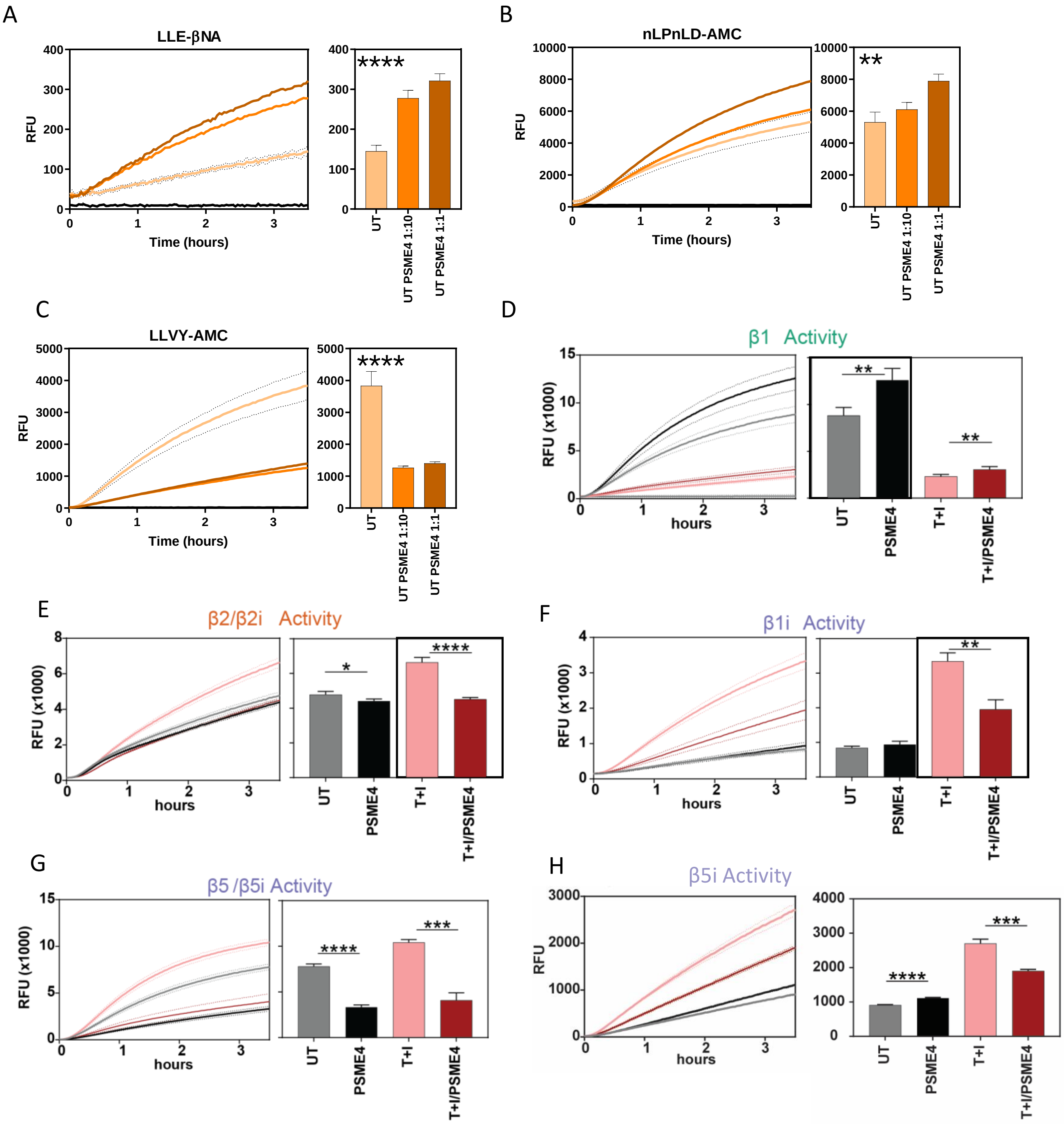
A
LLE-βNA
RFU
Time (hours)
UT
UT PSME4 1:10
UT PSME4 1:1
****
B
nLPnLD-AMC
RFU
Time (hours)
UT
UT PSME4 1:10
UT PSME4 1:1
**
C
LLVY-AMC
RFU
Time (hours)
UT
UT PSME4 1:10
UT PSME4 1:1
****
D
β1 Activity
RFU (x1000)
hours
UT
PSME4
T+I
T+I/PSME4
E
β2/β2i Activity
RFU (x1000)
hours
UT
PSME4
T+I
T+I/PSME4
F
β1i Activity
RFU (x1000)
hours
UT
PSME4
T+I
T+I/PSME4
G
β5 /β5i Activity
RFU (x1000)
hours
UT
PSME4
T+I
T+I/PSME4
H
β5i Activity
RFU
hours
UT
PSME4
T+I
T+I/PSME4

### Figure S8

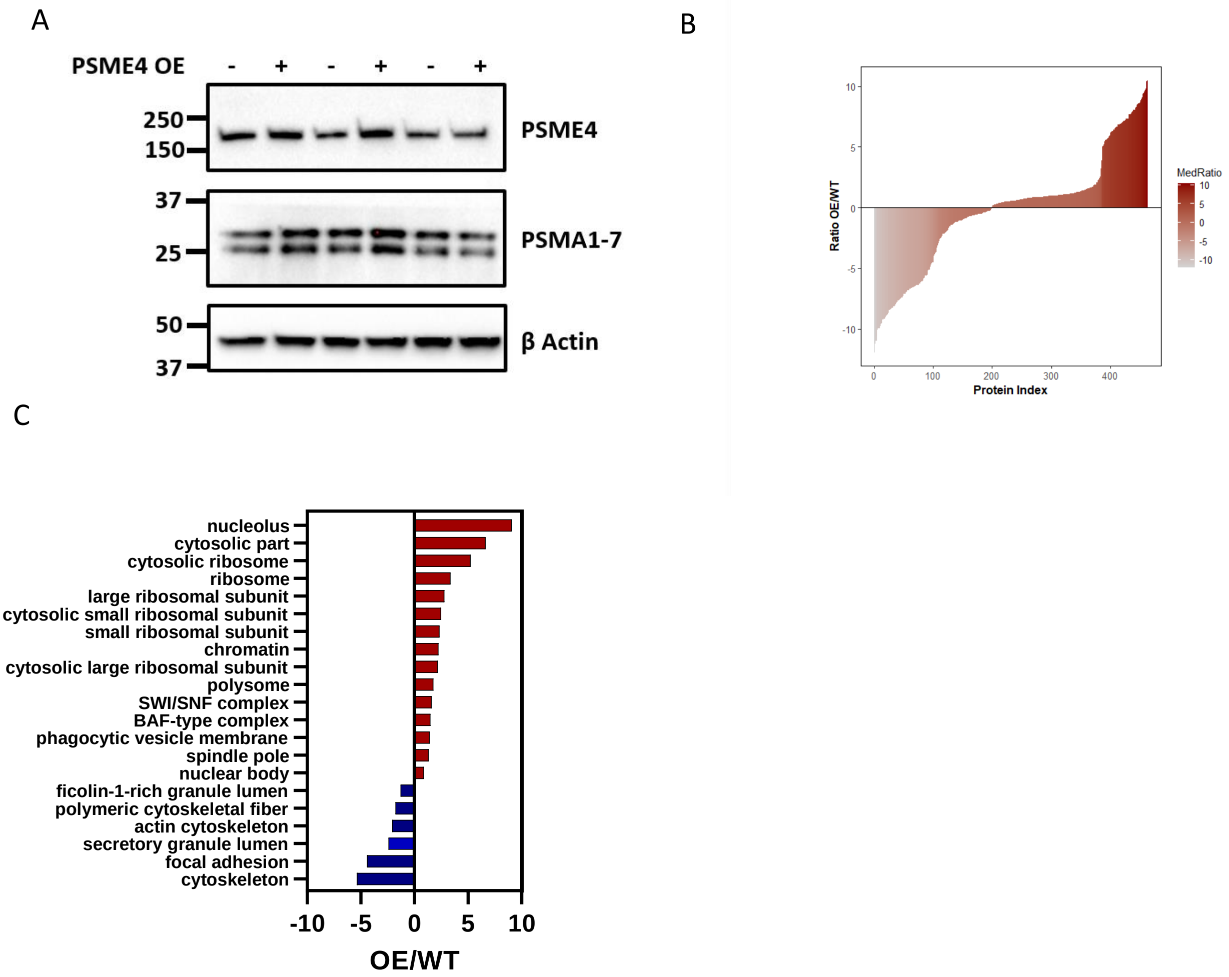
A
PSME4 OE
- + - + - +
250
150
PSME4
37
25
PSMA1-7
50
37
β Actin
B
Ratio OE/WT
10
5
0
-5
-10
0
100
200
300
400
Protein Index
MedRatio
10
5
0
-5
-10
C
nucleolus
cytosolic part
cytosolic ribosome
ribosome
large ribosomal subunit
cytosolic small ribosomal subunit
small ribosomal subunit
chromatin
cytosolic large ribosomal subunit
polysome
SWI/SNF complex
BAF-type complex
phagocytic vesicle membrane
spindle pole
nuclear body
ficolin-1-rich granule lumen
polymeric cytoskeletal fiber
actin cytoskeleton
secretory granule lumen
focal adhesion
cytoskeleton
-10
-5
0
5
10
OE/WT

Figure S9

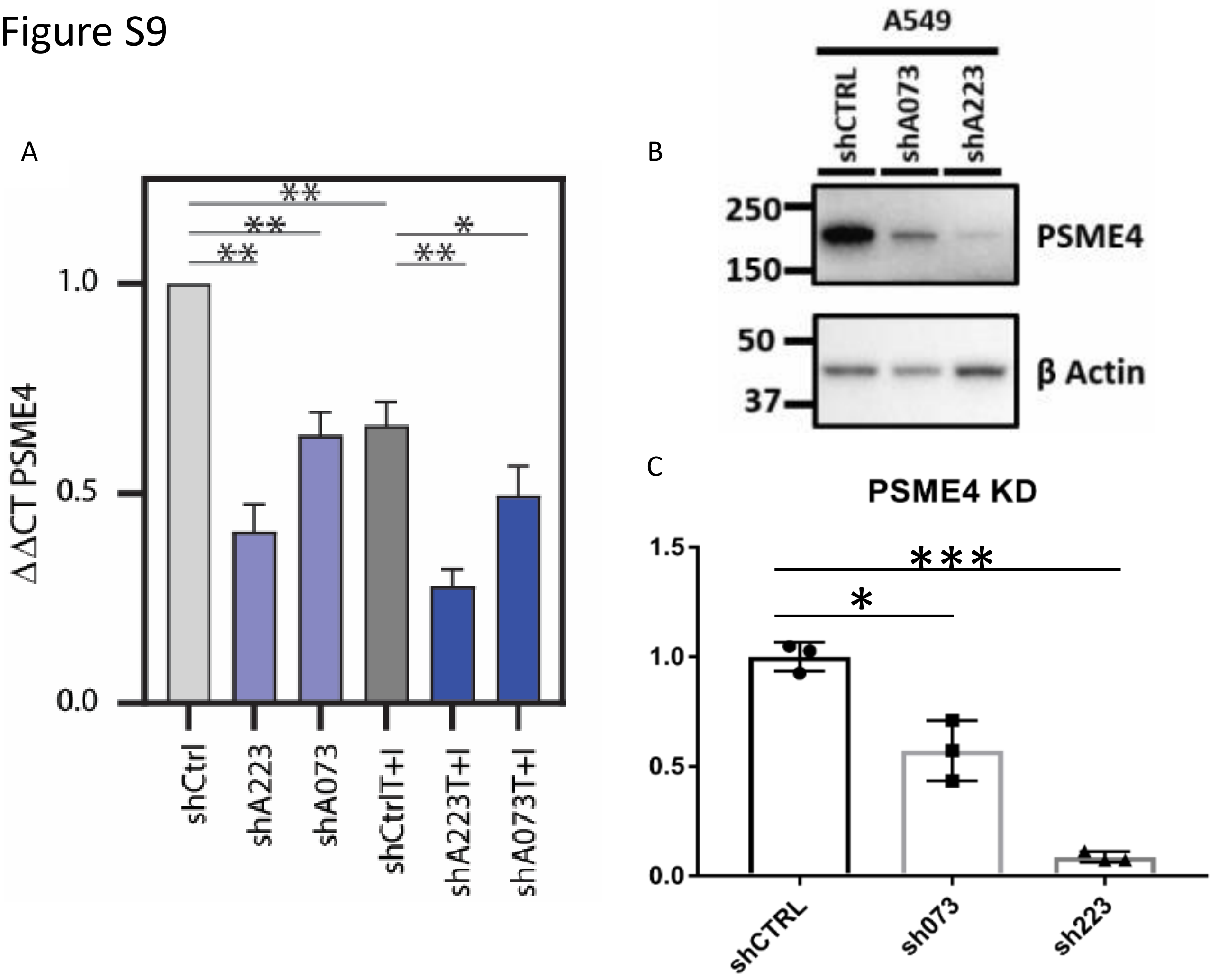
A
B
C
**
**
**
*
**
1.0
0.5
0.0
ΔΔCT PSME4
shCtrl
shA223
shA073
shCtrlT+I
shA223T+I
shA073T+I
A549
shCTRL
shA073
shA223
250
150
PSME4
50
37
β Actin
PSME4 KD
***
*
1.5
1.0
0.5
0.0
shCTRL
sh073
sh223

### Figure S10

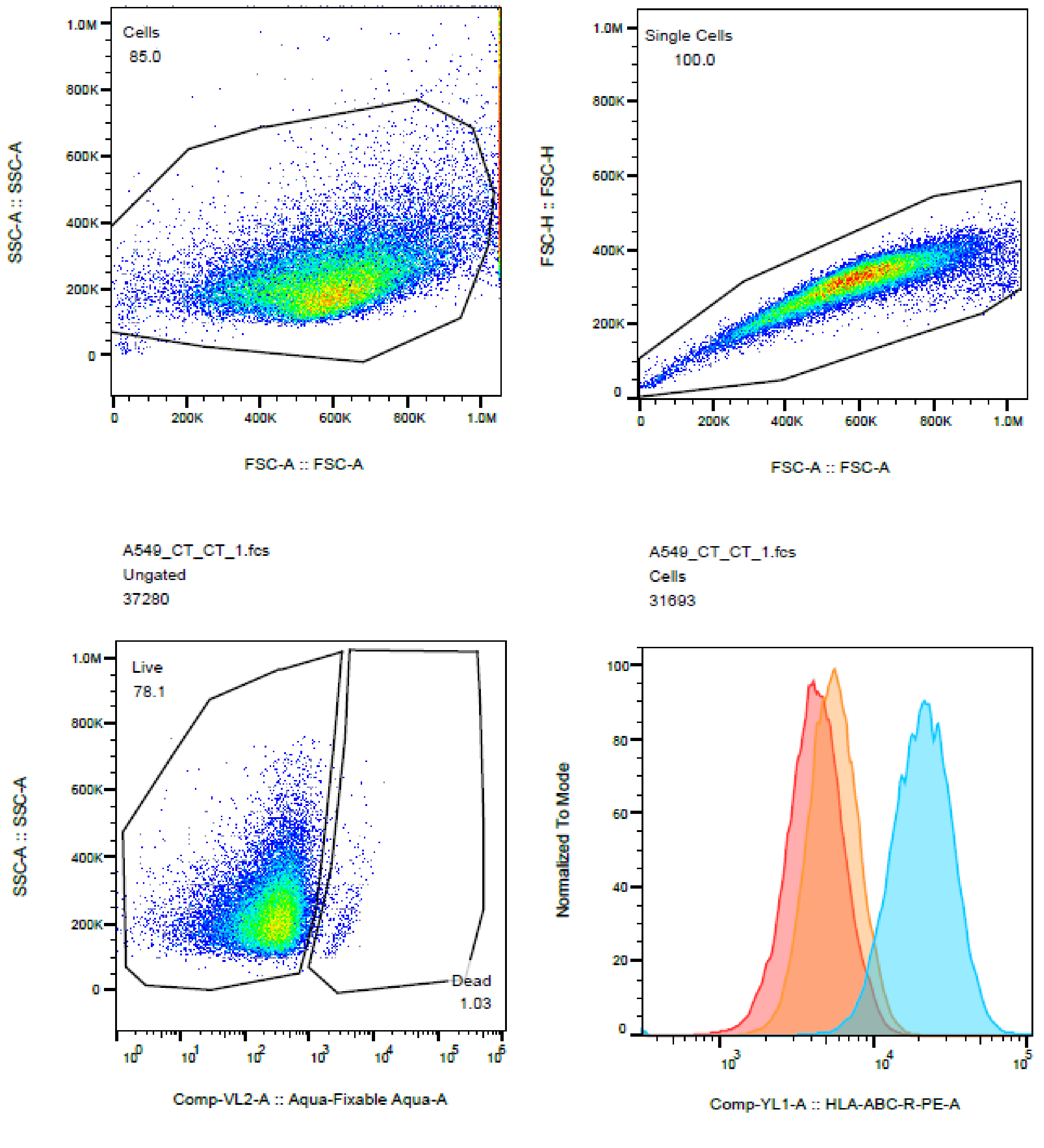


A549_CT_CT_1.fcs
Single Cells
31684

| | Sample Name | Subset Name | Count | Median : Comp-YL1-A |
|---|---|---|---|---|
| | A549_CT_TI_1.fcs | Live | 8698 | 20819 |
| | A549_KD223_CT_1.fc | Live | 21452 | 5480 |
| | A549_CT_CT_1.fcs | Live | 24758 | 4222 |

### Figure S11

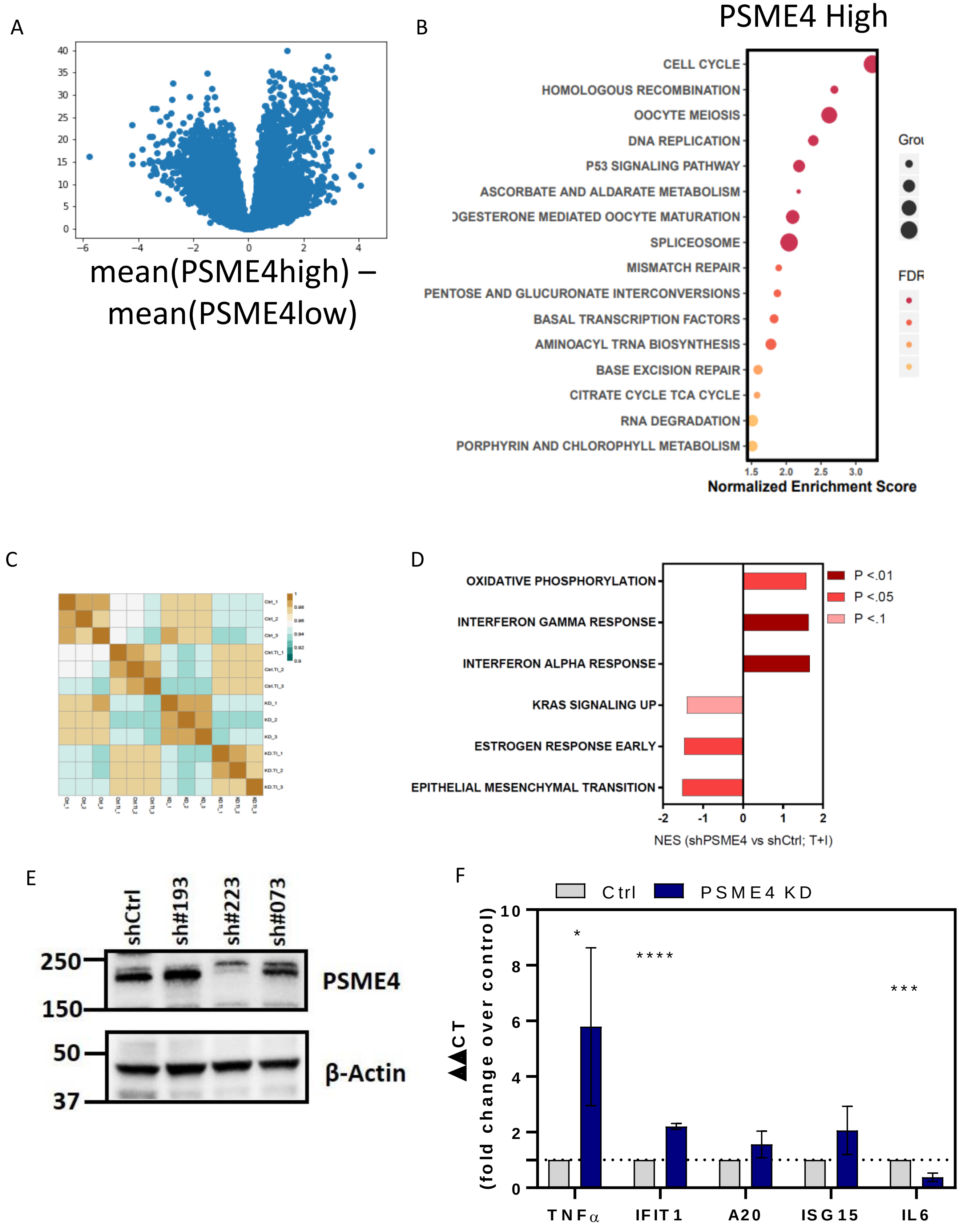
A
40
35
30
25
20
15
10
5
0
−6
−4
−2
0
2
4
mean(PSME4high) – mean(PSME4low)
B
PSME4 High
CELL CYCLE
HOMOLOGOUS RECOMBINATION
OOCYTE MEIOSIS
DNA REPLICATION
P53 SIGNALING PATHWAY
ASCORBATE AND ALDARATE METABOLISM
OGESTERONE MEDIATED OOCYTE MATURATION
SPLICEOSOME
MISMATCH REPAIR
PENTOSE AND GLUCURONATE INTERCONVERSIONS
BASAL TRANSCRIPTION FACTORS
AMINOACYL TRNA BIOSYNTHESIS
BASE EXCISION REPAIR
CITRATE CYCLE TCA CYCLE
RNA DEGRADATION
PORPHYRIN AND CHLOROPHYLL METABOLISM
1.5
2.0
2.5
3.0
Normalized Enrichment Score
Grou
FDR
C
D
OXIDATIVE PHOSPHORYLATION
INTERFERON GAMMA RESPONSE
INTERFERON ALPHA RESPONSE
KRAS SIGNALING UP
ESTROGEN RESPONSE EARLY
EPITHELIAL MESENCHYMAL TRANSITION
-2
-1
0
1
2
NES (shPSME4 vs shCtrl; T+I)
P <.01
P <.05
P <.1
E
shCtrl
sh#193
sh#223
sh#073
250
150
PSME4
50
37
β-Actin
F
Ctrl
PSME4 KD
ΔΔCT (fold change over control)
10
8
6
4
2
0
*
****
***
TNFα
IFIT1
A20
ISG15
IL6

### Figure S12

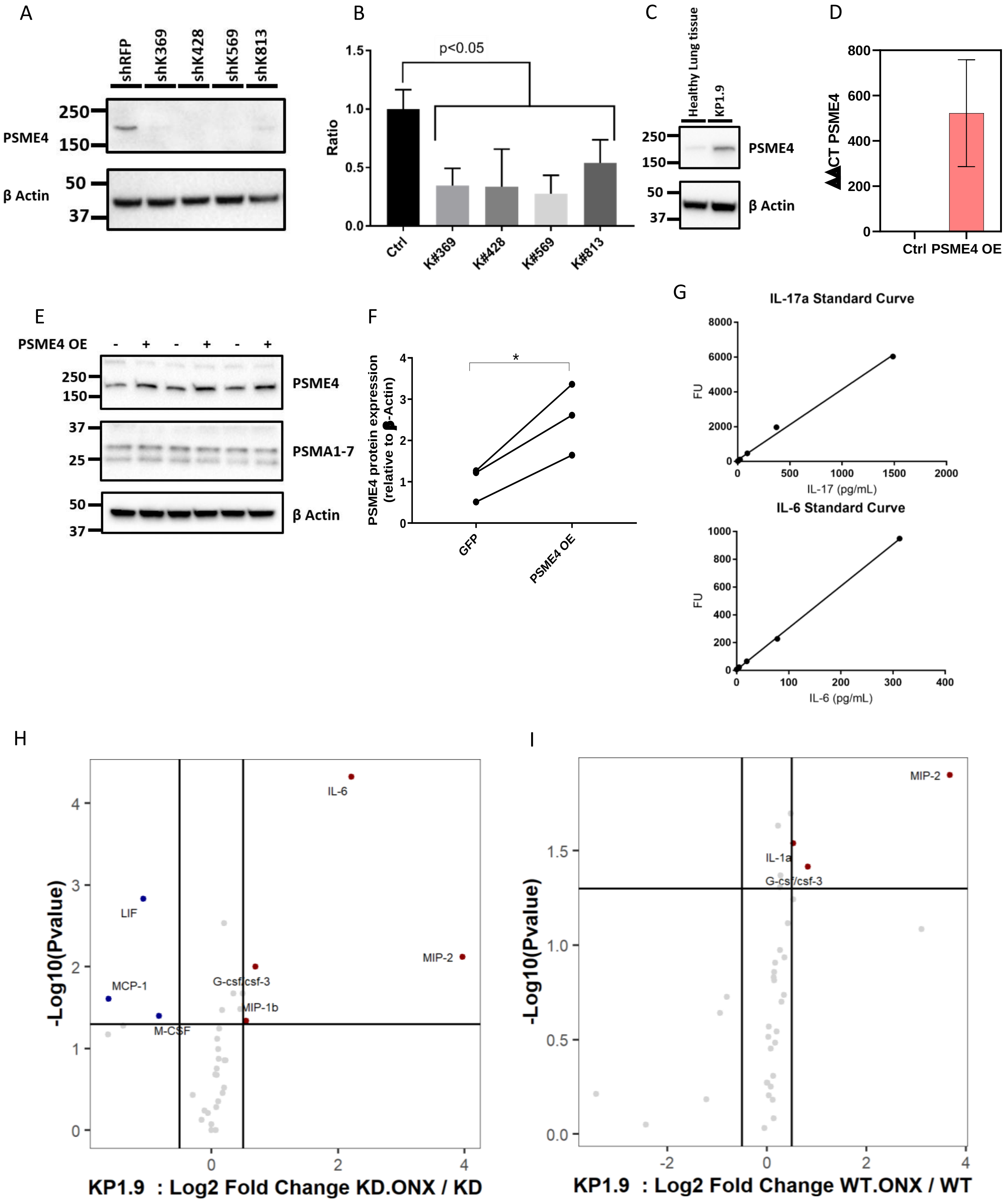
A
shRFP
shK369
shK428
shK569
shK813
250
150
50
37
PSME4
β Actin
B
p<0.05
Ratio
1.5
1.0
0.5
0.0
Ctrl
K#369
K#428
K#569
K#813
C
Healthy Lung tissue
KP1.9
PSME4
β Actin
D
ΔΔCT PSME4
800
600
400
200
0
Ctrl
PSME4 OE
E
PSME4 OE
PSME4
PSMA1-7
β Actin
F
PSME4 protein expression (relative to β-Actin)
GFP
PSME4 OE
*
G
IL-17a Standard Curve
FU
IL-17 (pg/mL)
IL-6 Standard Curve
IL-6 (pg/mL)
H
-Log10(Pvalue)
IL-6
LIF
MIP-2
G-csf/csf-3
MCP-1
MIP-1b
M-CSF
KP1.9 : Log2 Fold Change KD.ONX / KD
I
MIP-2
IL-1a
G-csf/csf-3
KP1.9 : Log2 Fold Change WT.ONX / WT

### Figure S13

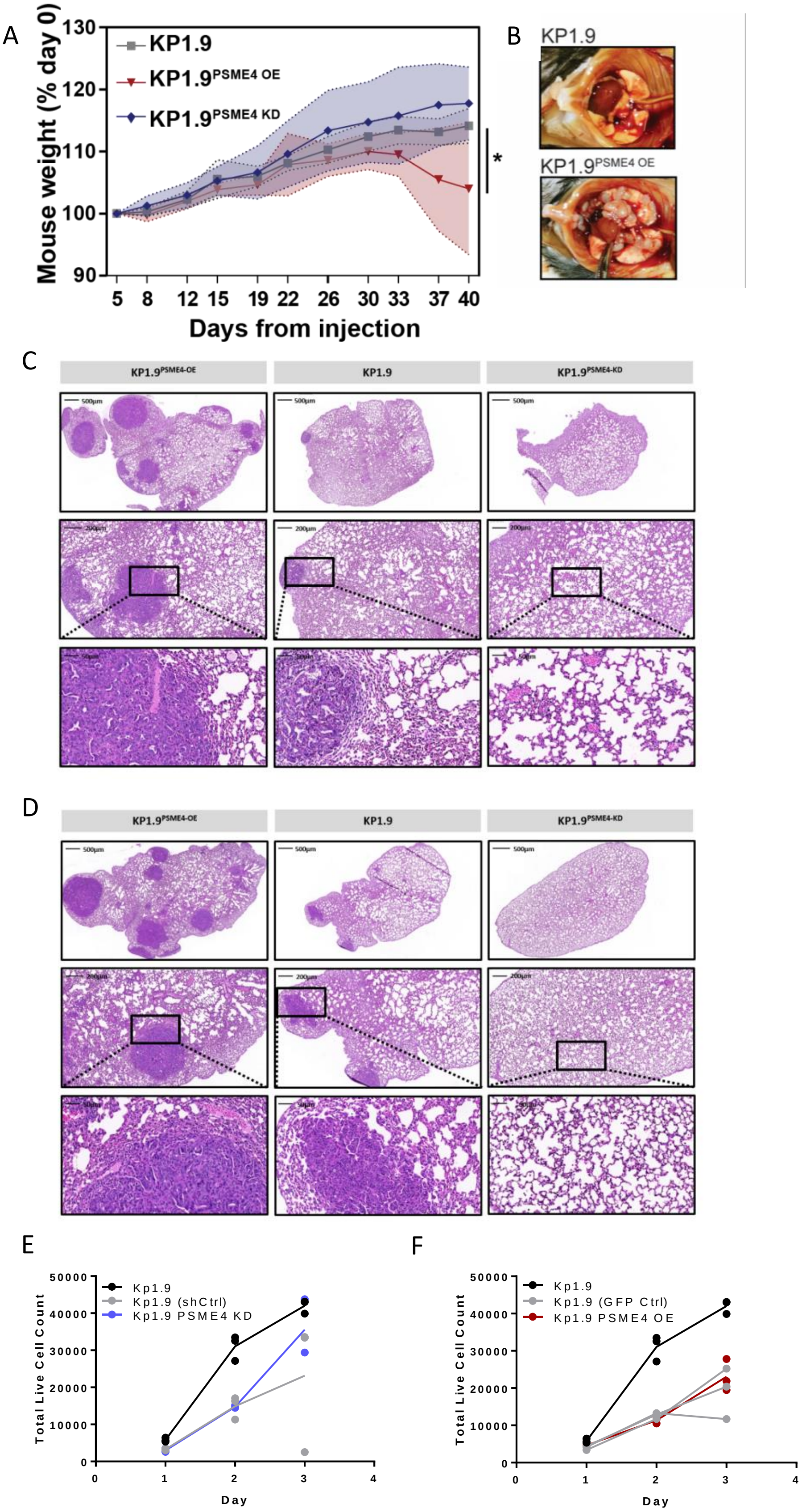
A
B
C
D
E
F
Mouse weight (% day 0)
130
120
110
100
90
KP1.9
KP1.9PSME4 OE
KP1.9PSME4 KD
*
5 8 12 15 19 22 26 30 33 37 40
Days from injection
KP1.9
KP1.9PSME4 OE
KP1.9PSME4-OE
KP1.9
KP1.9PSME4-KD
Total Live Cell Count
50000
40000
30000
20000
10000
0
Kp1.9
Kp1.9 (shCtrl)
Kp1.9 PSME4 KD
Kp1.9 (GFP Ctrl)
Kp1.9 PSME4 OE
0 1 2 3 4
Day

### Figure S14

A

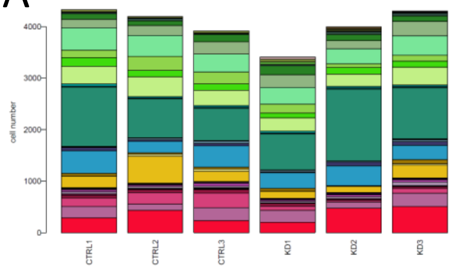


B

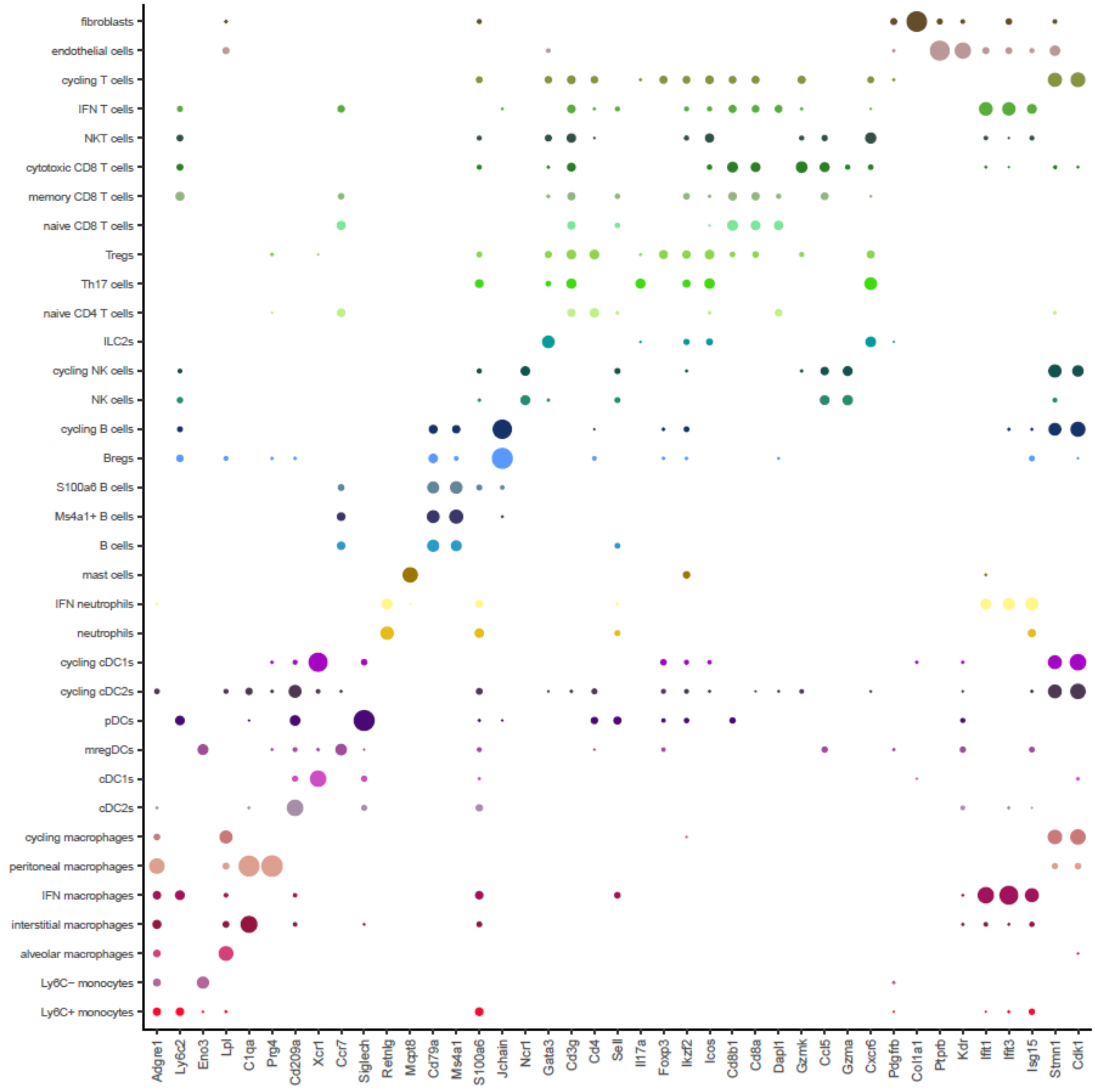

Figure S15

A

pDCs

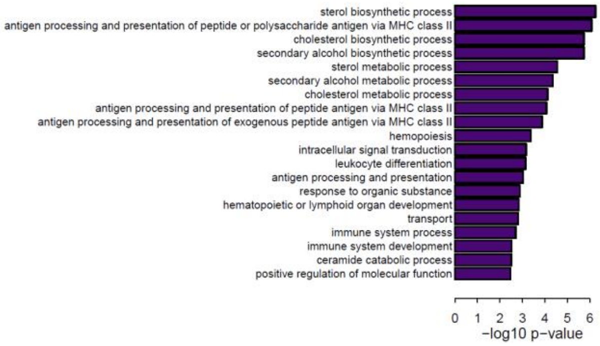


B

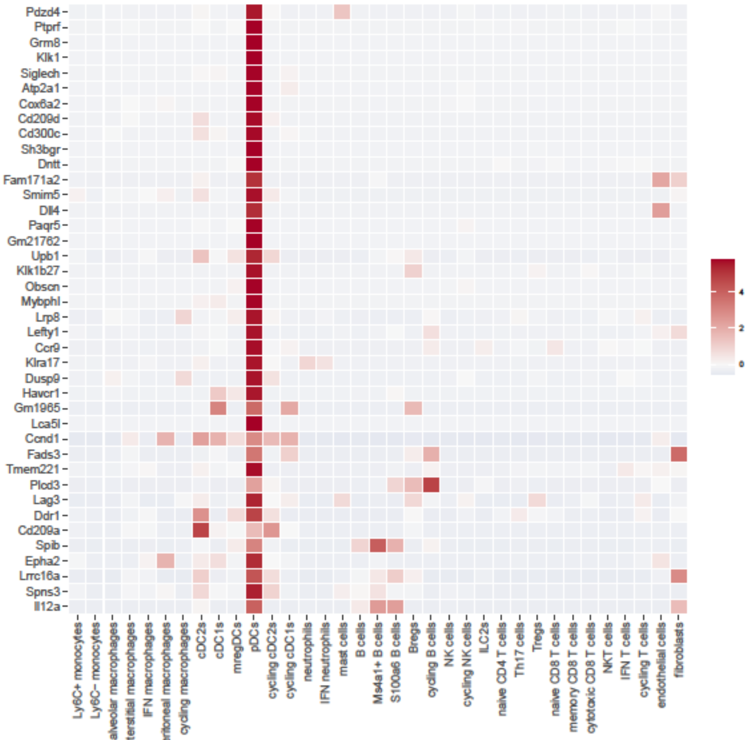


C

mregDCs

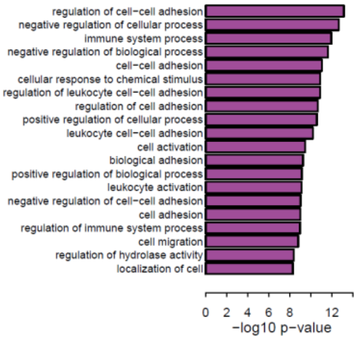


D

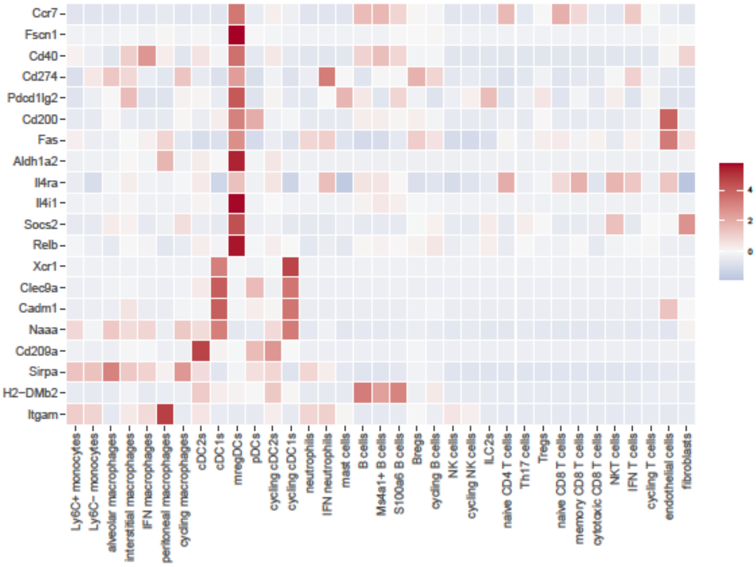

Figure S16

A

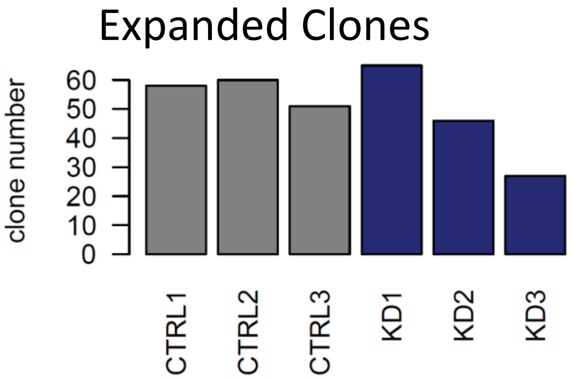


B

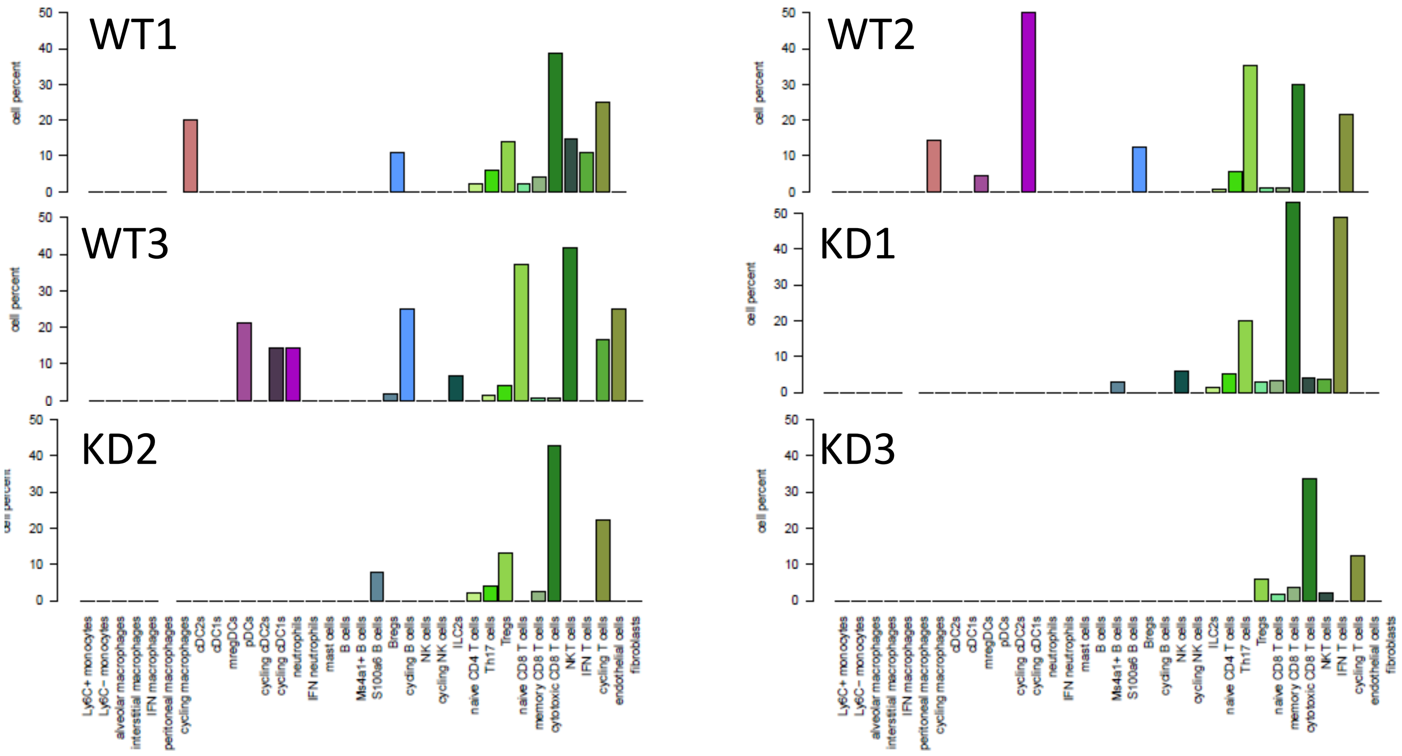

### Figure S17

A

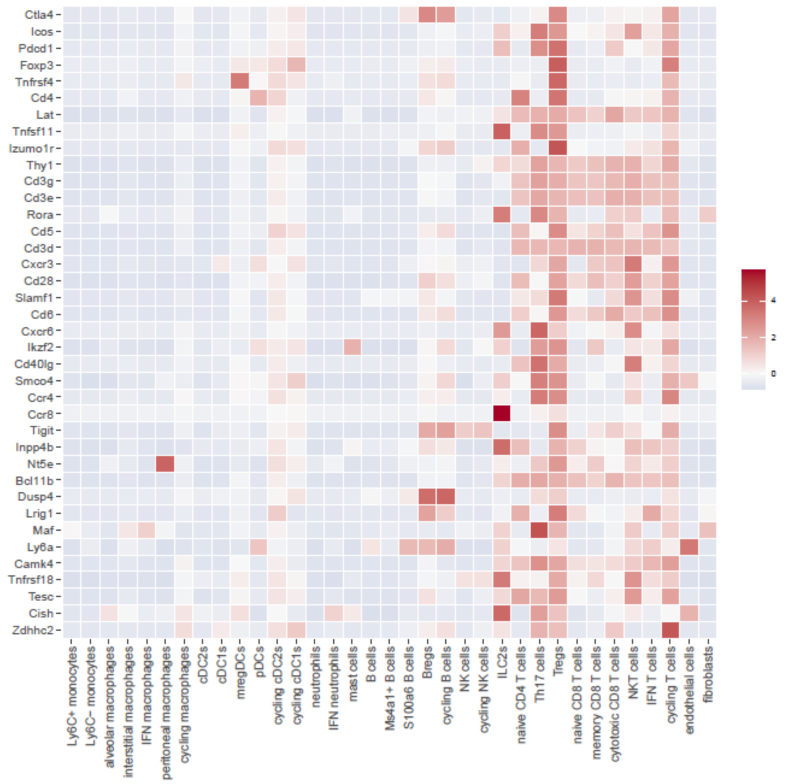


B

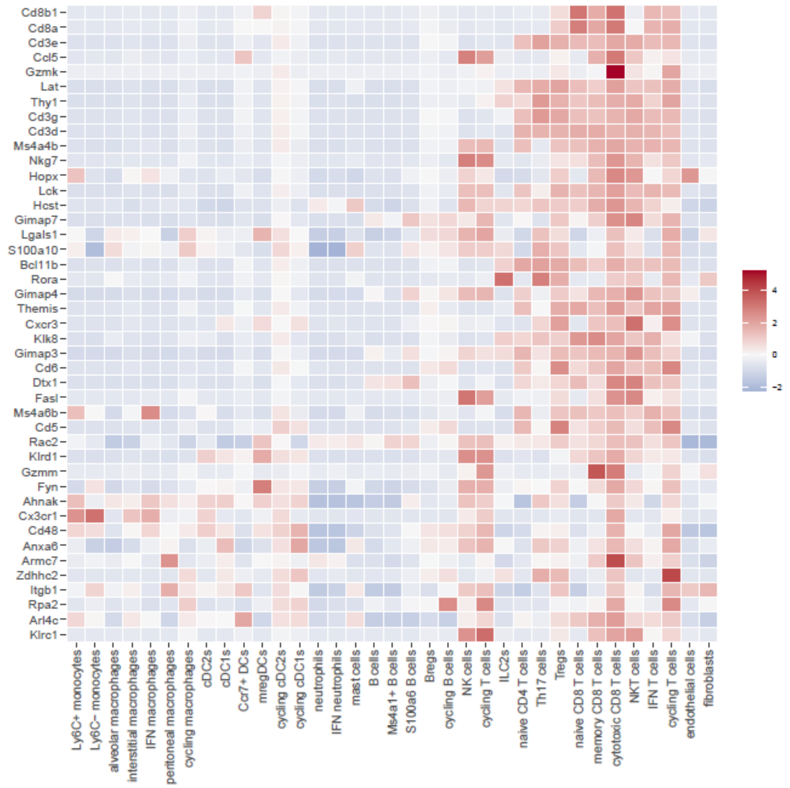

Figure S18

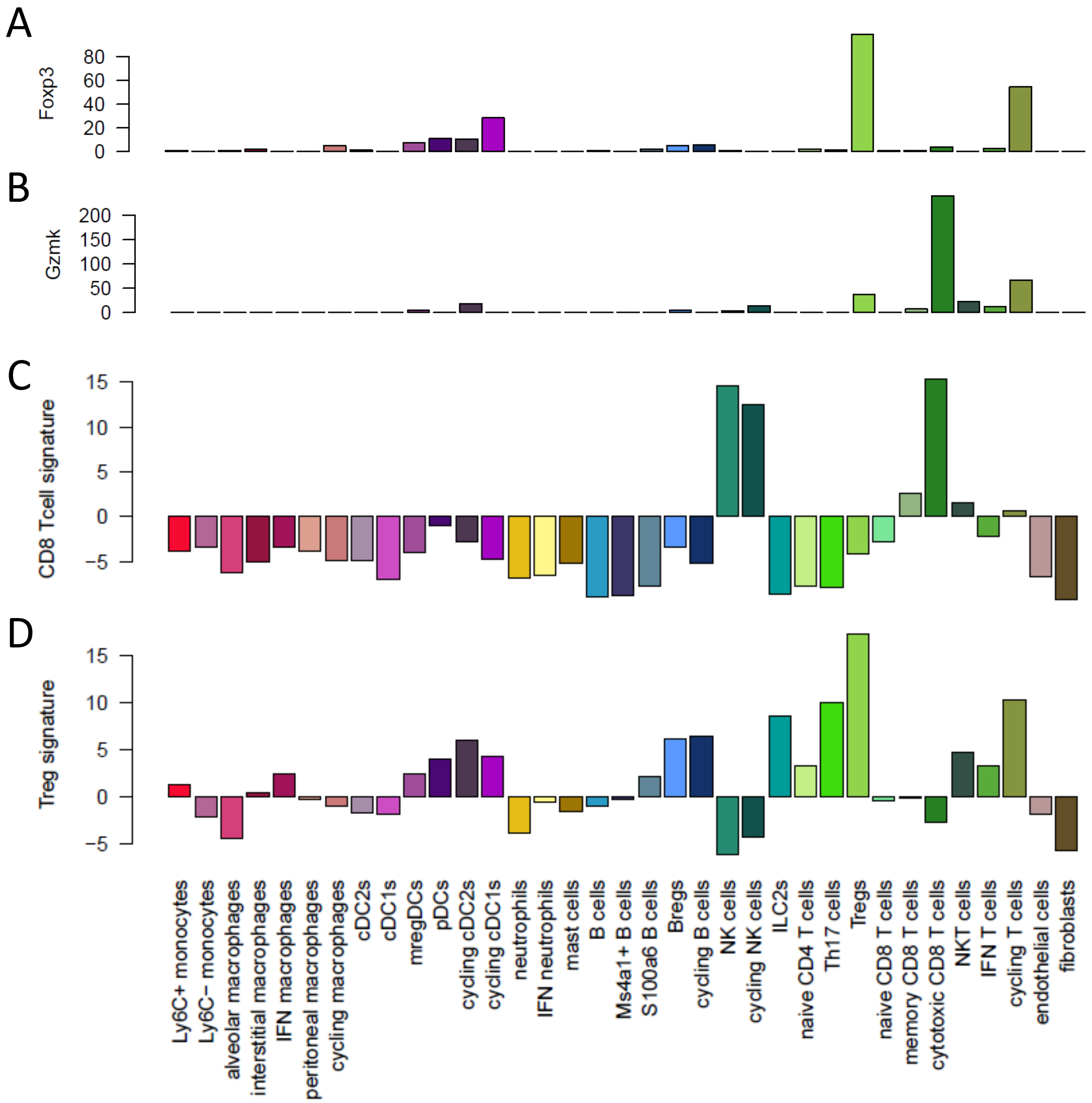
A
Foxp3
0
20
40
60
80
B
Gzmk
0
50
100
150
200
C
CD8 Tcell signature
−5
0
5
10
15
D
Treg signature
−5
0
5
10
15
Ly6C+ monocytes
Ly6C− monocytes
alveolar macrophages
interstitial macrophages
IFN macrophages
peritoneal macrophages
cycling macrophages
cDC2s
cDC1s
mregDCs
pDCs
cycling cDC2s
cycling cDC1s
neutrophils
IFN neutrophils
mast cells
B cells
Ms4a1+ B cells
S100a6 B cells
Bregs
cycling B cells
NK cells
cycling NK cells
ILC2s
naive CD4 T cells
Th17 cells
Tregs
naive CD8 T cells
memory CD8 T cells
cytotoxic CD8 T cells
NKT cells
IFN T cells
cycling T cells
endothelial cells
fibroblasts

### Figure S19

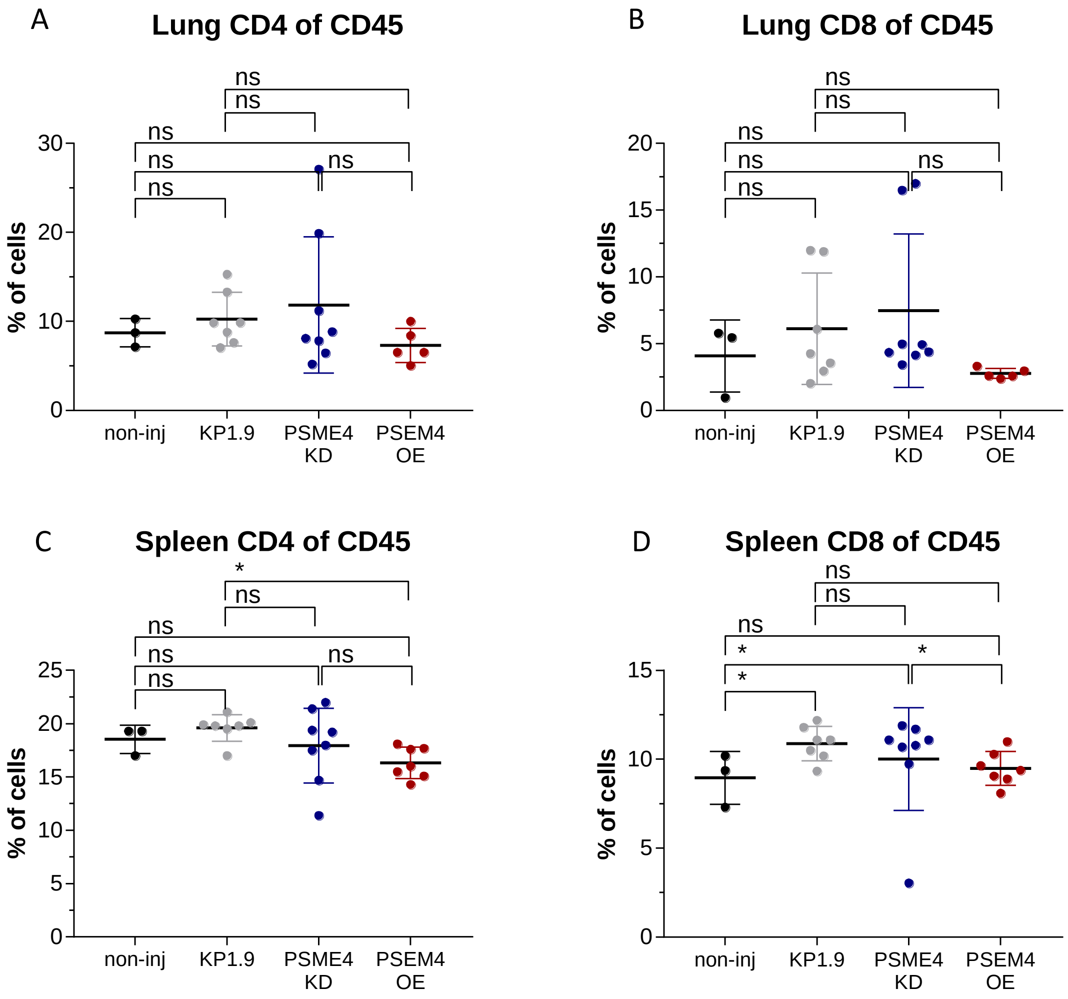

A
Lung CD4 of CD45
% of cells
non-inj
KP1.9
PSME4 KD
PSEM4 OE
ns
B
Lung CD8 of CD45
% of cells
non-inj
KP1.9
PSME4 KD
PSEM4 OE
ns
C
Spleen CD4 of CD45
% of cells
non-inj
KP1.9
PSME4 KD
PSEM4 OE
ns
*
D
Spleen CD8 of CD45
% of cells
non-inj
KP1.9
PSME4 KD
PSEM4 OE
ns
*

### Figure S20

A

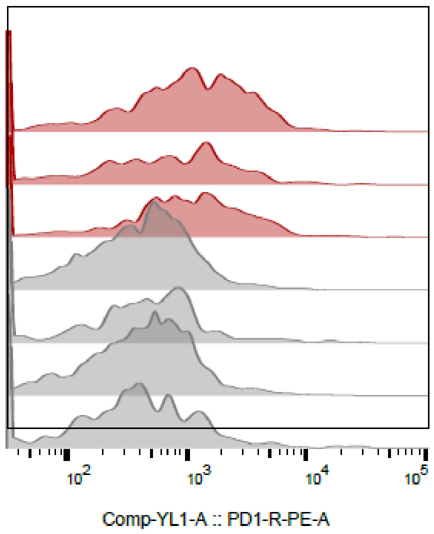


B

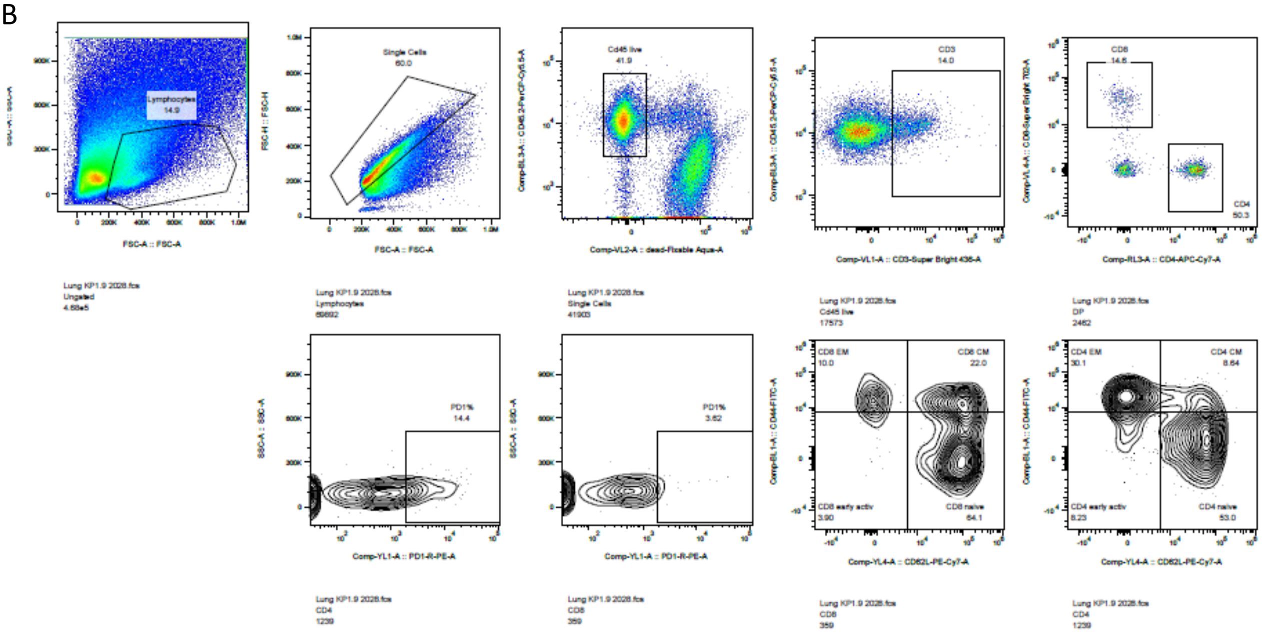


C

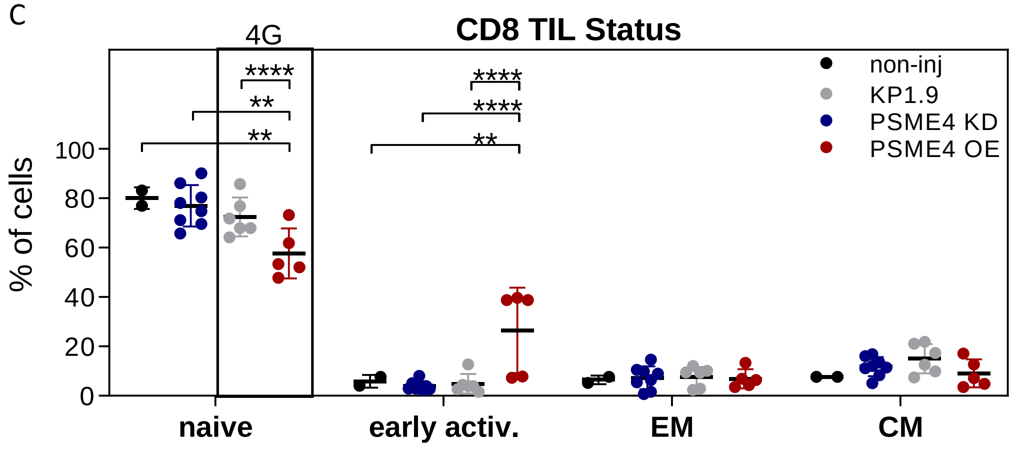

### Figure S21

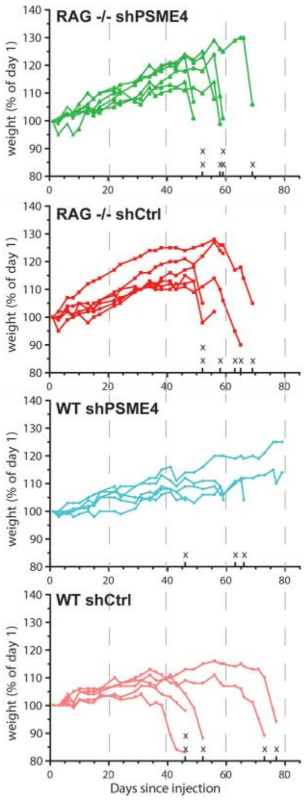

RAG -/- shPSME4
RAG -/- shCtrl
WT shPSME4
WT shCtrl
weight (% of day 1)
Days since injection

### Figure S22

A

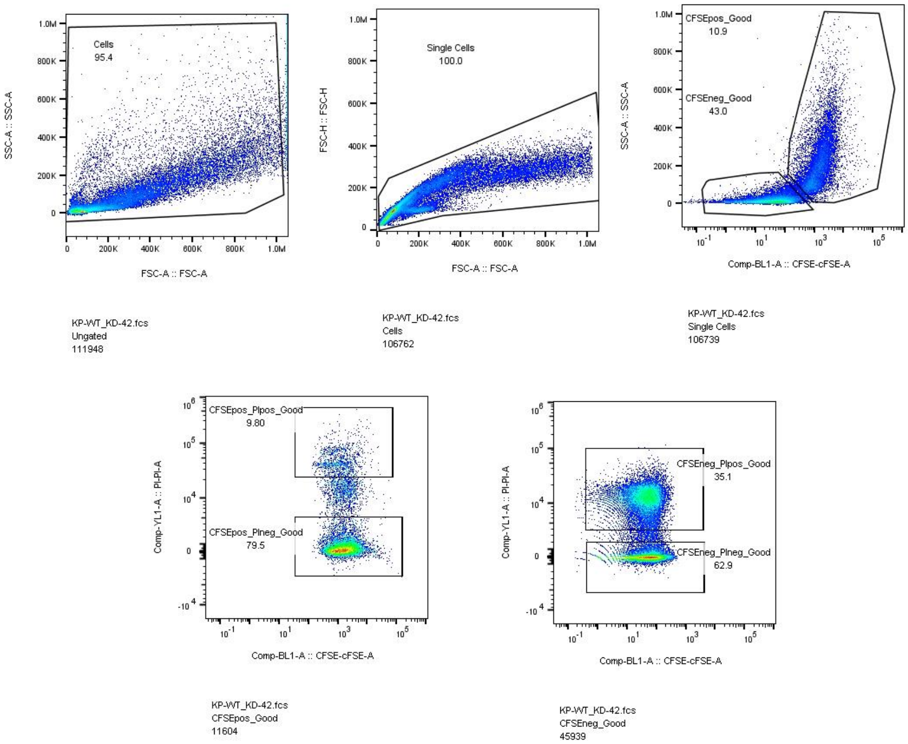


B

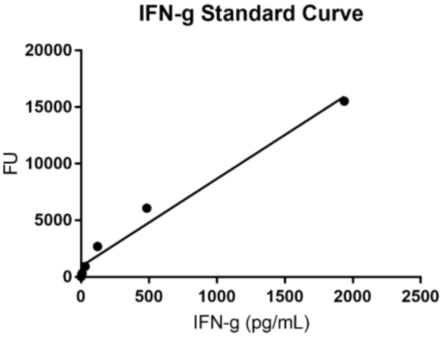


C

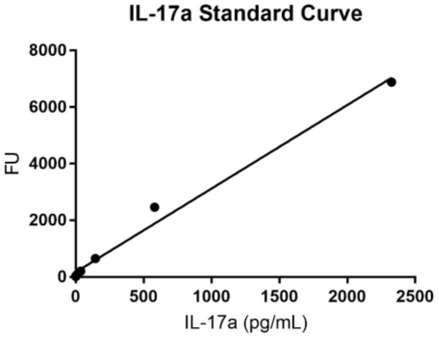


D

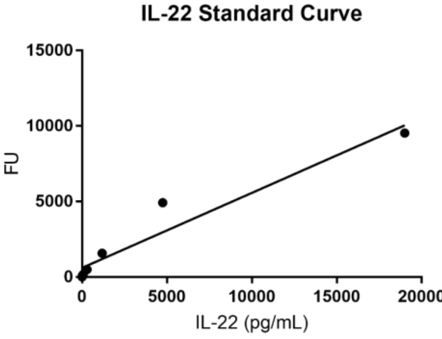

### Figure S23

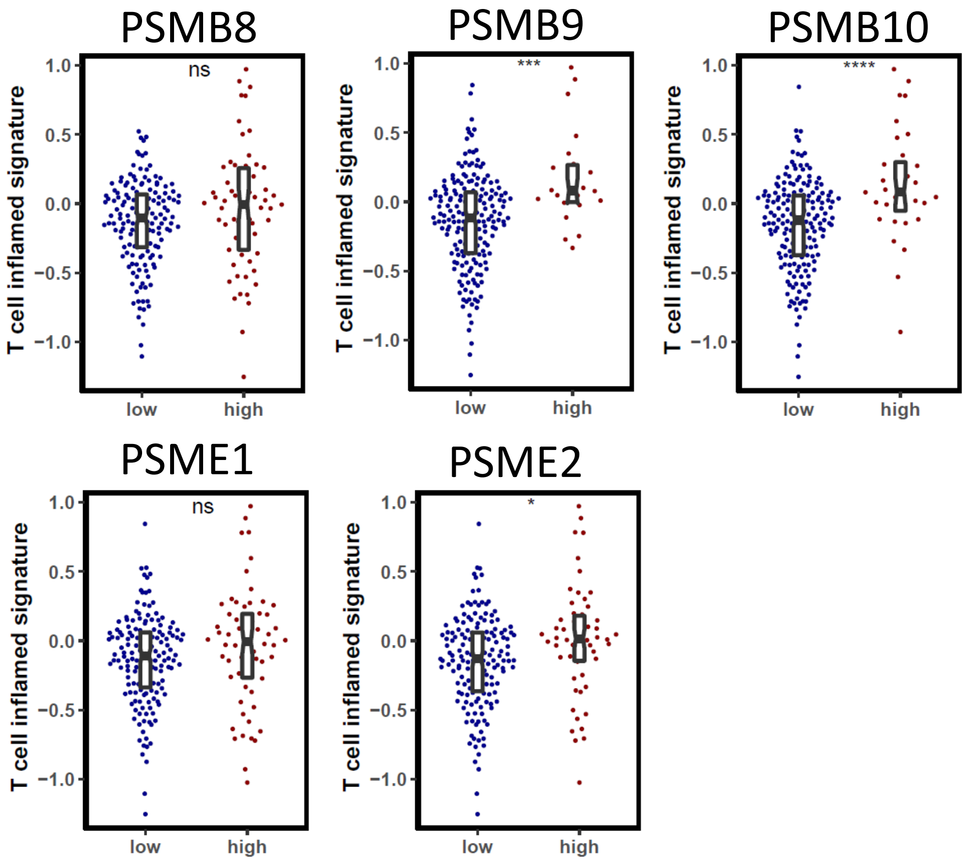

PSMB8
ns
PSMB9
***
PSMB10
****
PSME1
ns
PSME2
*
T cell inflamed signature
1.0
0.5
0.0
−0.5
−1.0
low
high

### Figure S24

A

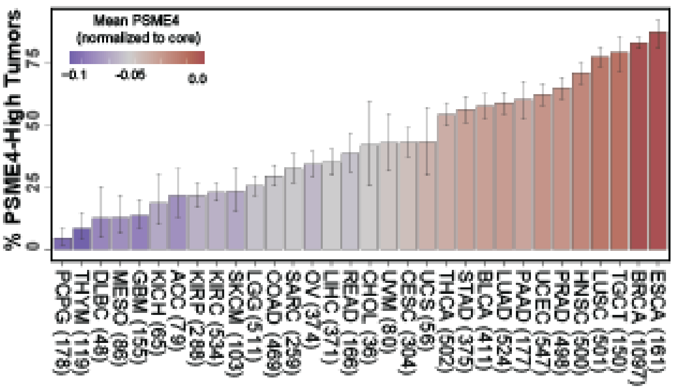


B

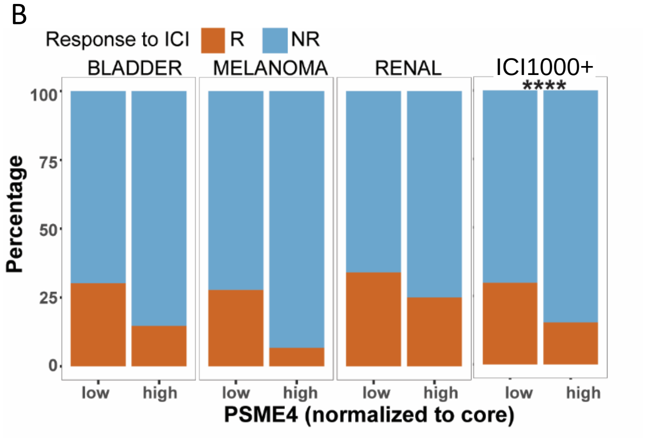


C

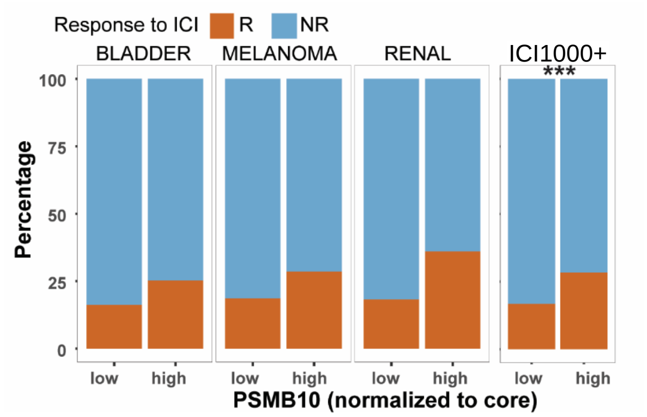

### Figure S25

A

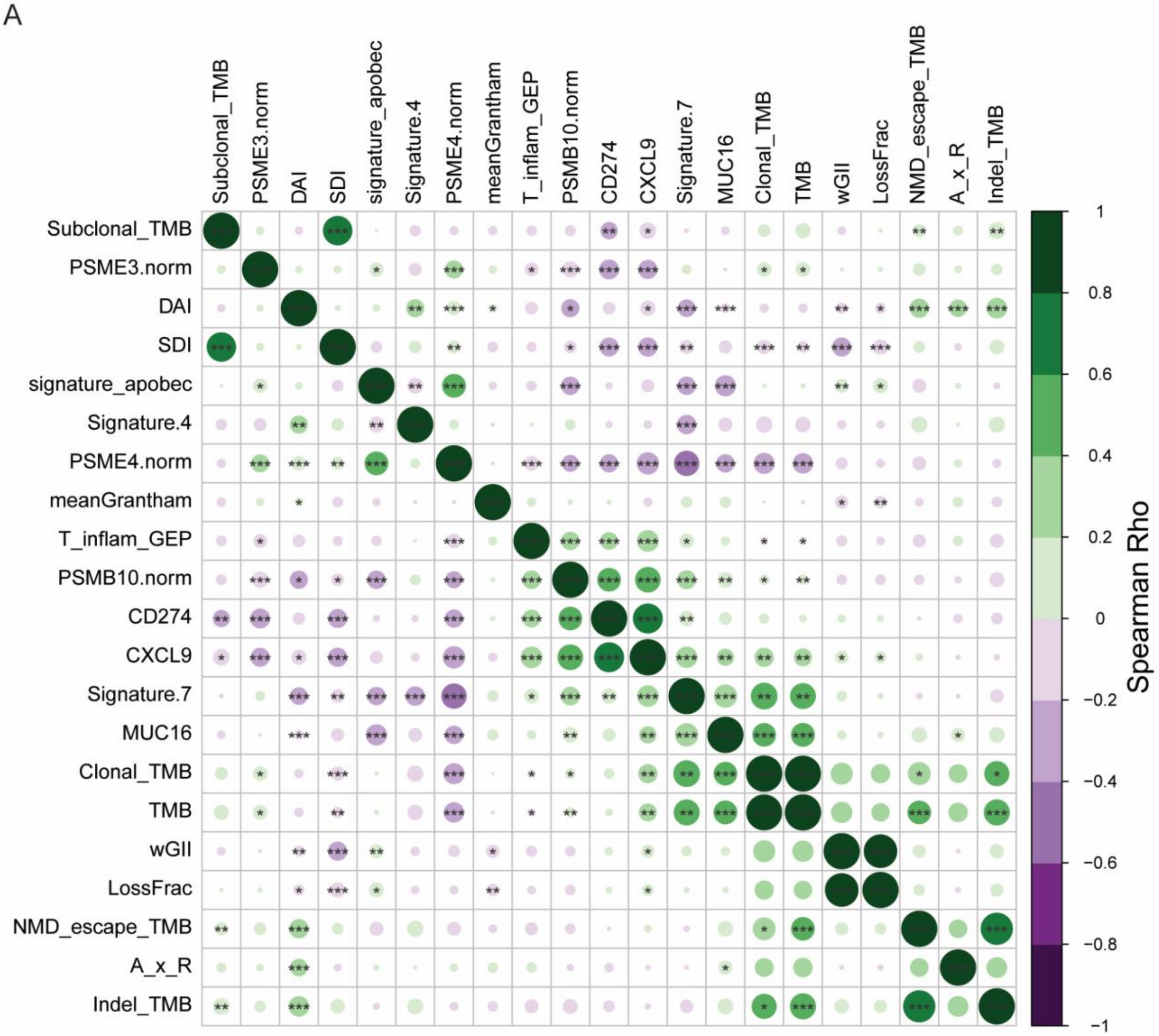


B

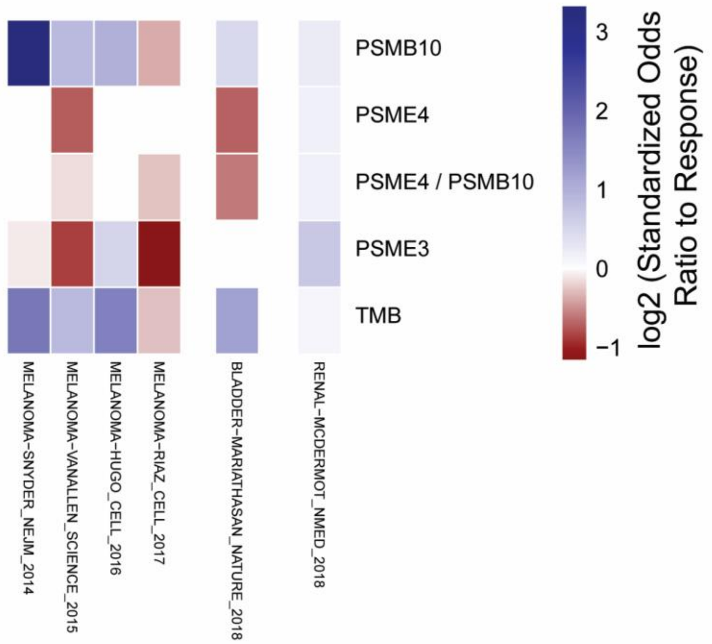

### Figure S26

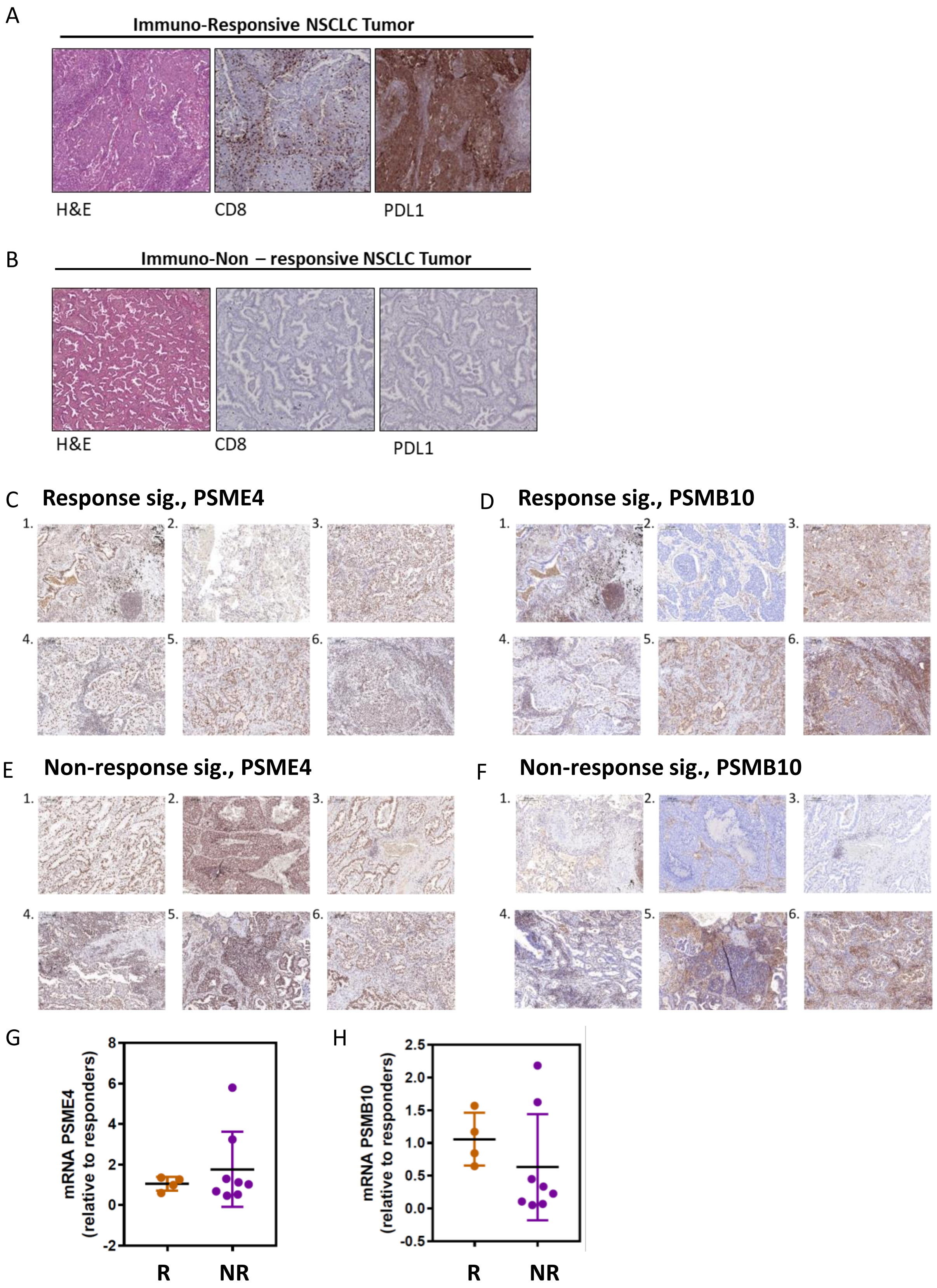

A
Immuno-Responsive NSCLC Tumor
H&E
CD8
PDL1
B
Immuno-Non – responsive NSCLC Tumor
H&E
CD8
PDL1
C
Response sig., PSME4
1.
2.
3.
4.
5.
6.
D
Response sig., PSMB10
1.
2.
3.
4.
5.
6.
E
Non-response sig., PSME4
1.
2.
3.
4.
5.
6.
F
Non-response sig., PSMB10
1.
2.
3.
4.
5.
6.
G
mRNA PSME4 (relative to responders)
8
6
4
2
0
-2
R
NR
H
mRNA PSMB10 (relative to responders)
2.5
2.0
1.5
1.0
0.5
0.0
-0.5
R
NR

Figure S1.

**Proteome and Degradome landscapes classify tumor and adjacent tissues across patients. (A-B)** Pairwise Pearson correlation between the proteins identified in each sample by MAPP (A) or the whole cell proteome (B **(C-D)** Principal component analysis based on the identities and abundances of the proteins identified by MAPP (C) or whole cell proteome (D). Tumor (T) and adjacent (A) samples are annotated and plotted with principal component PC1 against PC2.

Figure S2.

**Proteins selectively degraded in tumor or adjacent tissue across patients. (A)** The abundance of proteins that were different between tumor and adjacent lung tissue as determined by MAPP are shown in four datasets: MAPP, WC Proteomics, CPTAC proteomics and TCGA LUAD. Proteins not identified in a dataset are shown as grey, otherwise the normalized abundance / expression is shown as a red to blue scale. **(B)** The fold change in abundance for each protein between the tumor and adjacent tissue in the WC proteome is plotted against the significance of their difference (negative log10 transformed P value) in grey. The proteins identified as differential in MAPP (listed in table S2) are annotated in red.

Figure S3.

**Nuclear degradation is enriched in lung adenocarcinoma. (A)** Proteins identified in MAPP organized by their annotated cellular localization (Human Protein Atlas). Proteins are colored based on the mean of degradation ratios between tumor and adjacent tissues across patients (orange: above median degradation ratio, blue: degradation ratio below 0, black: ratio between median and 0). Proteins that are annotated to localize at more than one organelle appear more than once. **(B-C)** The mean ratio of degradation (MAPP) between the tumor and adjacent tissues for proteins annotated to the nucleus (B) or cytosol (C) versus other proteins. The nuclear proteins on average had significantly higher ratio compared to the background proteins (Wilcoxon p = 0.037).

Figure S4

**PSME4 is increased in expression in NSCLC (A)** The tumor to adjacent tissue ratio of the abundance of all proteasome subunits based on the proteasome immunoprecipitation (top) or across the samples of the CPTAC lung adenocarcinoma cohort (bottom). Colors and diagrams indicate the identity of the proteasome subunits. A subset of the Figure Salso appears in Figure S1E. **(B)** Immunoblotting band intensity across tumor or adjacent tissues were quantified for PSME4 and normalized to actin as a loading control (paired t-test *P = 0.0437). **(C)** Immunoblotting band intensity across tumor or adjacent lung tissues from an independent cohort were quantified for PSME4 and normalized to actin as a loading control (paired t-test *P $\leq$0.01). **(D)** H&E staining of tumor or adjacent tissue from 2 representative patients. **(E)** Immunohistochemistry staining for PSME4 of tumor or adjacent tissue from 3 representative patients.

Figure S5

**Cohort stratification by PSME4 abundance. (A-E)** Quantification of PSME4 in NSCLC subtypes lung adenocarcinoma (LUAD) and squamous cell lung carcinoma (SQCLC) measured by immunohistochemistry staining. Samples are stratified by subtype (A), Smoking level (B), T stage (C), N stage (D) or TNM stage (E). Dots are colored by subtype (LUAD – yellow, SQCLC - red).

Figure S6

**Analyzing proteasome complex composition reveals hybrid PSME4 populations. (A)** The carboxy terminal residue of peptides was used to classify these peptides based on the proteasome activity attributed to their cleavage. The abundance of PSME4 in the samples based on WC proteomics correlated with the chymotryptic-like signature (spearman rho = 0.33). **(B)** Workflow of the size exclusion separation. A549 cells treated with TNFα and IFNγ (TI) or untreated then recombinant PSME4 was added to TI treated A549 lysates. **(C)** Lysates were separated by size using Superose6 gel filtration column and blotted against different proteasome subunits as indicated. **(D)** Lysates of A549 cells treated with TNFα and IFNγ (TI) or untreated used for immunoprecipitation shown in Figure S2E blotted with the indicated antibody.

Figure S7

**Proteasome activity assay calibration with different fluorogenic substrate. (A-C)** Proteasome activity assays using fluorogenic substrates LLE-bNA (caspase, β1;A), nLPnLD-AMC (caspase, β1;B), or LLVY-AMC (chymotryptic, β5;C) for 3.5 hours. Relative fluorescence (RFU) of the substrate is shown across the 3.5 hours of the experiment (left) or at the endpoint (right). Recombinant PSME4 in different ratios, as indicated, was added to A549 lysates (one-way ANOVA; **P <0.01, **** P < 0.0001). **(D-H)** Proteasome activity assays using the nLPnLD-AMC (caspase, β1; D), RLR-AMC (tryptic, β2/ β2i; E), PAL-AMC (chymotryptic, β1i; F), LLVY-AMC (chymotryptic, β5/ β5i; G) or ANW-AMC (chymotryptic, β5i; H) substrate. Relative fluorescence (RFU) of the substrate is shown across the 3.5 hours of the experiment (left) or at the endpoint (right). A549 lysates were treated with TNFα and IFNγ (T+I; red) or untreated (UT; black) and recombinant PSME4 was added to the lysate where indicated (paired T-test; **P <0.01,***P <0.001, **** P < 0.0001) Black squares indicate the portion of the Figure reproduced in Figure 2.

Figure S8

**The cellular immunopeptidome is altered by PSME4 overexpression. (A)** A549 cells transfected with a PSME4-expressing plasmid or empty vector as a control. Cell lysates were blotted for PSME4 and PSMA1-7. β-Actin was blotted as a loading control. **(B)** The mean fold change ratio between overexpression and control peptides is plotted for the 463 peptides differentially presented upon PSME4 overexpression. **(C)** The ratio between the enrichment score for the cellular component groups enriched in peptides increased in expression upon PSME4 overexpression (OE) over control (WT).

Figure S9

**Validation of PSME4 depletion by shRNA. (A)** qPCR of PSME4 in A549 cell line following depletion of PSME4 with shA223 or shA073 compared to shCtrl with and without stimulation with TNFα and IFNγ (T+I) (paired T test **P < 0.01 and *P ≤ 0.05). **(B)** Lysates of A549 cells with PSME4 knockdown (shA223 and shA073) or control (shCtrl) were blotted for PSME4 and β-Actin as a loading control. **(C)** Quantification of PSME4 band intensity in A549 cell line with PSME4 KD (A223 or A073) or shCtrl (Ctrl) across three biological repeats normalized to actin as a loading control and to shCtrl (Welch's corrected T-test *P = 0.0186; ***P = 0.0005).

Figure S10

**Gating strategy for HLA-A, B, C expression on A549 cells.** The main cell population was gated on FSC-A and SSC-A, followed by doublet discrimination by FSC-A vs FSC-H. Live cells were gated as Zombie Aqua dead cell staining negative. Finally, HLA-A, B, C histograms were overlaid.

Figure S11

**Cellular inflammation is increased following PSME4 depletion (A-B)** The 20% of the LUAD cohort with the highest PSME4 expression was compared to the lowest 20%. The difference between the means and the significance of the difference (negative log 10 p value) are plotted for the genes in the cohort (A). Pathways in the reactome or biocarta annotation sets which were found by GSEA to be significantly enriched in the PSME4 high group are listed (B). **(C)** Pairwise Pearson correlation between the proteins identified following WC proteomics of A549 cells with depletion of PSME4 (KD) or control shRNA (ctrl) and stimulation with TNFα and IFNγ (TI) or untreated. (**D**) Normalized enrichment score (NES) from hallmark pathways significantly enriched in cells stimulated with TNFα and IFNγ and expressing either shPSME4 or shCtrl. Color indicates the significance of the enrichment. **(E)** Lysates of H460 cells with PSME4 knockdown (shA223, sh913 and shA073) or control (shCtrl) were blotted for PSME4 and β-Actin as a loading control. **(F)** qPCR of genes downstream to IFNγ activation (TNFα, IFIT1, A20 and ISG15) following stimulation with TNFα and IFNγ. H460 cells expressing shPSME4 (shA223) are compared to those expressing shCtrl (paired T test ****P < 0.0001, **P < 0.01 and *P ≤ 0.05).

Figure S12

**Categorizing PSME4 depletion in KP1.9 cells. (A)** Lysates of KP1.9 cells with PSME4 knockdown (shK369 and shK428,shK569 or shK813) or control (shRFP) were blotted for PSME4 and β-Actin as a loading control. **(C)** Quantification of PSME4 band intensity in KP1.9 cell line with PSME4 KD or Ctrl (shRFP) across three biological repeats normalized to actin as a loading control and to shRFP .**(C)** KP1.9 cells and healthy mouse lung tissue were blotted for PSME4. β-Actin was blotted as a loading control. **(D)** qPCR shows the expression of PSME4 in KP1.9 transfected with a PSME4 overexpression plasmid. **(E)** KP1.9 transfected with a PSME4 overexpression plasmid (+) or GFP (-) as a control. Cell lysates were blotted for PSME4 or PSMA1-7. β-Actin was blotted as a loading control. **(F)** Band intensities of (E) were quantified for PSME4 and normalized to actin as a loading control (paired t-test *P = 0.0331). **(A)** Lysates of KP1.9 cells with mPSME4 knockdown (shK369, shK428, shK569 and shK813) or control (shRFP) were blotted for PSME4 and β-Actin as a loading control. **(B)** Quantification of PSME4 band intensity of (A) across three biological repeats were normalized to actin as a loading control and to shCtrl. **(G)** Standard curves for quantifying levels of IL-17a or IL-6 in tumor secretomes. **(H-I)** The level of cytokine secretion was examined for PSME4-deficient KP1.9 cells (H) or KP1.9 controls (I) inhibited with ONX-0914 or left untreated. The log2 transformed ratio between the two conditions is plotted against the negative log10 transformed p value.

Figure S13

**Tumor burden is increased following injection of PSME4-overexpressing KP1.9 cells. (A)** Weights of C57/B6 mice bearing orthotopic wild type (KP1.9), PSME4-overexpressing (OE), or PSME4-deficient (KD) KP1.9 tumors (n = 7-8 per group). Weights are normalized to the starting weight of each mouse. Overexpression shows significantly decreased weight as a proxy for increased disease severity (matched 2-way ANOVA *p = 0.0322). **(B)** Mice bearing PSME4 overexpressing (OE) tumors showed significantly increased tumor burden compared to the control KP1.9 tumors. **(C-D)** Two biological repeats of H&E staining of lungs from mice bearing KP1.9, KP1.9$^{PSME4\ OE}$, or KP1.9$^{PSME4\ KD}$ tumors presented in Fig. 3 **(E)** Measurement of the cell growth of KP1.9 (WT), KP1.9 expressing shCtrl or shPSME4 (PSME4 KD) for 3 days. **(F)** Measurement of the cell growth of KP1.9 (WT), KP1.9 expressing GFP overexpression (GFP Ctrl) or PSME4 overexpression (OE) for 3 days.

Figure S14

**Classification of populations in single cell RNA sequencing (A)** Stacked bar plot of the number of cells per cell type identified **(B)** Key markers used to describe cluster identity and link it to cell type.

Figure S15

**Classification of mreg and plasmacytoid DC inflammatory profile (A-D)** GO-term enrichment (A, C) and heatmap (B, D) of key markers from the plasmacytoid DC (A, B) or mature DCs enriched in immunoregulatory molecules (mregDCs) (C, D) population.

Figure S16

**Analysis of clonal expansion in T cell clusters (A)** The number of expanded clones (i.e. having non-unique TCR) in individual mice bearing KP1.9 (Ctrl) or KP1.9$^{PSME4\ KD}$ (KD) tumors. (B) Bar graph showing number of expanded cells in all cell populations from mice injected with KP1.9 (n = 3) or PSEM4 knock down KP1.9 (PSEM4KD) (n = 3) cells.

Figure S17

**Cytotoxic CD8 and T reg marker genes. (A-B)** Heatmaps of marker signatures of T regulatory cells (A) or cytotoxic T cells (B).

Figure S18

**Cytotoxic CD8 and T reg signature genes. (A-B)** Bar plots showing pseudobulk tpm expression of *Foxp3* (A) or *Gzmk* (B) across the clusters. **(C-D)** A CD8 T cell (C) or Treg (D) signature score for each of the clusters.

Figure S19

**Categorization of proliferation and infiltrate populations in KP1.9 model. (A-D)** The percentages of the CD45 positive population in the lung (A, B) or spleen (C, D) from the mice bearing KP1.9, KP1.9$^{PSME4\ OE}$, or KP1.9$^{PSME4\ KD}$ tumors or mice not injected (non-inj) with tumor cells that are CD4 (A, C) or CD8 (B, D) positive. One way ANOVA with post-hoc TUKEY analysis was used to compare populations (*$P \leq 0.05$).

Figure S20

**Gating strategy for lung tissues. (A)** Histograms of PD-1 fluorescence intensity from mice bearing KP1.9 (grey, n=4) or KP1.9$^{PSME4\ OE}$ (pink, n=3)**. (B)** The lymphocyte population was first gated by FSC-A and SSC-A and doublets discriminated by FSA-A vs FSC-H. Live cells were determined by Zombie Aqua negativity and the lymphocytes by CD45 positivity. T cells were gated as CD3e positive, followed by CD4 and CD8 to discriminate T helper and cytotoxic T cells. PD-1 expression was gated by an FMO samples on CD4 and CD8 cells. The activation status of CD4 and CD8 T cells was examined by CD62L vs CD44 staining as indicated. **(C)** The percent of different subsets of CD8-positive lymphocytes (naïve, early activated, effector memory [EM] or central memory [CM], which are CD62L and/or CD44 positive, in the lung of mice bearing KP1.9$^{PSME4\ OE}$, KP1.9$^{PSME4\ KD}$ or KP1.9 tumors compared to mice not bearing tumors (non-inj). The portion of the figure marked in the box is reproduced in Figure 4G.

Figure S21

**PSME4 modulation of tumor growth is immune-mediated.** The body weight of wild-type or immunocompromised RAG -/- mice injected with KP1.9 lung tumors expressing either shPSME4 or Ctrl shRNA. The weight of each mouse is normalized to the starting weight at day 1. 'X' indicates death.

Figure S22

**Characterization of splenic lymphocytes. (A)** The main cell population was gated by FSC-A vs SSC-A followed by FSC-A vs FSC-H (two populations are visible as the gate contains lymphocytes and KP1.9 tumor cells). KP1.9 cells are gated as CFSE-positive and SSC-A low-high; the lymphocytes as CFSE-negative and low SSC-A. Finally, the dead cell population is gated versus PI-A (propidium iodide) to determine live (PI negative) and dead (PI positive) populations. **(B-D)** Standard curves for quantifying levels of IFNγ (B), I7a (C) or IL-22 (D) in secretomes.

Figure S23

**The immunoproteasome characterizes the hot tumor state.** T-cell inflammation signature was calculated for the LUAD samples in the CPTAC proteomics cohort (n = 213 samples). Samples were stratified based on the distribution of the subunit in the adjacent tissue where those that are more abundant than the top 5% of the adjacent tissues are considered high abundance (Wilcoxon *$P \leq 0.05$, *** $P \leq 0.001$, ****$P \leq 0.0001$).

Figure S24

**PSME4 and PSMB10 but not PSME3 correlate with responsiveness to ICI. (A)** The percentage of tumors in each cancer type with above average levels of PSME4 (PSME4-high Tumors). The colors indicate the mean expression of PSME4 normalized to the core proteasome subunits **(B-C)** The expression of PSME4 (B) or PSMB10 (C) normalized to the core proteasome were used to stratify samples into low and high expressers (upper and lower 50% of samples). The percentage of responders and non-responders to ICI was counted for each group per cancer type and across the ICI1000+ cohort ($\chi^2$ test, B**** P = 0.000096, D*** P = 0018).

Figure S25

**PSME4 does not correlate with other important ICI biomarkers. (A)** The spearman correlation between different biomarkers from the ICI1000+ study. Circle size shows absolute correlation coefficient and asterisks indicate significance. **(B)** The odds ratio of association with survival for the biomarker indicated for each of the cohorts within the ICI1000+ study.

Figure S26

**Characterization of tumor inflammation through EVOC system. (A-B)** Immunohistochemistry and H&E from tumor sections with response-signature (A) and non-response-signature (B) EVOC section. **(C-F)** Immunohistochemistry of resected tumors matching EVOCs with response (C, D) and non-response (E, F) signatures as indicated for PSME4 (C, E) and PSMB10 (D, F) for 6 representative patients. **(G-H)** qPCR of PSME4 (G) or PSMB10 (H) expression in EVOC sections with response and non-response signatures.
